# NMN Works in HFD-Induced T2DM by Interesting Effects in Adipose Tissue, and Not by Mitochondrial Biogenesis

**DOI:** 10.1101/2023.10.27.564318

**Authors:** Roua Gabriela Popescu, Anca Dinischiotu, Teodoru Soare, Ene Vlase, George Cătălin Marinescu

## Abstract

Nicotinamide mononucleotide (NMN) has emerged as a promising therapeutic intervention for age-related disorders, including Type 2 Diabetes.

In this study, we investigated the effects of NMN treatment on glucose uptake and its underlying mechanisms in various tissue and cell lines. Through a comprehensive proteomic analysis, we uncovered a series of distinct organ-specific effects that contribute to the observed improvements in glucose metabolism.

Notably, we observed the upregulation of thermogenic UCP1, promoting enhanced glucose utilization in muscle tissue. Additionally, liver and muscle cells displayed a unique response, characterized by spliceosome down-regulation and concurrent upregulation of chaperones, proteasomes, and ribosomes, leading to a mildly impaired and energy-inefficient protein synthesis machinery. Adipose tissue exhibited increased protein synthesis and degradation, fatty acid degradation, Lysosome and mTOR cell proliferation signalling up-regulation, while showing a surprising repressive effect on mitochondrial biogenesis.

Furthermore, our findings revealed a remarkable metabolic rewiring in the brain, involving increased production of ketone bodies, down-regulation of mitochondrial OXPHOS components and the TCA cycle, and the induction of known fasting-associated effects.

Collectively, our data elucidate the multifaceted nature of NMN action, highlighting its organ-specific effects and their role in modulating glucose metabolism. These findings deepen our understanding of NMN’s therapeutic potential and pave the way for novel strategies in managing metabolic disorders.

## 1. Introduction

The incidence of type 2 diabetes mellitus (T2DM) has been steadily rising on a global scale. The International Diabetes Federation (IDF) estimates that the number of adults living with diabetes has quadrupled since 2000. In 2021, it was reported that approximately 537 million adults had diabetes, and this number is projected to rise to 783 million by 2045 if current trends continue [1]. Factors contributing to the increased incidence include: lack of physical exercise, unhealthy dietary patterns, obesity, population aging, and urbanization [2–4]. The rise in T2DM cases presents a major public health challenge, necessitating concerted efforts in prevention, early detection, and effective management strategies to mitigate its impact on individuals and healthcare systems worldwide. The treatment of diabetes has become an increasingly expensive burden on global healthcare systems, with the worldwide expenditure reaching 966 billion USD in 2021, according to the International Diabetes Federation [1], representing a significant increase by 32.8% compared to the 727 billion USD spent in 2017 [5], underscoring the urgent need for innovative treatments and interventions.

Glucose metabolism is a fundamental process vital for energy homeostasis in mammals, orchestrated through a complex interplay among various organs and tissues. Among these, the liver, muscle, adipose tissue, and brain stand as the primary consuming and target tissues for glucose utilization [6]. The liver regulates blood glucose levels, while muscle tissue plays a significant role in glucose uptake and energy expenditure [7,8]. Adipose tissue, traditionally viewed as an energy reservoir, is increasingly recognized for its active involvement in glucose metabolism [9]. The brain, a glucose-dependent organ, demands a continuous and tightly regulated supply of glucose to support its functions [10]. Understanding the intricate correlations between these vital organs is crucial for unraveling the complexities of glucose homeostasis, metabolic health, and related disorders, like T2DM.

Previous studies have reported that β-nicotinamide mononucleotide (NMN) as a precursor of nicotinamide adenine dinucleotide (NAD^+^), ameliorate several age-related diseases, including T2DM, characterized by high blood glucose levels and insulin resistance [11–13]. Beside increasing the intracellular levels of NAD^+^, which plays a vital role in regulating cellular redox state, NMN acts through several biological and chemical processes, most notably being DNA repair, energy metabolism and stress response [14]. Previously, it was proposed that raised NAD^+^ levels activate SIRT1, leading to mitochondrial biogenesis by upregulating the expression of transcription factor A mitochondrial (TFAM) and peroxisome proliferator-activated receptor gamma coactivator 1-alpha (PGC-1α), as well as increased DNA repair and AMP-activated protein kinase (AMPK) activation [12,15,16]. The efficiency of NMN in the treatment of diet and age-induced diabetes in mice was previously reported [11,17].

However, human trials results are disappointing [18], and previously proposed NMN induced mitochondrial biogenesis idea is doubtful. Previous mechanistic approaches failed to provide a significant response to the question: If NMN works in T2DM, where is glucose going? The main flaws identified in these studies were most probably unreliable transcriptomics data, unreliable western blots (for example choosing actin as reference protein, which our data shows that is dramatically affected by NMN treatment (**Figure S20**).

In the metabolic and signalling pathways, enzymes (that are proteins), play the effector role in chemical reactions, contributing to the cellular maintenance and allowing the rapid adaptation to the environmental changes. Quantifying their expression levels might provide novel insights into the intricate connections between diverse metabolic and signalling pathways [19]. Considering these, our study represents an effort to see the big picture, a proteome-centric approach to elucidate the underlying mechanisms and the therapeutic potential of NMN in the treatment of T2DM through the analysis of its proteome level effects, including energy metabolism, in an *in vivo* diabetic mouse model, as well as in two *in vitro* models (C2C12 derived myotubes and HepG2 cells, respectively). This investigation used state-of-the-art data-independent acquisition (DIA) technologies in mass spectrometry, specifically SWATH-MS, as well as label-free and library-free peptide mapping using neural networks and interference correction to achieve deep proteome coverage. An integrated map of the molecular mechanisms underlying the NMN effects in T2DM treatment is provided.

## 2. Results

The differential expression changes in the proteome of an *in vivo* T2DM mouse model as well as in *in vitro* models of hyperglycemic muscle (myotubes derived from C2C12 cells) and liver (HepG2 cells) have been explored. A thorough library-free and label-free proteomic analysis has been conducted. The spectra library was created from actual DIA data coupled with fasta files for the *Mus musculus* and *Homo sapiens* proteomes, respectively. Overall, we have quantified a total of 1728 unique proteins and 20857 precursors in mouse muscle tissue at a false discovery rate (FDR) < 1%. In mouse liver, we have identified 3097 unique proteins and 31501 precursors, while in mouse adipose tissue, we have found 3217 unique proteins and 35516 precursors. Furthermore, in mouse brain, we have quantified 3814 unique proteins and 43320 precursors, and in myotubes derived from C2C12 cells, we have identified 3855 unique proteins and 35481 precursors. Finally, in HepG2 cells, we have detected 3541 unique proteins and 43565 precursors across five biological replicates, each multiplied by three technical replicates. The effect of the NMN treatment was evaluated by highlighting significant changes in protein expression levels. Limma test was performed in PolySTest tool [20], which found significantly changed (FDR < 5%) 252 proteins in mouse muscle tissue, 337 proteins in mouse liver, 411 proteins in mouse adipose tissue, 120 proteins in mouse brain, 1200 proteins in C2C12 myotubes, and 984 proteins in HepG2 liver cells. All significantly changed proteins are available in Supplemental Information (**Tables S1-S6**), and have been used as input for pathway enrichment analysis.

The PathfindR module in R statistical software package was used to construct the protein-protein interaction network (PIN) for *Mus musculus* and *Homo sapiens* from the STRING proteins database. Using PathfindR (version 1.6.4), an enrichment chart containing the top 20 affected pathways and a Term gene graph containing the top 10 pathways (**Figures 3,5,7 and 8**) were generated. The KEGG pathways [21–23] were generated by the Pathview R module (version 1.38.0) and show the relative expression of each protein from the NMN treated condition compared to the untreated condition.

### 2.1. NMN significantly improved glucose uptake from bloodstream in HFD mice, however not through insulin resistance effects in muscle or liver

No significant differences in the serum total cholesterol and triglyceride levels between the HFD group and the HFD+NMN group have been noticed (**Figure 1**B). The HFD+NMN group had higher levels of total cholesterol (4.954 mmol /L) compared to the HFD group (3.848 mmol /L). Similarly, the triglyceride levels were slightly higher in the HFD+NMN group (1.378 mmol /L) compared to the HFD group (1.364 mmol /L), however, the differences were not statistically significant (*p* =0.14) respective (*p* =0.11). The results of the IPGTT test indicate differences in glucose absorption patterns between the HFD group and the HFD+NMN treated group (**Figure 1**C). After NMN treatment (**Figure 1**D), the HFD+NMN group had significant lower glucose levels compared to the HFD group after 15, 30, 60, and 120-minutes after glucose intraperitoneal injection. Surprisingly, GLUT4 glucose receptor was significantly down regulated in treated HFD mice muscle tissue, as well as GS (Glycogen Synthase).

**Figure 1.**
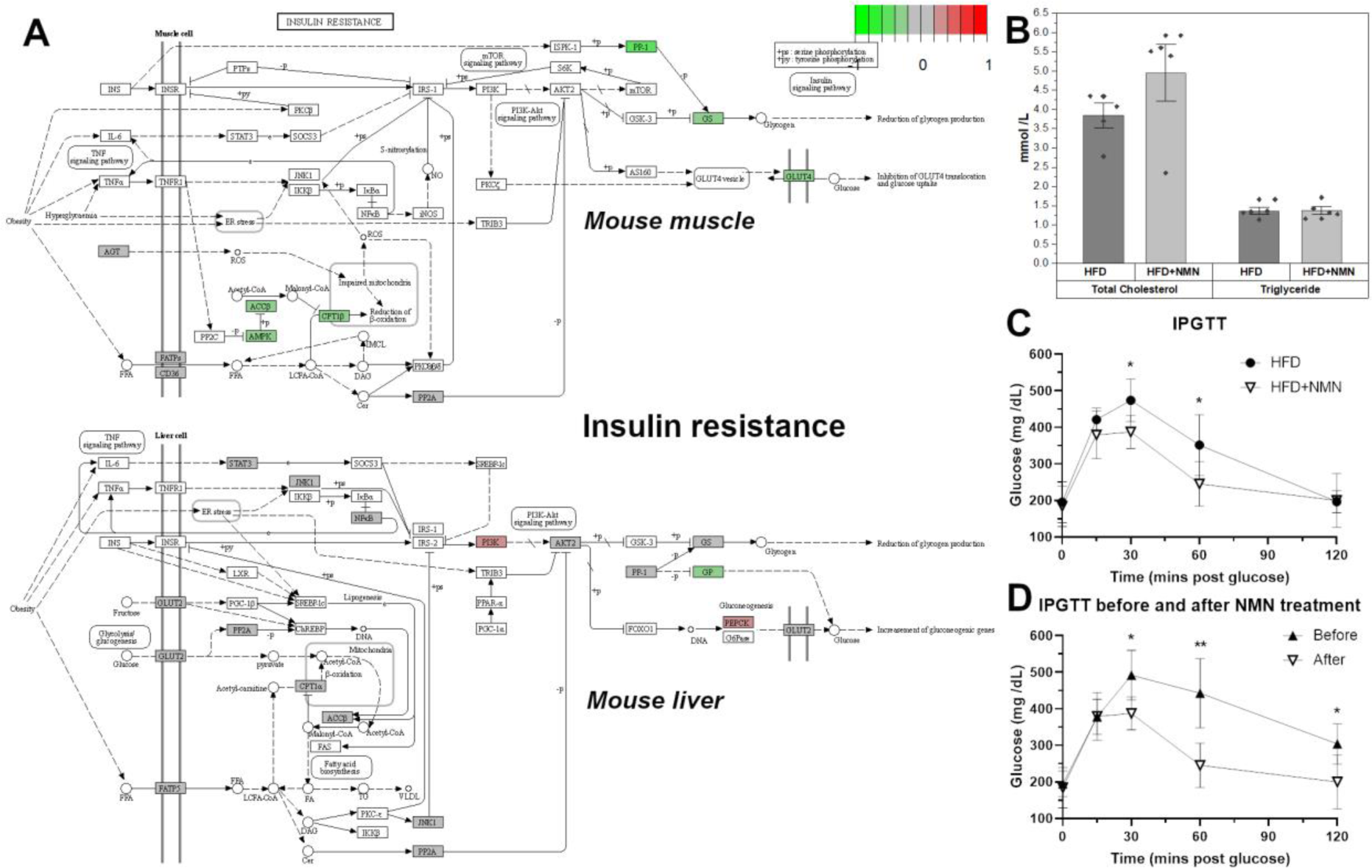
C57BL/6J mice develop severe glucose uptake deficiency on high-fat diet (HFD). (A) NMN treatment effects on the KEGG insulin resistance pathway in mouse muscle and liver. The colour of the boxes represents the log2 fold change of the protein abundances, represented for HFD+NMN group versus HFD group comparison. Red: up-regulated; green: down-regulated; grey: no significant expression change. GS and GLUT4 are significantly down-regulated in treated mice muscle tissue. (B) Serum total cholesterol and triglycerides in HFD respective HFD+NMN mice,7 days after treatment. (C) Intraperitoneal Glucose Tolerance Test (IPGTT) in HFD and HFD+NMN treated mice after 7 days. (D) IPGTT in HFD+NMN treated mice before and after NMN treatment. The data are illustrated as average values of the groups (n = 5) ± standard deviation of the mean (STDEV) and statistical significance between HFD and HFD+NMN groups. \**p* <0.05; \*\**p* <0.01.

### 2.2. Protein synthesis is mildly impaired by NMN through spliceosome down-regulation in HFD mouse liver

In mouse liver samples 103 proteins (Figure 2) showed a significant difference in expression level between the HFD and HFD+NMN treated groups. Data were filtered for log_2_FC above 0.3 or below −0.3 and FDR (Limma) < 0.05. The log_2_FC ranged from −1.402 on the under-expression side respectively 1.618 on the over expression side. Among these proteins, 19 proteins were upregulated and 17 proteins were downregulated. The top five upregulated proteins were Acnat2 (log_2_FC=1.62, FDR=0.024), Cyp2b10 (log_2_FC=1.24, FDR=0.003), Cyp2b9 (log_2_FC=1.03, FDR=0.023), Atxn7l3b (log_2_FC=0.73, FDR=0.016) and Creld1 (log_2_FC=0.57, FDR=0.002). The most significantly downregulated protein was Ces3b (carboxylesterase 3B) with a log_2_FC of −0.54 and an FDR < 0.005. Other proteins with significantly affected were Atp2a1 (log_2_FC=-1.40, FDR=0.005) Apex1 (log_2_FC=-0.30, FDR=0.005), Clic4 (log_2_FC=-0.31, FDR=0.0002), Cml5 (log_2_FC=-0.54, FDR=0.029), Ctsf (log_2_FC=-0.34, FDR=0.023) and Ctsz (log_2_FC=-0.31, FDR=0.023) that have been downregulated, and Aldh1a1 (log_2_FC=0.41, FDR=0.0001), Akr1d1 (log_2_FC=0.44, FDR=0.001), and Cyp2b10 (log_2_FC=1.23, FDR=0.002) have been upregulated, respectively.

**Figure 2.**
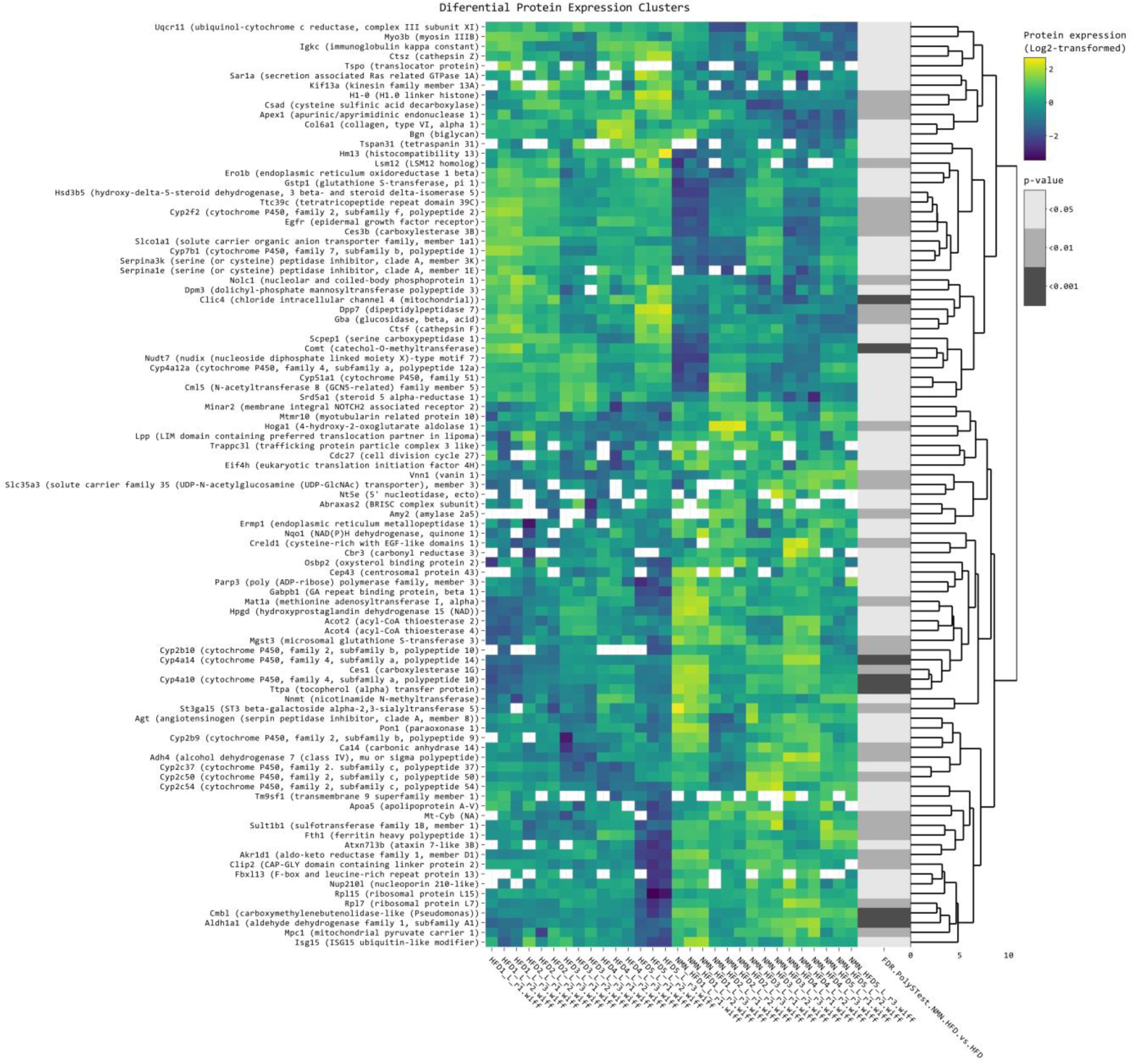
Clustered heatmap of the differentially expressed proteins in mouse liver. Clustered heatmap of the 103 differentially expressed proteins in mouse liver tissue, filtered with log_2_FC threshold set to exclude the interval −0.3, 0.3, FDR <0.05. Yellow color represents upregulation, while blue represents downregulation. From left to right, expression values (log_2_ transformed) for replicates (5 biological x 3 technical) are shown for the HFD group and for the HFD + NMN treated group, followed by significance values of the comparison to HFD group.

Enrichment chart (**Figure 3**A) and Term gene graph (**Figure 3**B) analysis generated in PathfindR from mouse liver proteomics data revealed several biological pathways that were significantly affected. The top affected pathways were steroid hormone biosynthesis, retinol metabolism, ribosome, arachidonic acid metabolism, metabolism of xenobiotics by cytochrome P450, spliceosome, fatty acid degradation, and inflammatory mediator regulation of TRP channels. The steroid hormone biosynthesis pathway (**Figure S9**) was significantly altered with upregulated proteins, including Akr1d1, Cyp2b10, Cyp2b9, Cyp2c29, Cyp2c37, Cyp2c50, Cyp2c54, and Cyp2e1. Similarly, the retinol metabolism pathway was affected by upregulated proteins such as Adh4, Aldh1a1, Cyp2b10, Cyp2b9, Cyp2c29, Cyp2c37, Cyp2c50, Cyp2c54, and Cyp4a10. On the other hand, the ribosome pathway (**Figure S10**) showed upregulated proteins such as Mrps5, Rps9, Rps24, Rpl7, and Rpl15, while proteins Rps6, Rps8, Rps15, Rps21, Rps27l, Rpl17, and Rpl31 were downregulated. The arachidonic acid metabolism pathway (**Figure S11**) showed upregulated proteins such as Cbr3, Cyp2e1, Cyp4a10, Cyp4a14, Cyp2b10, Cyp2b9, Cyp2c29, Cyp2c37, and Cyp2c54, and downregulated proteins, such as: Lta4h and Cyp4a12a. Similarly, the metabolism of xenobiotics by cytochrome P450 pathway was affected with upregulated proteins such as Gsta3, Gstt1, Gstt2, Mgst3, Ephx1, Cyp2e1, Cbr3, and Adh4, while proteins Gstp1 and Cyp2f2 were downregulated.

**Figure 3.**
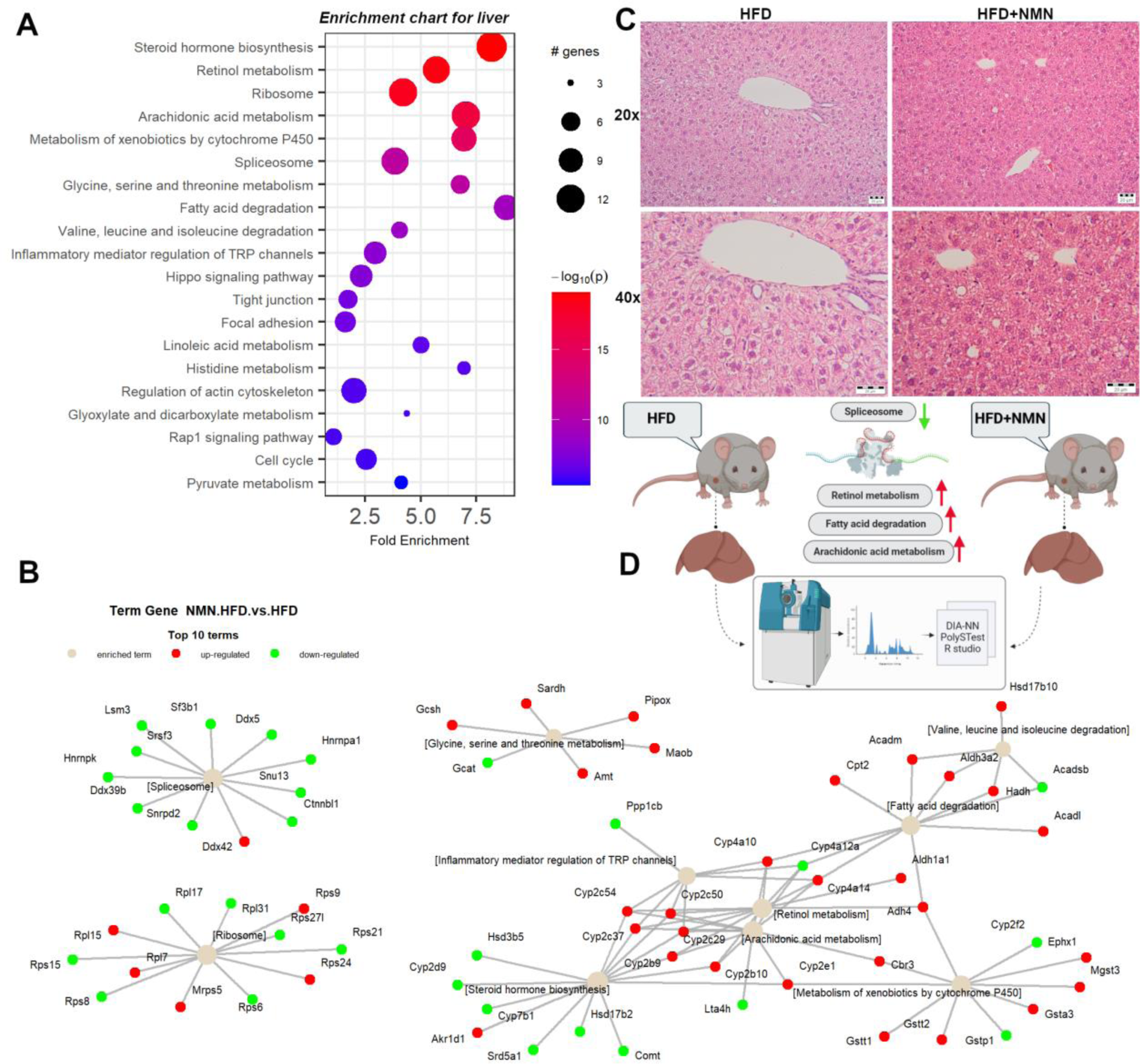
Integrated proteomics data analysis of NMN treated HFD mouse liver. (A) Enrichment chart for top 20 KEGG Pathways sorted by lowest *p* value. (B) Term-gene graph for top 10 terms. (C) Representative images of hematoxylin and eosin staining for mouse liver tissue. Scale bars, 20 μm. (D) Experiment summary (BioRender).

Other upregulated pathways were: inflammatory mediator regulation of TRP channels, hippo signalling pathway, glycine, serine and threonine metabolism, fatty acid degradation, valine, leucine and isoleucine degradation, linoleic acid metabolism, histidine metabolism, regulation of actin cytoskeleton, glyoxylate and dicarboxylate metabolism, and pyruvate metabolism. Downregulated proteins were observed in tight junction, focal adhesion, Rap1 signalling pathway, and cell cycle pathways.

### 2.3. Thermogenesis pathway is up-regulated by NMN in muscle tissue

Skeletal muscle is a highly specialized tissue that is essential for movement and controls the body’s whole glucose metabolism. It is responsible for 75–80% of glucose uptake during hyperinsulinemia, followed by the adipose tissue and liver. The complete list of identified proteins including log_2_ fold change (log_2_FC), FDR, expression values for each individual and technical replicate is provided in the Supplemental Information.

Clustered heatmap of the 119 differentially expressed proteins in all samples (5 biological x 3 technical replicates in each condition /diet) (**Figure 4**) shows the proteins grouped by several similar expression patterns. The proteins in the first small cluster are grouped in three subclusters. Among these, Gnmt, Aldob, and Hmgcs2 had the highest log_2_FC of 1.39, 1.21, and 1.06, respectively. The second subcluster had six differentially expressed proteins, with Thbs4 having the highest log_2_FC of 0.76. The third subcluster had four differentially expressed genes, with Ucp1 having the highest log_2_FC of 1.64 among all the identified proteins.

**Figure 4.**
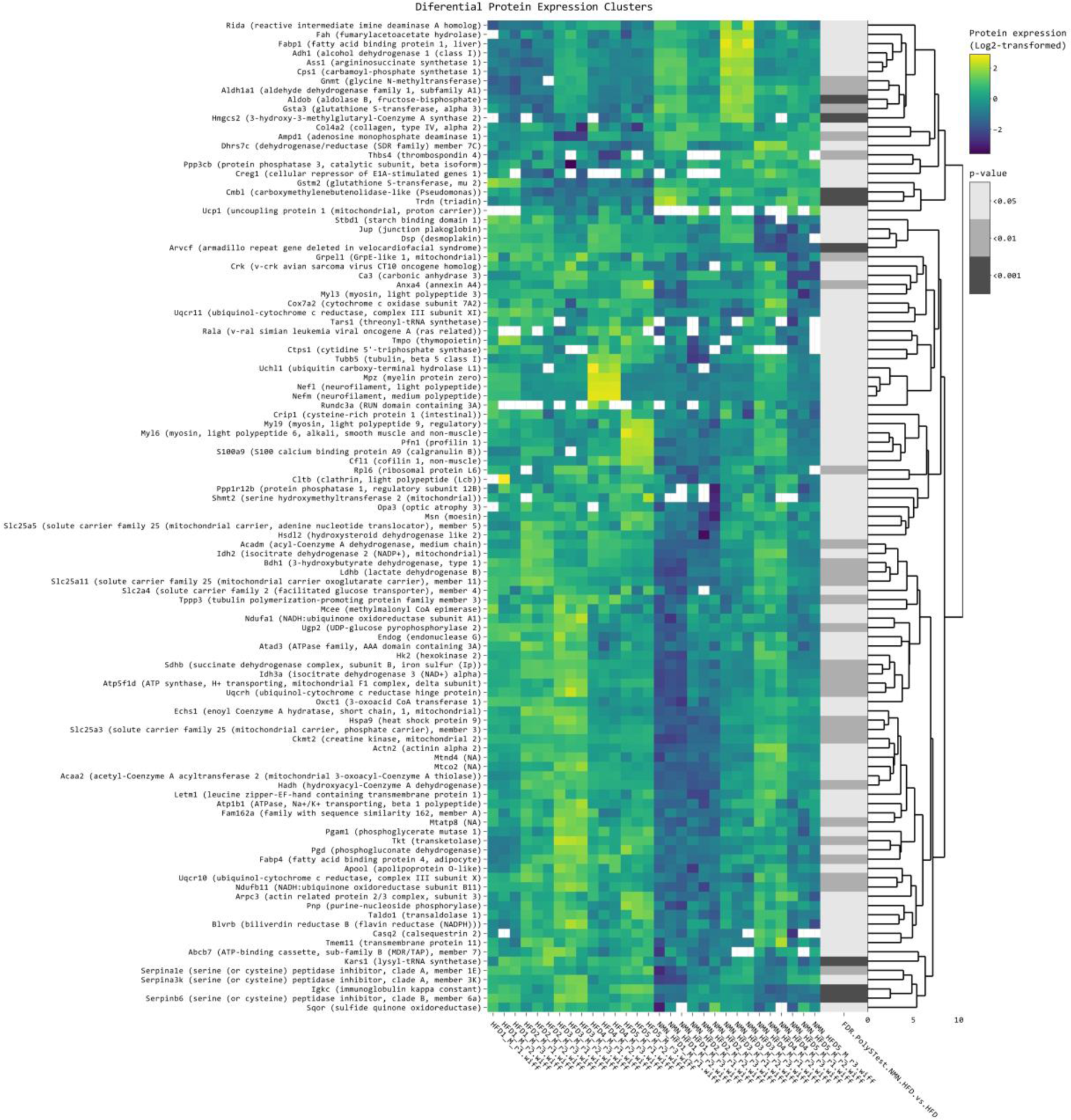
Clustered heatmap of the differentially expressed proteins in mouse skeletal muscle tissue. Clustered heatmap of the 119 differentially expressed proteins in mouse muscle tissue, filtered with log_2_FC threshold set to exclude the interval −0.3, 0.3, allowing FDR < 0.05. Yellow color represents upregulation, while blue represents downregulation in HFD+NMN treated group compared to HFD group. From left to right, expression values (log_2_ transformed) for replicates (5 biological x 3 technical) are shown for the HFD group and for the HFD + NMN treated group, followed by significance values of the comparison to HFD group.

Analysis of the second and the larger cluster revealed significant changes in the expression of 38 proteins, with log_2_FC ranging from −1.8576 to −0.3068 and FDR values ranging from 0.00098 to 0.04585. The most significantly downregulated protein expression was Arvcf (log_2_FC = −1.86), followed by S100a9 (log_2_FC = −1.64), Mpz (log_2_FC = −1.58), and Nefl (log_2_FC = −1.40). We also observed downregulation of several other proteins involved in muscle function, including Myl3 (log_2_FC = −1.44), Myl9 (log_2_FC = −0.53), Myl6 (log_2_FC = −0.40), and Tubb5 (log_2_FC = −0.42). Moreover, we observed downregulation of genes involved in mitochondrial function, including Cox7a2 (log_2_FC = −0.32), Uqcr11 (log_2_FC = −0.52), and Slc25a5 (log_2_FC = −0.31).

Enrichment pathway analysis (**Figure 5**A, C) shows downregulated proteins involved in mitochondrial functions, including oxidative phosphorylation and TCA Cycle. We observed downregulation of proteins involved in glycolysis/gluconeogenesis, glycine/serine/threonine metabolism, valine/leucine/isoleucine degradation, metabolism of xenobiotics by cytochrome P450, regulation of actin cytoskeleton, focal adhesion, fatty acid degradation, and cysteine and methionine metabolism. Additionally, some pathways showed both upregulated and downregulated proteins, such as glycolysis/gluconeogenesis (**Figure S15**), glycine/serine/threonine metabolism (**Figure S16**), and pyruvate metabolism (**Figure S17**). Moreover, we identified specific proteins that were significantly upregulated or downregulated in certain pathways, such as Aldob, Adh1, and Aldh7a1 in glycine, serine and threonine metabolism and pentose phosphate pathway, Ucp1 in thermogenesis; and Aldh7a1 and Hmgcs2 in valine, leucine and isoleucine degradation (**Figure S18**). Notably, metabolism of xenobiotics by cytochrome P450 was significantly up-regulated in NMN treated mice muscle tissue (**Figure 5**C).

**Figure 5.**
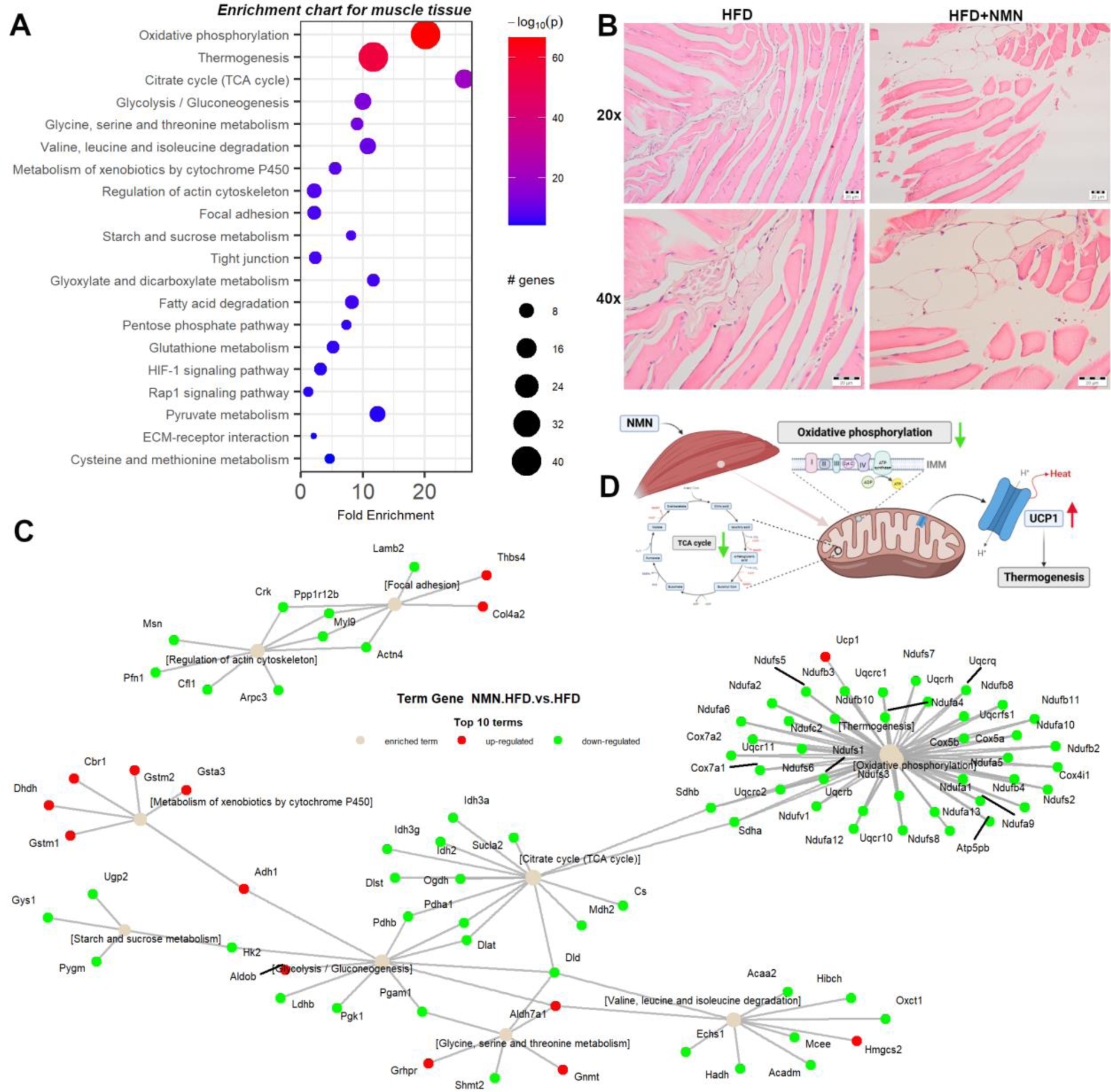
Integrated proteomics data analysis of mouse muscle tissue. (A) Enrichment chart for top 20 KEGG Pathways sorted by lowest *p* value in mouse muscle tissue. (B) Representative images of hematoxylin and eosin staining and semiquantitative analysis from mouse muscle tissue. Scale bars, 20 μm. (C) Term-gene graph for top 10 terms in mouse liver. (D) Experiment summary (BioRender).

### 2.4. NMN stimulates adipose cells proliferation by up-regulating mTOR pathway

The heatmap analysis of mouse adipose tissue (**Figure 6**) revealed significant changes in a cluster formed by 104 proteins. Among the significantly upregulated proteins, the most highly induced were Aoc1 (amine oxidase, copper-containing 1) with a log_2_FC of 4.85 and FDR < 0.03, followed by Epx (eosinophil peroxidase) with a log_2_FC of 2.16 and FDR < 0.01, and Aldob (aldolase B, fructose-bisphosphate) with a log_2_FC of 1.40 and FDR < 0.01. The significantly downregulated proteins included Fut11 (fucosyltransferase 11) with a log_2_FC of −1.04 and FDR < 0.002, followed by H1-5 (H1.5 linker histone, cluster member) with a log_2_FC of −0.50 and FDR < 0.04, and Gnmt (glycine N-methyltransferase) with a log_2_FC of 1.15 and FDR < 0.01.

**Figure 6.**
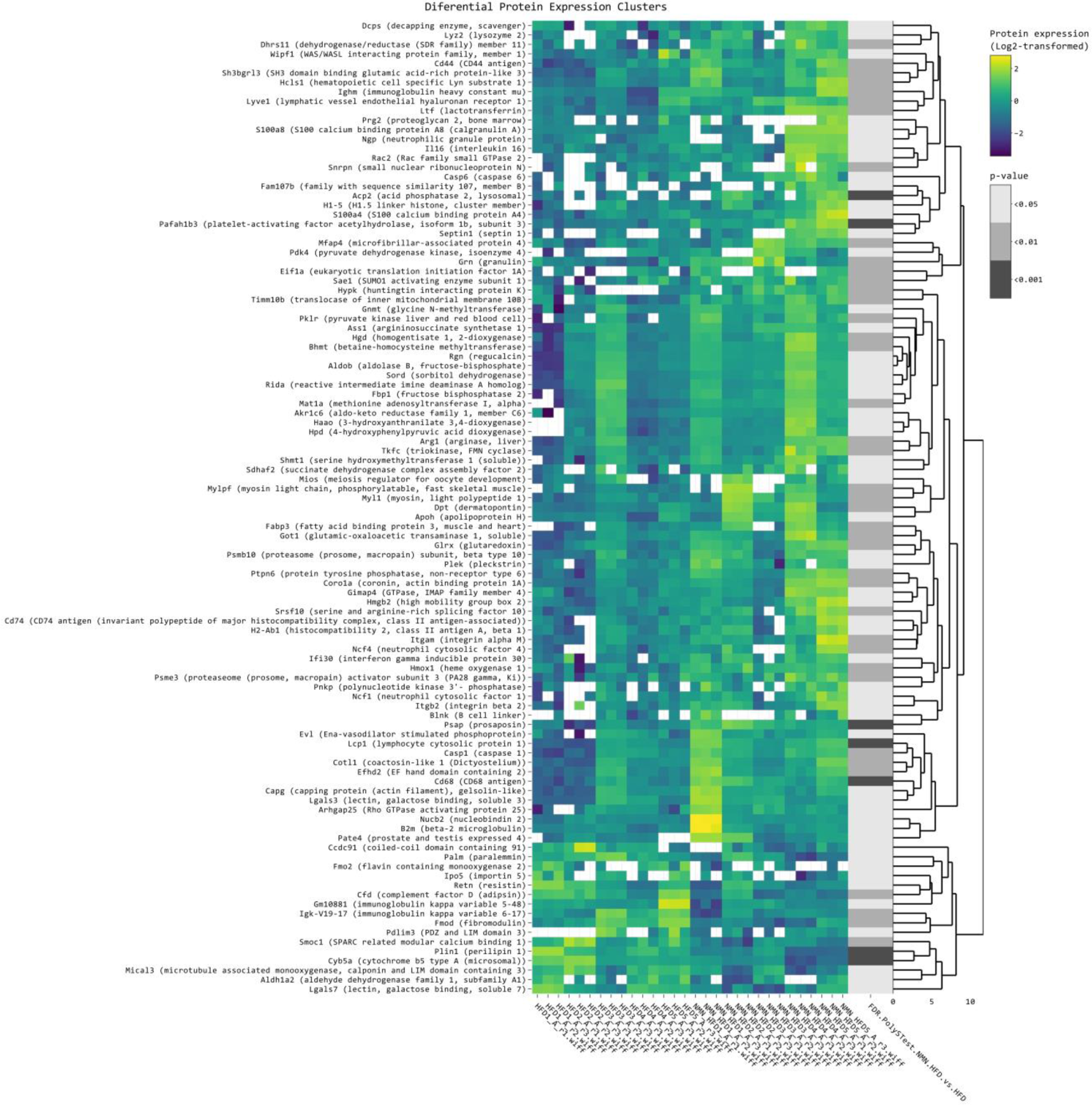
Clustered heatmap of the differentially expressed proteins in mouse adipose tissue. Clustered heatmap of the 119 differentially expressed proteins, filtered with log_2_FC threshold set to exclude the interval −0.5, 0.5 and 0.05 FDR threshold. Yellow color represents upregulation, while blue represents downregulation in NMN treated versus HFD group. From left to right, expression values (log_2_ transformed) for replicates (5 biological x 3 technical) are shown for the HFD group and for the HFD + NMN treated group, followed by significance values of the comparison to HFD group.

In the second and the smallest cluster, sixteen proteins were found to have significantly altered expression levels (FDR < 0.05). Of these, 15 proteins showed a decrease in expression in response to NMN treatment, with log_2_FC ranging from −0.973 to −0.503. One of the proteins significantly repressed was resistin, with a log_2_FC of −0.78 and an FDR < 0.045. Also, other proteins that showed significant changes in expression levels were: Cfd (complement factor D), Gm10881 (immunoglobulin kappa variable 5-48), Igk-V19-17 (immunoglobulin kappa variable 6-17), and Cyb5a (cytochrome b5 type A (microsomal)). These proteins are involved in various physiological processes such as immune response, lipid metabolism, and oxidative stress response (**Figure 7**A). The upregulation of Aoc1, Epx, and Aldob suggests increased oxidative stress and lipid metabolism, whereas the downregulation of Fut11, H1-5, Gnmt, Cltb and Hspa2 indicates altered cellular processes such as immune response and DNA packing (**Figure 7**B). Moreover, enrichment chart (**Figure 7**A) and Term graph (**Figure 7**B) revealed significant changes in expression level of proteins involved in various biological processes and pathways: the spliceosome pathway (**Figure S19**) showed upregulation of Snrpf, Ddx5, Tcerg1, Sf3b1, Sf3b3, Sf3b6, Lsm6, Prpf31, Snu13, Prpf19, Snw1, Bud31, Pcbp1, Srsf2, Srsf3, Srsf6, Srsf7, and Srsf10, while Hspa2 was downregulated.

**Figure 7.**
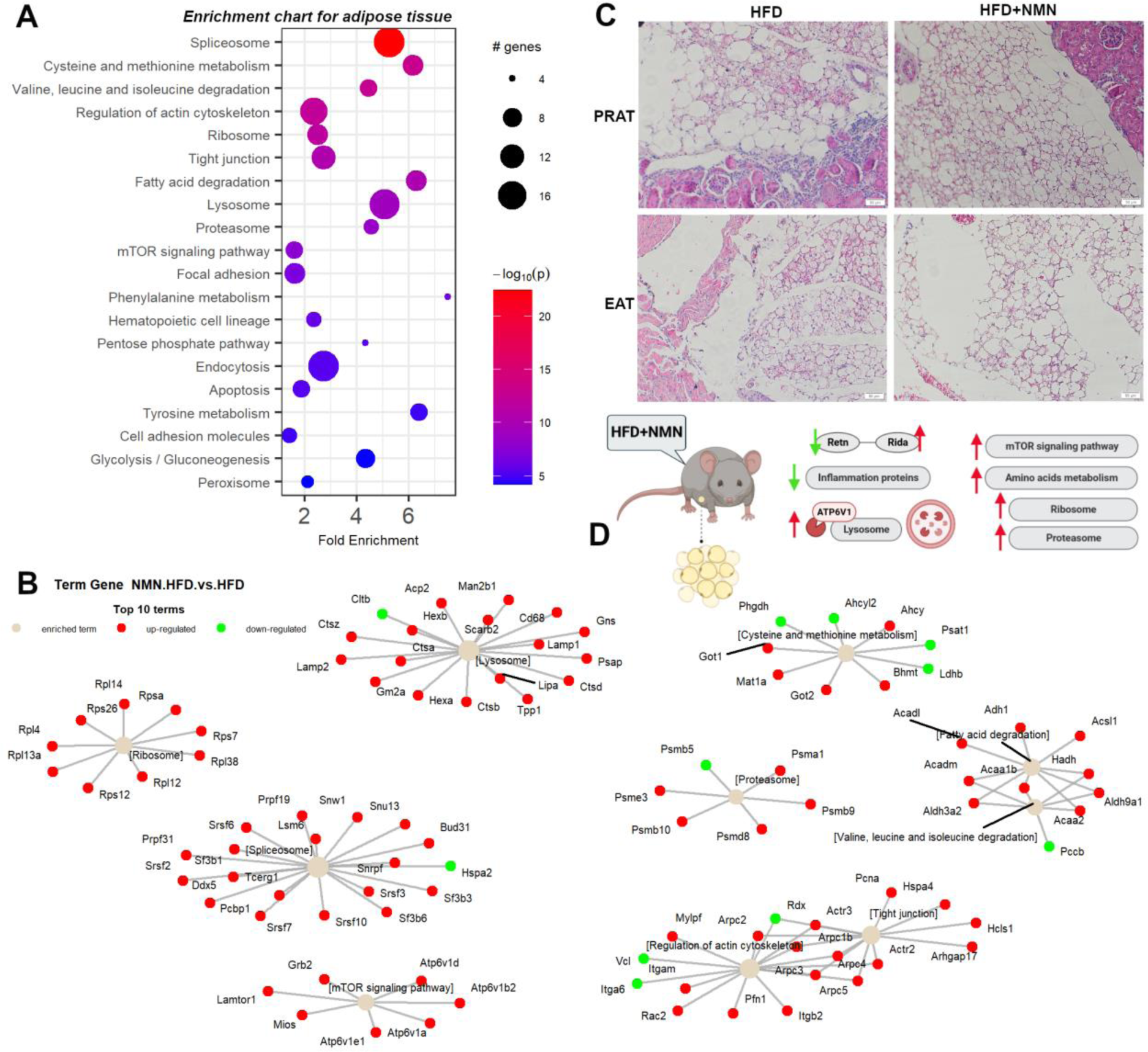
Integrated proteomics data analysis of NMN treated HFD mouse adipose tissue. (A) Enrichment chart for top 20 KEGG Pathways sorted by lowest *p* value in mouse adipose tissue. (B) Term-gene graph for top 10 terms. (C) Representative images of hematoxylin and eosin staining for perirenal adipose tissue (PRAT) and epicardial adipose tissue (EAT). Scale bars, 50 μm. (D) Experiment summary (BioRender).

In cysteine and methionine metabolism pathways, Bhmt, Mat1a, Ahcy, Got1, and Got2 were upregulated while Ahcyl2, Ldhb, Phgdh, and Psat1 were downregulated. Valine, leucine, and isoleucine degradation was altered with Acadm, Hadh, Acaa1b, Acaa2, Aldh3a2, and Aldh9a1 being upregulated, while Pccb was downregulated. At the level of Actin cytoskeleton (**Figure S20**) upregulation of Itgam, Itgb2, Rac2, Mylpf, Actr2, Actr3, Arpc1b, Arpc2, Arpc3, Arpc4, Arpc5, and Pfn1 was noticed, while Itga6, Rdx, and Vcl were down-regulated (**Figure 7**B). Ribosome pathway showed upregulation as component proteins like Rps7, Rps12, Rps26, Rpsa, Rpl4, Rpl12, Rpl13a, Rpl14, and Rpl38 where upregulated. At the level of Tight junctions, overexpression of Arhgap17, Pcna, Hspa4, Hcls1, Actr2, Actr3, Arpc1b, Arpc2, Arpc3, Arpc4, and Arpc5 was observed, while Rdx was downregulated. Fatty acid degradation showed upregulation by overexpression of Acaa1b, Acaa2, Hadh, Acadm, Acadl, Acsl1, Adh1, Aldh3a2, and Aldh9a1. Most of the Lysosome pathway components were upregulated (Ctsa, Ctsb, Ctsd, Ctsz, Tpp1, Hexa, Hexb, Man2b1, Gns, Lipa, Acp2, Psap, Gm2a, Lamp1, Lamp2, Cd68, and Scarb2) while Cltb was downregulated. Proteasome pathway (**Figure S21**) showed upregulation of Psmd8, Psme3, Psma1, Psmb9, and Psmb10, while Psmb5 was downregulated. The mTOR signaling pathway was upregulated in adipose tissue by overexpression of Atp6v1a, Atp6v1b2, Atp6v1d, Atp6v1e1, Lamtor1, Mios, and Grb2. Focal adhesion pathway showed upregulation of Lama2, Mylpf, Rac2, and Grb2, while Lama4, Itga6, Ilk, Vcl, and Cav1 were downregulated.

### 2.5. Down-regulated OXPHOS proteins and up-regulated ketone bodies production were shown in NMN treated HFD mice brain

We performed heatmap proteomics analysis (**Figure 8**A) of mouse brain samples for comparing the protein expression between HFD group and HFD+NMN treated group. We identified a total of 45 differentially expressed proteins with a false discovery rate (FDR) < 0.05 (Limma Test) Among these, 23 proteins were down-regulated and 14 proteins were up-regulated in the HFD+NMN group compared to HFD group. The most significantly down-regulated protein was Svs5 (seminal vesicle secretory protein 5) with a log_2_FC of −2.02 (FDR = 0.011). Other significantly down-regulated proteins included Tnnt3 (troponin T3, skeletal, fast), Glyat (glycine-N-acyltransferase), and Slc27a5 (solute carrier family 27 (fatty acid transporter), member 5) with log_2_FC of −1.23, −1.18, and −0.51, respectively. The most significantly up-regulated protein was Creld1 (cysteine-rich with EGF-like domains 1) with a log_2_FC of 0.80 (FDR = 0.04). Other significantly up-regulated proteins included Gnmt (glycine N-methyltransferase), Alb (albumin), and Abcb6 (ATP-binding cassette, sub-family B (MDR/TAP), member 6) with log_2_FC of 0.82, 0.33, and 0.38, respectively.

**Figure 8.**
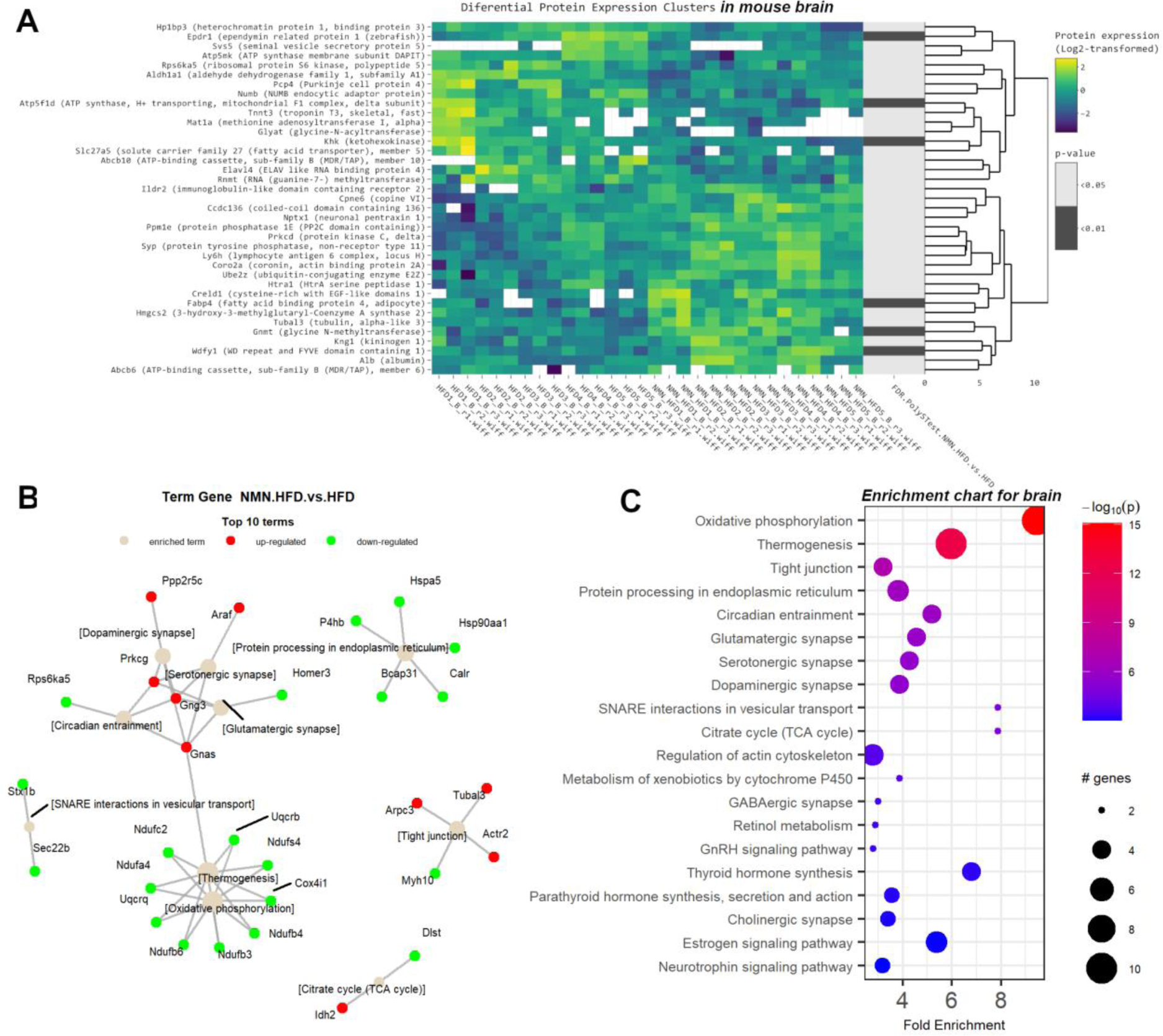
Significant changes induced by NMN treatment in the HFD mouse brain proteome. (A) Clustered heatmap of the 45 differentially expressed proteins, filtered with log_2_FC threshold set to exclude the interval −0.2, 0.2; 0.05 FDR threshold. Yellow color represents upregulation, while blue represents downregulation in HFD+NMN treated group compared to HFD group. From left to right, expression values (log_2_ transformed) for replicates (5 biological x 3 technical) are shown for the HFD group and for the HFD + NMN treated group, followed by significance values of the comparison to HFD group. (B) Term-gene graph for top 10 terms. (C) Enrichment chart for top 20 KEGG Pathways sorted by lowest *p* value.

Term gene graph (**Figure 8**B) and enrichment chart (**Figure 8**C) suggest that NMN treatment induces significant changes in the mouse brain proteome, respectively in the expression of proteins involved in various biological processes including fatty acid transport, energy metabolism, and signal transduction. These findings provide insights into the molecular mechanisms underlying the beneficial effects of HFD+NMN treatment on brain function. One of the most affected pathways was oxidative phosphorylation (**Figure S24**), with Ndufs4, Ndufa4, Ndufb3, Ndufb4, Ndufb6, Ndufc2, Uqcrb, Uqcrq, and Cox4i1 identified as downregulated proteins. Thermogenesis was mostly downregulated in brain, as it shares many proteins with OXPHOS pathway, except for Gnas identified as upregulated. Tight junctions (**Figure S25**) were also affected, with Actr2, Arpc3, and Tubal3 identified as upregulated and Myh10 downregulated. Protein processing in endoplasmic reticulum (**Figure S26**) members like Hspa5, Calr, P4hb, Bcap31, and Hsp90aa1 were upregulated. Circadian pathway was mostly upregulated by the member proteins Gng3, Gnas, and Prkcg, while Rps6ka5 was downregulated. Glutamatergic synapse and dopaminergic synapse were also affected, with Prkcg, Gnas, and Gng3 identified as upregulated proteins in both pathways. SNARE interactions in vesicular transport pathway were also modified, with Stx1b and Sec22b downregulated. Finally, citrate cycle (TCA cycle) had Idh2 identified as upregulated and Dlst downregulated.

### 2.6. NMN treatment decreased mitochondrial function with increased membrane potential and higher ROS production in muscle cells while in hepatic cells mitochondrial mass was higher and mitochondrial membrane potential was reduced

The results of the flow cytometry analysis for HepG2 cells and C2C12 myotubes in six experimental conditions are presented in the **Figure S1** and **Figure S2**. The conditions tested were normoglycemic (NN), hyperglycaemic followed by culture media switch to normoglycemic during NMN treatment (HN), hyperglycaemic before and during treatment (HH), treated with 100 μM NMN versus untreated, and the parameters analyzed were mitochondrial mass, mitochondrial function, mitochondrial membrane potential, mitochondrial ROS, intracellular neutral lipids and intracellular polar lipids.

In accordance with the *in vivo* experiment, we investigated the influence of the NMN treatment in HepG2 cells (as *in vitro* liver model for T2DM) by flow cytometry. NMN treatment led to a significant increase in the mitochondrial mass (**Figure S1**A) by 3.56% in NN condition (*p* < 0.05), 113% in HN condition (*p* < 0.0001), and 27.38% in HH condition (*p* < 0.01) respectively. There was no significant change in neutral lipids (**Figure S1**B), polar lipids (**Figure S1**C), mitochondrial function (**Figure S1**E) nor mitochondrial ROS (**Figure S1**G). Mitochondrial membrane potential (**Figure S1**F) was significantly decreased by 42.33% (*p* < 0.0001), and 33.91% (*p* < 0.01) respectively, in NN and HN conditions after NMN treatment. No significant changes in mitochondrial membrane potential were observed in the HH condition.

The effects of NMN treatment on mitochondrial parameters were evaluated in differentiated C2C12 myotubes. The statistical significance was calculated for each parameter by comparing the treated to untreated conditions. Regarding mitochondrial mass (**Figure S2**A) in the NN condition, NMN treatment resulted in a significant decrease in mitochondrial mass by 29.73% (*p* <0.01). However, NMN treatment induced no significant changes in mitochondrial mass, neutral lipids (**Figure S2**B) nor polar lipids (**Figure S2**C) NMN treatment led to a significant decrease in mitochondrial function (**Figure S2**E) by 14.77% (*p* < 0.0001) in NN conditions, 6.34% (*p* < 0.05) in HN condition, and 21.76% (*p* < 0.0001) in HH condition, respectively. Mitochondrial membrane potential (**Figure S2**F) was significantly decreased by 24.95% (*p* < 0.001) in NN condition, but significantly increased by 23.27% (*p* < 0.05) in HN condition after NMN treatment. However, no significant changes in mitochondrial membrane potential were induced by NMN treatment in HH condition. Mitochondrial ROS (**Figure S2**G) were significantly decreased by 8.43% and 6.11% (*p* < 0.01) in NN respectively HH conditions, and significantly increased by 11.39% (*p* < 0.0001) in HN condition after NMN treatment.

### 2.7. NMN down-regulates spliceosome proteins while up-regulating ribosome proteins in hepatocytes

The heatmap analysis of NMN treated HepG2 cells under HN conditions (**Figure S3**A) revealed downregulation of ATP5PD, ATPAF2, CERS4, CTNND1, EXOSC8, GAPD, H2A/k, H3-7, HEL-S-107, HIST1H4J, KEAP1, NDUFA3, RPF2, RPL13A, RPL15, RPL19, RPL29, and RPL35 proteins while DHCR24, GOT2, GRIK2, HGH1, HLA-Cw, KIAA0406, NEDD9, OK/KNS-cl.6, PNKD, SLC1A3, SSBP1, TOM1, TYMS, and UBE2K proteins were significantly upregulated.

On the other hand, heatmap analysis of NMN treated HepG2 cells under HH conditions (**Figure S3**B) revealed significant changes in the expression of 85 proteins with a false discovery rate (FDR) <0.05 and a log2FC below −0.3 or above 0.3. Of these, 56 proteins were upregulated, while 29 were downregulated in the HH condition treated with NMN (H100H) compared to untreated control (H0H condition). Among the upregulated proteins were AP2B1, FABP1, FAU, and GALT, while downregulated proteins included ATE1, AURKB, EPHX1, and NGEF. We observed changes in proteins involved in diverse biological processes, including protein synthesis (CDK105), signal transduction (G3BP), and carbohydrate metabolism (GOT2).

Pathway enrichment analysis (**Figure S4**B and **Figure S5**B) showed that the ribosome pathway (**Figure S27**) was significantly influenced in both HN and HH conditions. In HN condition (**Figure S4**C), RPS4X, RPS16, RPS20, and RPL22 were upregulated, while 29 ribosomal proteins were downregulated. In contrast, in HH condition (**Figure S5**A), RPS5, RPS6, RPS8, RPS9, RPS11, RPS13, RPS15, RPS18, RPS23, RPS24, RPS26, RPS28, RPS29, FAU, MRPL2, and 28 ribosomal proteins were upregulated, while MRPL12, MRPL11, MRPL13, MRPL14, MRPL15, MRPL17, MRPL19, and MRPL23 were downregulated.

The oxidative phosphorylation pathway (**Figure S28**) was also significantly influenced in both conditions, with NDUFB10 and COX5A upregulated in HN condition. Moreover, ATP6V1A and ATP6V1C1 were upregulated in HH condition. Interestingly, the OXPHOS pathway was affected in both HN and HH conditions, but the set of up- and downregulated proteins was different. In HN condition, NDUFB10 and COX5A were upregulated, while RPS6, NDUFS4, NDUFV1, NDUFA3, NDUFA9, and NDUFB11 were downregulated. In HH condition, RPS6KA3 and ACSL4 were upregulated, while TSC1, ACSL1, and many other proteins involved in oxidative phosphorylation were downregulated.

The spliceosome pathway was significantly influenced by NMN treatment only in HN condition, with DDX5, PHF5A, U2SURP, and HNRNPU upregulated and SNRPD1, PRPF8, and HSPA8 downregulated. In HH condition, several spliceosome proteins were downregulated, but the pathway was not significantly influenced (**Figures S5**A, **S5**B, and **S29**). Finally, the citrate cycle and valine, leucine and isoleucine degradation pathways were significantly influenced only in HH condition, with different sets of up- and downregulated proteins in each pathway.

### 2.8. NMN down regulates proteasome and up-regulates DNA replication and cell cycle pathways in muscle cells

Heatmap analysis of C2C12 myotubes in HN condition (**Figure S6**A) revealed differential expression of several proteins upon treatment with 100 μM NMN (H100N) compared to untreated condition (H0N). Out of the 63 proteins, the most significantly upregulated protein was Islr with a log_2_FC of 0.79 and an FDR value of 0.026. On the other hand, the most significantly downregulated protein was Hgsnat with a log_2_FC of −1.18 and an FDR value of 0.022. Among the upregulated proteins, Ankrd2 had the highest log_2_FC (0.35) and the lowest FDR value (1.2E-06). Among the downregulated proteins, Ash2l had the lowest log_2_FC (−0.89), while Casq1 had the lowest FDR value (0.0001).

Moreover, several proteins related to muscle function have shown significantly different expression levels in myotubes upon treatment with NMN (H100N) compared to H0N. Creatine kinase (Ckm) and myosin binding protein H (Mybph) were upregulated with log_2_FC of 0.36 and 0.33, respectively. Instead, Actin alpha 1, skeletal muscle (Acta1) was downregulated with a log_2_FC of −0.44. Other extracellular matrix proteins such as collagen, type I, alpha 1 (Col1a1) and collagen, type III, alpha 1 (Col3a1) were upregulated with log_2_FC of 0.34 and 0.43, respectively.

To investigate the effect of NMN treatment on C2C12 myotubes in hyperglycemic conditions (HH) we conducted proteomics data analysis by heatmap and pathway enrichment. Among the 30 proteins from H100H versus H0H comparison in heatmap (**Figure S6**B), 23 were upregulated and 7 were downregulated as the effect of NMN treatment. The protein with the largest upregulation was Abcb6 (ATP-binding cassette, sub-family B (MDR/TAP), member 6) with a log_2_FC of 0.624 and an FDR < 0.015. Other significantly upregulated proteins included Man1a2 (mannosidase, alpha, class 1A, member 2), A2m (PZP, alpha-2-macroglobulin like), and Vtn (vitronectin). On the other hand, the most repressed protein was Pdp1 (pyruvate dehydrogenase phosphatase catalytic subunit 1) with a log_2_FC of −0.953 and an FDR < 0.02. Other significantly downregulated proteins included Macroh2a2 (macroH2A.2 histone) and Heatr5b (HEAT repeat containing 5B).

To better understand the biological significance of the differentially expressed proteins identified, we performed pathway enrichment analysis using PathfindR. The results of the enrichment chart (**Figure S7**B and **Figure S8**A) show several functional pathways, including amino acid metabolism, cytoskeleton organization, protein processing and energy metabolism.

Term-gene graph revealed significant changes in protein expression in various pathways: ribosome-related proteins (**Figure S31**) were upregulated in the HN condition (**Figure S7**C), including Rps2, Rps4x, Rps6, Rps8, Rps9, Rps11, Rps12, Rps13, Rps18, Rps19, Rps23, Rps24, Rps25, Rps26, Rps29, Fau, Rpsa, Rpl3, Rpl4, Rpl6, Rpl7, Rpl7a, Rpl8, Rpl10a, Rpl13, Rpl13a, Rpl14, Rpl15, Rpl17, Rpl18a, Rpl19, Rpl21, Rpl22l1, Rpl23, Rpl23a, Rpl24, Rpl26, Rpl27, Rpl27a, Rpl28, Rpl29, Rpl32, Rpl34, Rpl35, Rpl36, Rpl37a, and Rpl39. In contrast, several ribosome-related proteins were downregulated, including Rps28, Mrpl4, Mrpl12, Mrpl19, Mrpl27, Rpl31, and Rplp1 in the HN condition. The cell cycle pathway was upregulated by NMN in the HH condition, including Cdk1, Rad21, Mad2l1, Pcna, Mcm2, Mcm3, Mcm5, Mcm6, and Mcm7 (**Figure S8**B and **Figure S32**).

Also, the oxidative phosphorylation pathway (**Figure S33**) was significantly influenced in both conditions, with Atp6v1b2 and Ppa2 upregulated in HN condition and Ndufs4, Ndufv2, Ndufb11, Uqcrb, Cox5b and Cox6c upregulated in HH condition. However, several other proteins were downregulated in the HN condition, including Ndufs1, Ndufs2, Ndufs3, Ndufs4, Ndufs8, Ndufv2, Ndufa2, Ndufa5, Ndufa9, Ndufa10, Ndufb4, Ndufb5, Ndufb6, Ndufb8, Ndufb10, Ndufb11, Sdhb, Sdhc, Uqcrfs1, Cyc1, Uqcrc1, Uqcrc2, Uqcrb, Uqcr10, Cox4i1, Cox5a, Cox5b, Cox6c, and Cox7a2, and Cycs in HH condition. DNA replication was upregulated in HH condition, including Pcna, Mcm2, Mcm3, Mcm5, Mcm6, and Mcm7.

## 3. Discussion

Global cost of treating type two diabetes mellitus and associated diseases is rising globally. Unlike other treatments, NMN previously showed great potential in treating T2DM [11] with no observed adverse effects. It was previously proposed that NMN as NAD^+^ precursor activates NAD^+^ dependent Sirt1 which in turn, through TFAM and c-Myc transcription factors induce mitochondrial biogenesis [12]. However, this seems to have a minor if not a totally absent effect, as trials in humans does not report the expected results [18]. Therefore, a discovery proteomics method was necessary in order to find hints of where is glucose going after NMN treatment. Proteins are stable, they are the effectors and regulators of all biological functions of cells in the living organisms [24]. Recent, Important advance in DIA data processing methods using artificial intelligence /neural networks, significantly improved proteome coverage and limit of quantification [25].

As T2DM hallmark is impaired glucose uptake, we performed differential proteomics on the tissues responsible for most of the glucose consumption: liver, muscle, adipose tissue, and brain.

In our cultured HepG2 cells T2DM model, although cardiolipin content (**Figure S1**A), as measure for mitochondrial mass, was significantly higher in both NMN treated conditions (HN and HH), mitochondrial function was not significantly affected (**Figure S1**E), while membrane potential was lower (**Figure S1**F). Notably, in HH conditions, all the OXPHOS proteins were down-regulated (**Figure S5**A, **Figure S28**), the same pattern being observed in HN conditions, where OXPHOS complex I component NDUFA3 (NADH: ubiquinone oxidoreductase subunit A3) and complex V subunits ATP5PD (ATP synthase peripheral stalk subunit d) and ATPAF2 (ATP synthase mitochondrial F1 complex assembly factor 2) were significantly repressed, while COX5a and NDUFb10 were significantly overexpressed by NMN treatment. Also, ATP6V1, a protein functioning as a channel which selectively allows protons to enter cell compartments like lysosomes was over expressed in HH conditions. In these hepatic cells, we also observed a down-regulated proteasome pathway in HH conditions (**Figure S3**B, **Figure S5**), whereas in the same conditions in myotubes this pathway was up-regulated (**Figure S8**). Interestingly, in HN and HH conditions, in HepG2 cells, ribosome components were over-expressed concomitant with repressed spliceosome components, except HSPA1B which was up-regulated (**Figure S4**C and **Figure S5**A). This is more similar with muscle cells HN conditions, and possibly NMN somehow influences the splicing machinery (**Figure S7**C) resulting abnormal proteins which are directed to chaperones for refolding or to proteasome for recycling [26]. In HepG2 HN conditions, spliceosome components are down-regulated along with processing in endoplasmic reticulum, amino acids degradation and fatty acids degradation (**Figure S3**A and **Figure S4**). Also, HSPA8 was repressed, as probably misfolded proteins were in a lower number and proteins synthesis in general was down-regulated. It worth mentioning that we found that in hepatocytes, NMN treatment in HN conditions (**Figure S3**A), induced an up-regulation of HLA-Cw, a leukocyte antigen (HLA) class I gene product, known to be involved in transplanted liver rejection [27]. This might rise concerns regarding NMN administration to liver transplant patients.

Our data show also up-regulated TYMS, also known as thymidylate synthetase, an enzyme catalysing the conversion of deoxy uridine monophosphate (dUMP) to deoxythymidine monophosphate (dTMP). It is involved in the regulation of DNA synthesis and cell proliferation. TYMS expression is upregulated in hepatocellular carcinoma, and its overexpression has been associated with poor prognosis and tumour progression [28]. SSBP1, also known as single-stranded DNA-binding protein 1, that plays a critical role in DNA replication, recombination, and repair as well as the maintenance of genome stability [29] and KIAA0406, also known as TELO2 interacting protein 1 (TTI1), playing a critical role in the regulation of DNA damage response and cell cycle progression [30] were found significantly up-regulated in HN conditions NMN treated HepG2 cells. Moreover, GRIK2 found upregulated in this condition, was previously reported as involved in regenerating liver after partial hepatectomy [31].

*In vivo*, our data showed similar down-regulation of spliceosome components in liver (**Figure 3** and **Figure S14**), also observed in liver and muscle cells *in vitro*. Our data agree with those of Jiao et all. [32] that proved that in contrast to the m^7^G cap, that has a stabilizing role of mRNA, the 5’end NAD^+^ capping of eukaryotic RNA targets the rapid decay of mRNA in mammalian cells through the DXO de-capping enzyme. Probably, the energetic state of cells could impact NAD^+^ capping and mRNA turnover [33].

In the mouse liver, ribosome components were also affected, in almost equal proportion, some components were up-regulated while others were down-regulated (**Figure 2, Figure 3**, and **Figure S10**), probably because Nolc1, also known as Nucleolar and Coiled-Body Phosphoprotein 1, a protein that localizes to the nucleolus and plays a role in ribosome biogenesis [34] (**Figure S12**), was down-regulated. Down-regulated Nolc1 and Clic4 (Chloride Intracellular Channel 4) are probably among the most important positive effects of NMN in liver, as it was previously reported that Nolc1 contributes to activation of hepatic stellate cells, which are key players in the development of liver fibrosis [35]. Both Nolc1 and Clic4 appear to be involved in the regulation of extracellular matrix production and may contribute to the progression of fibrosis [36]. Arachidonic acid metabolism, retinol, and fatty acid degradation were up-regulated, probably due to the increased NAD^+^ concentrations, that stimulated the desaturase activity of linoleic acid [37], the activities of retinol dehydrogenase and aldehyde dehydrogenases that generated all-trans-retinoic acid, an agonist of *α*, *β*, *γ* receptors for retinoic acids [38], as well as the activity of L-*β* -hydroxy acyl CoA dehydrogenase. Steroid hormone biosynthesis was also affected having some of the involved proteins up-regulated, and others down-regulated (**Figure 3, Figure S9**). Metabolism of xenobiotics by cytochrome P450 was up-regulated, suggesting a better hepatic detoxification activity of the liver after NMN treatment (**Figure S13**).

Notably, Mt-Cyb (Mitochondrially Encoded Cytochrome B protein) was over-expressed in the liver of the NMN treated mice (**Figure 2**), which could improve the electron carrier function of mitochondria. This effect was not observed in myotubes *in vitro*, probably suggesting that this mechanism occurs in other cells of muscle tissue but not in myocytes or it is dependent on a signalling pathway. Overexpression in muscle of NMN treated mice was noticed for: MPC1 (mitochondrial pyruvate carrier 1), NUP210L (nucleoporin 210-like), a protein that plays a role in the transport of macromolecules between the nucleus and cytoplasm, Cep43 (centrosomal protein 43), localized in centrosome, and involved in the regulation of microtubule dynamics and spindle formation during mitosis, Vanin 1 (VNN1) an enzyme involved in the regulation of oxidative stress and inflammation, Cell division cycle 27 (CDC27), a regulatory protein of the cell cycle, Trafficking protein particle complex 3 like (TRAPPC3L) that is involved in the transport of proteins between the endoplasmic reticulum (ER) and Golgi apparatus, all found over-expressed in the muscle of NMN treated mice (**Figure 4**). Also, an interesting case is Ferritin heavy polypeptide 1 (FTH1), a protein extremely important in iron storage and homeostasis [39], which is significantly over-expressed in liver tissue, but also in cultured myotubes (**Figure 2** and **Figure S6**).

Skeletal muscle plays a vital role in the regulation of post-prandial glucose levels. Following ingestion, approximately 80% of glucose is absorbed by skeletal muscle through a process known as insulin-dependent glucose uptake [40]. Both insulin-dependent and insulin-independent mechanisms are involved in the disposal of glucose by skeletal muscle. This process entails the delivery of glucose from the bloodstream to the muscle, its movement across the extracellular matrix towards the cell membrane, uptake facilitated by specialized glucose transporters present either continuously on the cell membrane or translocated in response to insulin or exercise stimuli. Furthermore, intracellular glucose metabolism influences the creation of a glucose gradient, thereby facilitating glucose transport within the muscle [41].

In the muscle tissue, our data show that nothing happens in terms of reducing insulin resistance, the expression of glucose receptor GLUT4 responsible for glucose internalisation in muscle cells is significantly reduced, the Glycogen Synthase (GS) enzyme is downregulated as well as most mitochondrial proteins in NMN treated mice, which is contrary to the NMN induced mitochondrial biogenesis hypothesis [12]. Our proteomic data strongly suggest that in the muscle tissue of HFD mice, NMN acts as a mild repressor for mitochondrial proteins. Skeletal muscle tissue is not made only by muscle cells, it might contain blood vessels, adipose cells, etc. Unexpectedly we have detected significant overexpression of UCP1 (**Figure 9**), which is a protein of the inner mitochondrial membrane of the brown adipose tissue, that acts as a regulated protons channel dissipating the proton gradient formed during the oxidation of NADH and FADH_2_ resulted from the metabolization of oxidable substrates. The energy of proton gradient is not used for ATP synthesis but for heat generation [42]. This result confirms a previous study correlating NMN administration with thermogenesis [43].

**Figure 9.**
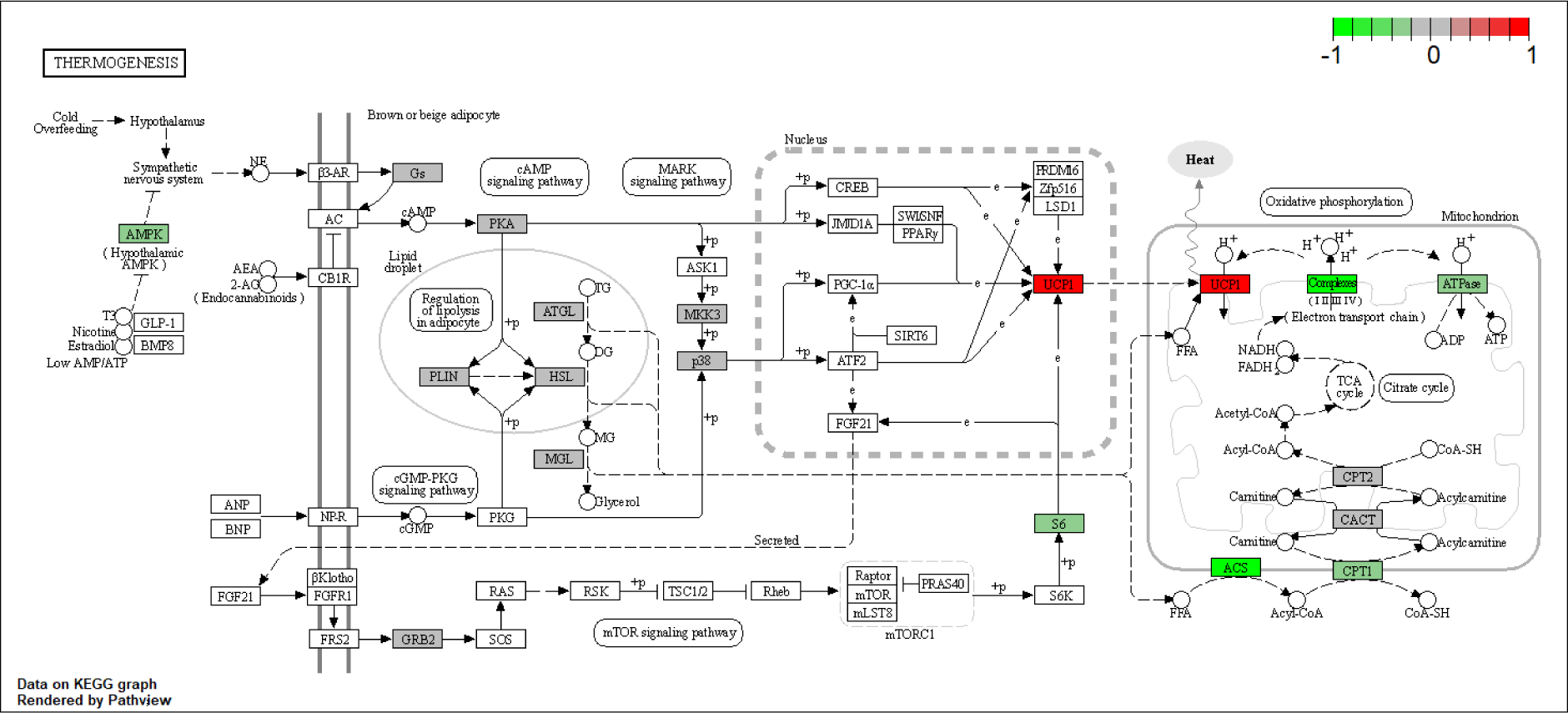
NMN treatment effects on the KEGG thermogenesis pathway in mouse muscle. The colour of the boxes represents the log2 fold change of the protein abundances, represented for HFD+NMN group versus HFD group comparison. Red: up-regulated; green: down-regulated; grey: no significant expression change.

To rule out the hypothesis that NMN could induce the expression of UCP1 in muscle cells, we used myotubes differentiated from C2C12 myoblasts in two different *in vitro* conditions, one with high glucose and high insulin growth medium during NMN treatment (HH), and the second with normoglycemic and normal insulin level medium (HN). None of the two conditions generated expression of UCP1, suggesting that, probably, in animals, NMN either stimulate the differentiation of preadipocytes infiltrated in skeletal muscle tissue into brown adipose cells or stimulates overexpression of UCP1 in existing brown adipose cells. These were not observed on the histology images of the muscle tissue (**Figure 5**B) probably 7 days of treatment was too short for these brown adipose cells to grow or multiply to a visible extent. as brown adipose cells were not observed in the skeletal muscle tissue. Furthermore, the examination of liver histology (**Figure 3**C) did not reveal the presence of brown adipose cells. It is known that BAT is present in most mammals, including mice and humans, in specific anatomical locations such as the interscapular region, as well as in proximity to organs such as the kidney, pancreas and heart. In the present study, both WAT and BAT were observed in both conditions, specifically the perirenal adipose tissue (PRAT) and epicardial adipose tissue (EAT), respectively. Examination of histology images (**Figure 7**C) featuring PRAT and EAT reveals a coexistence of WAT and BAT, implying a transitional state from WAT to BAT, a phenomenon previously described by other researchers as „brite” or beige [44]. Notably, a clear morphological distinction exists between the two adipose tissue types. WAT exhibits conspicuous lipid accumulations manifesting as large intracytoplasmic vacuoles, in contrast to BAT, which displays numerous fine lipid vacuoles with a brownish hue, hence the designation. These findings further underscore the functional diversity and metabolic adaptability of these adipose depots, thereby enhancing our comprehension of adipose tissue biology and its implications for energy homeostasis.

Mitochondrial mass evaluated by cardiolipin content (**Figure S2**A) was slightly lower in both HN and HH NMN treated cells. Also, mitochondrial function was reduced in both HN (*p* <0.05) and HH (*p* <0.0001) conditions while mitochondrial membrane potential was increased significantly in HN condition (*p* <0.05) and slightly but without statistical significance in HH condition (**Figures S2**D-S2F). Mitochondrial reactive oxygen species production was increased by NMN treatment in HN condition (*p* <0.0001) and decreased in HH condition (*p* <0.01) (**Figure S2**G). The expression of OXPHOS proteins (**Figure S33**) in HN condition was repressed by NMN treatment, the same effect being observed *in vivo* in similar conditions.

Decreased mitochondrial function with increased membrane potential and higher ROS production in muscle cells (**Figures S2**E-G) could be corelated with decreased Txnip (thioredoxin interacting protein) (**Figure S6**A): Txnip is a protein that binds to and inhibits the activity of thioredoxin, which is involved in redox regulation [45]. It is a negative regulator of glucose uptake, and its downregulation has been shown to increase glucose uptake in muscle cells [46] and increase in reactive oxygen species (ROS) production [47]. It is highly conserved and plays a key role in regulating cellular redox balance and was shown to extend lifespan of fruit flies [48]. Txnip was clearly observed down-regulated in our data along with improved glucose uptake, but not in muscle tissue.

Protein synthesis was clearly up-regulated in all NMN treated muscle cells. Interestingly, degradation of proteins in the framework of proteasome pathway (**Figure S34**) was repressed if the high glucose and high insulin levels were maintained during treatment and was up-regulated if normoglycemic medium was used during exposure. Also, TCA cycle proteins and fatty acid degradation were slightly down-regulated in HN conditions, not significantly affected in HH conditions.

During this discovery proteomics study, in the treated HN myotubes a down regulation of the spliceosome pathway (**Figure S35**), corelated with upregulation of chaperones (Hspa8, Hspa11) and proteasome proteins in the treated HN myotubes which suggest that NMN could lead to abnormal protein synthesis which activates chaperones and proteasomes for corrective actions (refolding, respective degradation). Also, NMN treatment leads to higher NAD^+^ concentration which in turn might cause the NAD^+^ capping followed by rapid decay of mRNA through the DXO de-capping enzyme. previously described by Jiao et all. [32]. However, in muscle cells, NMN treatment in the high glucose condition (HH) clearly shows down regulated proteasome and up-regulated DNA replication and cell cycle pathways (**Figure S8**B) suggesting that cells’ metabolism is on the growth and replication program. Overexpression of Znf827 (zinc finger protein 827) (**Figure S6**B) could suggest that NMN stimulates myoblasts differentiation of cultured cells in HH conditions. HN conditions treated cells showed up-regulated purine metabolism and collagen synthesis, and, interestingly, a component of kinetochore, Zwilch (Zwilch Kinetochore Protein) was up-regulated. Also, Hmg20a, a critical regulator of muscle differentiation and regeneration [49], was upregulated. There are a few other interesting protein expression differences in cultured myotubes shown on the heatmap (**Figure S6**) which might be useful as starting point for other studies. However, so far, we rule out the mitochondrial biogenesis hypothesis as main NMN effect in muscle.

A further tissue that consumes a considerable amount of glucose is white adipose tissue, whose metabolism undergoes significant alterations in individuals with type 2 diabetes, leading to dysregulation of lipid storage, release, and adipokine secretion, contributing to the pathogenesis of the disease [50]. In NMN treated mice, proteins from mTOR cell growth pathway (**Figure S22**) are overexpressed, as well as proteins involved in amino acids and protein synthesis and degradation (**Figure 6**). All identified proteins involved in lysosomal pathway (**Figure 11**) are strongly overexpressed, corelated with overexpression of ATP6V1 which is required for the acidification of lysosomes, necessary for the activation of lysosomal lipases [51]. Interestingly, ATP6V1 was overexpressed in NMN myotubes in culture (**Figure S7**C). This hydrolyses the triglycerides stored in lipid droplets into free fatty acids and glycerol, which can then be used as an energy source. Adipose tissue is composed of adipocytes and a stromal vascular fraction (SVF) that includes preadipocytes, immune cells, and endothelial cells. Overexpression of tight junction proteins (**Figure S23**) facilitate vascular permeability [52]. Inflammation related proteins (Cfd (complement factor D (adipsin)), Gm1088(immunoglobulin kappa variable 5-48), Igk-V19-17 (immunoglobulin kappa variable 6-17)) are down regulated (**Figure 6**).

**Figure 10.**
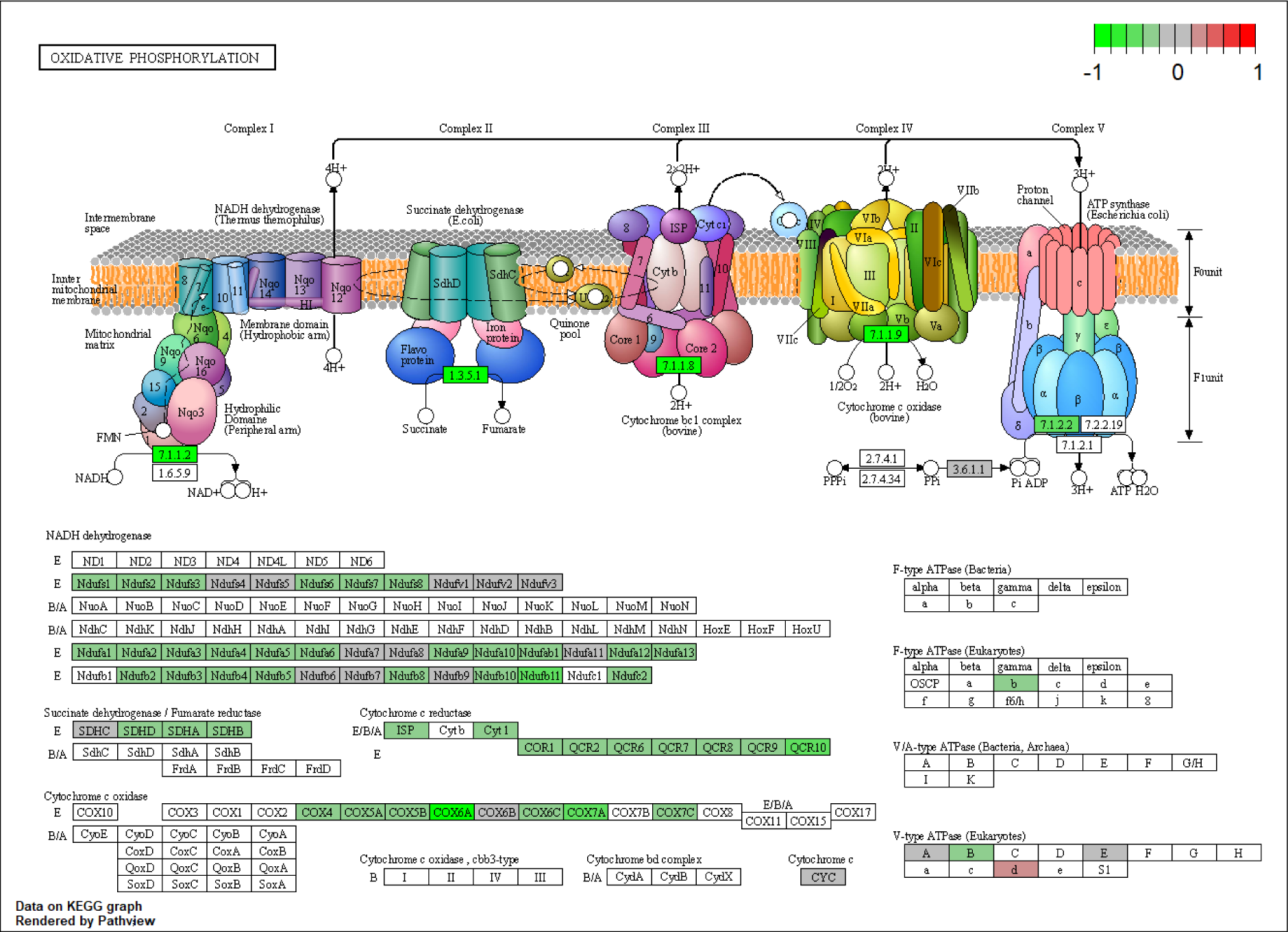
NMN treatment effects on the KEGG oxidative phosphorylation pathway in mouse muscle tissue. The colour of the boxes represents the log2 fold change of the protein abundances, represented for HFD+NMN group versus HFD group comparison. Red: up-regulated; green: down-regulated; grey: no significant expression change

**Figure 11.**
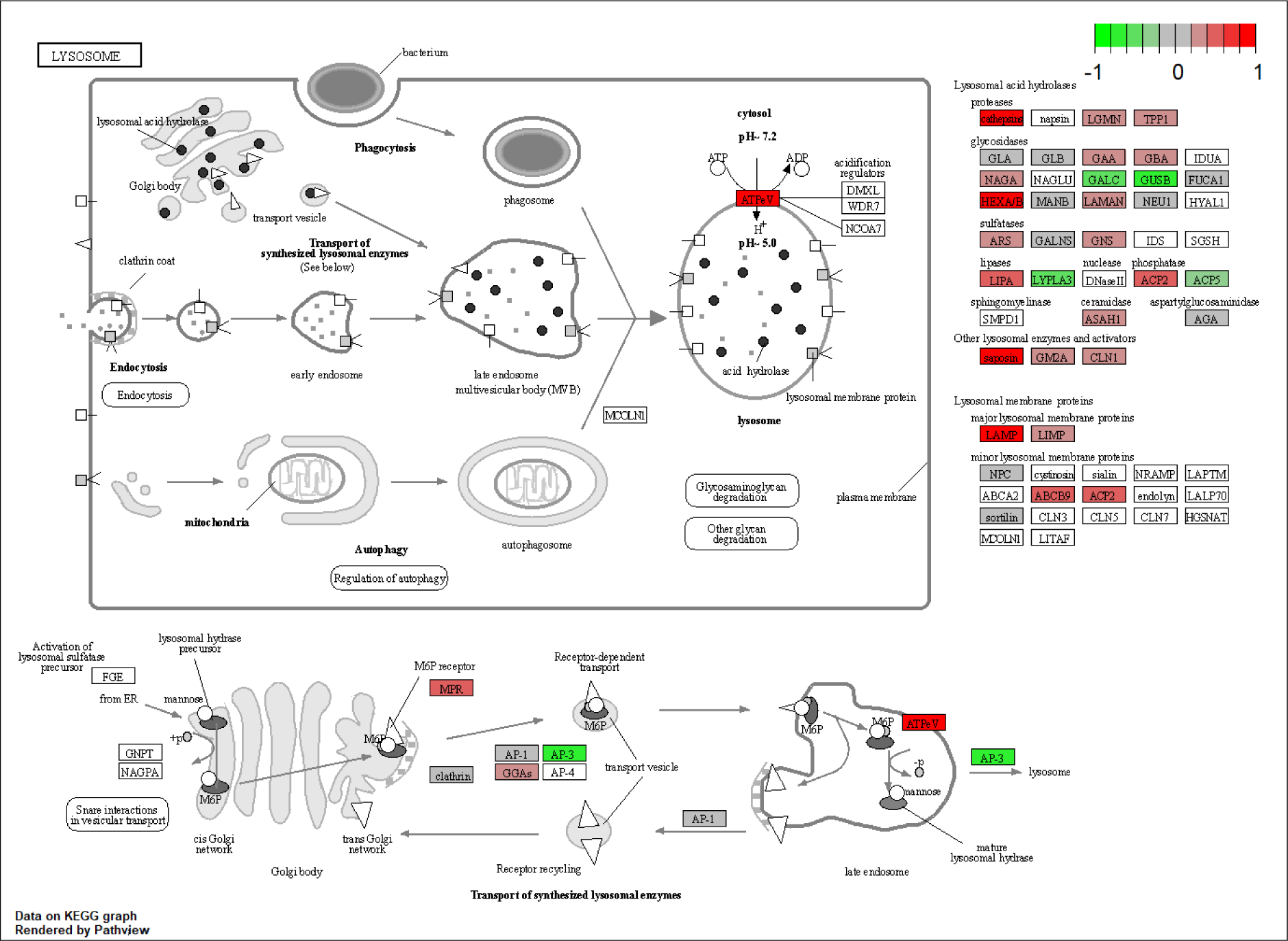
NMN treatment effects on the KEGG lysosome pathway in mouse adipose tissue. The colour of the boxes represents the log2 fold change of the protein abundances, represented for HFD+NMN group versus HFD group comparison. Red: up-regulated; green: down-regulated; grey: no significant expression change

Retn (Resistin) is an adipokine hormone that is mainly secreted by adipose tissue and is involved in insulin resistance and inflammation [53]. Our data show reduced expression level of resistin, corelated with up-regulated Rida (reactive intermediate imine deaminase A homolog) which was previously positively correlated with identified as repressed in insulin resistance [54]. We consider that this is the way NMN works in T2DM, and not as previously claimed by mitochondrial biogenesis. NMN treatment may have a beneficial effect on insulin sensitivity and inflammation by decreasing the expression level of Resistin and upregulating Rida. To our surprise, this is the only proteomics study which managed to relatively quantify Resistin by mass spectrometry.

The brain is also actively involved in the uptake and utilization of glucose, its primary energy source [55]. In NMN treated HFD mice, the brain tissue proteomic data (**Figure 8** and **Figure S24**) revealed down-regulated mitochondrial OXPHOS, Atp5mk (ATP synthase membrane subunit DAPIT) and ATP5F1D (ATP synthase, H^+^ transporting, mitochondrial F1 complex, delta subunit) being the most repressed, Tight junctions components were up-regulated (**Figure S25**), similar to the adipose tissue (**Figures 8**B, **8**C and **Figure 7**B) while SNARE interactions in vesicular transport pathway was down-regulated, as well as protein processing in endoplasmic reticulum (**Figure S26**). Purkinje cell protein 4 (Pcp4) is primarily expressed in the cerebellum, specifically in Purkinje cells, which are a type of neuron involved in motor control and coordination [56] and was downregulated in NMN treated mice brain tissue. During neural development, Numb (NUMB endocytic adaptor protein) is involved in regulating cell fate decisions by asymmetrically segregating into one daughter cell during cell division. This process, known as "Numb-mediated asymmetric cell division" leads to one daughter cell retaining stem cell properties, while the other becomes a neuron or glial cell [57]. In the context of neural development, repression of Numb (which is also our case) has been associated with an increase in symmetric cell division [58], where both daughter cells retain stem cell properties. Moreover, our studies revealed a downregulation of Ketohexokinase (KHK) in NMN treated mice. This is an enzyme primarily involved in the metabolism of fructose and implicated in the pathogenesis of Alzheimer’s disease [59]. Previous studies have suggested that fructose may contribute to the development of Alzheimer’s disease by increasing the production of amyloid beta, involved in the formation of amyloid plaques in brain [60]. By catalysing the phosphorylation of fructose, KHK generates accumulation of glyceraldehyde, a metabolite that has been shown to increase the production of amyloid beta [61]. Decreased levels of KHK, as shown by our data, might be a positive NMN effect in the brain.

Also, solute carrier family 27 member 5 (SLC27A5) has been implicated in the regulation of the levels of very long-chain fatty acids (VLCFAs) in brain. SLC27A5 could play a role in the transport of VLCFAs into astrocytes, where they can be metabolized and incorporated into myelin. In addition, SLC27A5 has been shown to be upregulated in response to certain pathological conditions [62], and in our experiments, it is down-regulated by NMN treatment.

ATP-binding cassette sub-family B member 10 (ABCB10) transporter is also downregulated by NMN exposure. ABCB10 has been shown to be highly expressed in neurons, and it has been suggested to play a role in the regulation of mitochondrial function in these cells [63]. It has been implicated in the response to oxidative stress, which is a common feature of many neurological disorders [64]. Its reduced level may be due to the reduced level of mitochondrial OXPHOS protein which in turn may cause lower oxidative stress and lower need for ABCB10, because the higher level of NAD^+^. The same mechanism might explain the observed reduced level of ELAV-like RNA-binding protein 4 (ELAVL4), previous studies suggesting that ELAVL4 may also play a role in the regulation of the expression of genes involved in the response to oxidative stress [65]. Also, ATP-binding cassette, sub-family B (MDR/TAP), member 6 (ABCB6) was found up-regulated, and is known as transporter of hem (a known source of reactive oxygen species) out of the cells. This might be another protecting effect, as a mild repressed mitochondrion might require lower levels of hem to function. Cytosol produced hem, not needed in mitochondria, is this way exported through overexpressed ABCB6 carrier [66]. Also, albumin was overexpressed, which was previously shown to have antioxidant role in the brain, acting like a free radical’s scavenger [67].

Wdfy1 (WD repeat and FYVE domain containing 1) is a large multidomain protein found up-regulated. It is involved in the endosomal trafficking, and seems to play a role in the maturation and trafficking of endosomes, but also in autophagy [68]. Up-regulated Kininogen 1 (Kng1) has also been shown to have some neuroprotective effects in the brain. It has been suggested that Kng1 may be involved in the regulation of cerebral blood flow and the protection of neurons from oxidative stress and inflammation [69]. Gnmt (glycine N-methyltransferase), found to be up-regulated by NMN treatment, is an enzyme primarily involved in the metabolism of glycine, but also in regulation of S-adenosylmethionine (SAM) levels. SAM is an important methyl donor in various biological processes, including the methylation of DNA and histones, which can affect gene expression [70]. Gnmt is involved in the catabolism of SAM, regulating SAM levels in the brain [71]. Decreased Gnmt expression has been observed with age and neurodegenerative diseases [72]. Also, TUBAL3, found up-regulated in our experiments, is involved in the regulation of microtubule stability and organization which are critical for proper neural progenitor cell division and migration [73]. Over-expression of Creld1 (cysteine-rich with EGF-like domains 1), known to be expressed in neural progenitor cells (NPCs) [74] suggest that NMN treatment might slightly stimulates growing population of NPC cells.

Interestingly, 3-hydroxy-3-methylglutaryl-Coenzyme A synthase 2 (HMGCS2), an enzyme involved in the synthesis of ketone bodies, which are important metabolic fuels for brain during periods of fasting or low glucose availability [75], is up-regulated in the brain of NMN treated mice. This is interesting, as these mice were not fasting nor exercising, glucose should be available as brain energy source and no need for increased ketone bodies production should exist. However, this might be an adaptation to reduced ATP production in mitochondria, also observed in our data and discussed above. Improved turnover of synaptic proteins might be induced by NMN treatment, as Ube2z (ubiquitin-conjugating enzyme E2Z) [76] is significantly up-regulated.

There are also several other up-regulated proteins involved in synaptic plasticity, neuronal survival, neural development and neuroinflammation: Prkcd (protein kinase C, delta), Syp (protein tyrosine phosphatase, non-receptor type 11) [77], Ly6h (lymphocyte antigen 6 complex, locus H), Coro2a (coronin, actin binding protein 2A), Ppm1e (protein phosphatase 1E), Nptx1 (neuronal pentraxin 1), Cpne6 (copine 6), Ildr2 (immunoglobulin-like domain containing receptor 2). Our data also show that NMN induced overexpression of Ccdc136 (coiled-coil domain containing 136) in the brain, however, almost nothing is known about this protein, its structure was software predicted, but its function is unknown [78].

## 4. Conclusions

Overall, improved glucose uptake observed after NMN treatment seems to be caused mainly by effects in the adipose tissue: Resistin down-regulation and increased protein synthesis and degradation, fatty acid degradation, Lysosome proteins up-regulation (most notably up-regulation of the ATP6V1 proton pump) along with mTOR cell proliferation signalling in white adipose tissue; differentiation of preadipocytes to brown adipose cells and/or over-expression of thermogenic UCP1. A series of other effects in an organ type dependent manner were also observed. Among these, it worth mentioning: spliceosome down-regulation corelated with up-regulated chaperones, proteasomes and ribosomes in liver and muscle cells, resulting in a slightly impaired and energy inefficient protein synthesis machinery; Increased production of ketone bodies through Hmgcs2, down regulating of some mitochondrial OXPHOS components and TCA cycle in brain. Overexpression of proteins involved in the metabolism of xenobiotics in liver. Notably, our data strongly suggest that NMN is not acting through mitochondrial biogenesis, the opposite seems to be the case, having a mild repressing effect on mitochondria, inducing known positive effects like those observed in animals during fasting. As a discovery study, this work was aimed to provide a clear picture of the NMN treatment effect in T2DB and a starting point for further investigations in order to deeper elucidate the mechanisms responsible for the most interesting of the observed effects.

## 5. Material and methods

### 5.1. Animal experiments and NMN treatment

For HFD model experiments, male C57BL/6J mice of 20 weeks of age were obtained from SPF Animal Facility. Mice were randomly divided into two groups, which consisted of 5 mice each. The animals were placed in open cages, provided with standard laboratory diet and water *ad libitum*. All animals were grouped and housed in an environmentally controlled room with temperature between 19 °C and 23 °C and 30% to 70% relative humidity, with a 12-hour light–dark cycle for an acclimation period of 7 days prior to the beginning of the experiment. After acclimation, both groups were fed with a high fat diet (HFD) containing 60% of the total calories from lard for a period of eight months in order to induce type 2 diabetes. The mice were considered diabetic when blood glucose levels at two hours’ time point after intraperitoneal glucose tolerance test (IPGTTs) was higher than 200 mg /dL. After 7 days of NMN treatment (intraperitoneally administrated 500 mg NMN per kg body weight /day) only for one group (HFD+NMN) the mice were euthanised. All experimental procedures have been executed in agreement with the regulations of the bioethical committee (authorisation # 01/2017) of the Independent Research Association (Bucharest, Romania) and with the European legislation concerning experiments performed using live animals.

### 5.2. Serum levels of triglycerides and cholesterol

Before euthanasia, blood samples were collected from the retro-orbital plexus for biochemical analysis of triglycerides and cholesterol using VetTest CHOL and VetTest TRIG strips with a clinical IDEXX VetTest Chemistry Analyzer System.

### 5.3. Histochemical observation

Immediately after euthanizing the mice, tissue samples from skeletal muscle, liver, white adipose tissues and brain were subsequently snap frozen by immersion in liquid nitrogen vapours and stored at – 80 °C for proteomic analyses and small pieces were fixed in 4% paraformaldehyde solution in phosphate buffer 0.05 M, pH 7.4, for a period of 48 h. After fixation, the specimens were embedded in paraffin wax. Sections (3–4 µm) were prepared and stained with haematoxylin and eosin (HE) for histological examination. Images were captured with Olympus BX41 microscope. Images were generated using Olympus DP25 Camera (Cell B software).

### 5.4. Cell culture, differentiation and NMN treatment

HepG2 cell line (ATCC HB-8065) was used as an *in vitro* model for liver, while C2C12 cell line (ATCC CRL-1772) was used as a model for muscle. The cells were grown in DMEM medium (Gibco, REF 31600-083) supplemented with 10% fetal bovine serum (FBS, Gibco), without addition of antibiotics, in a humidified atmosphere with 5% CO2 at 37°C.

At 70% confluence, C2C12 myoblasts were differentiated by serum deprivation using DMEM medium and 2% horse serum (Horse Serum, Donor Herd, H1270, Sigma). After 3 days of differentiation, myotubes were grown for 3 days in hyperglycaemic medium (DMEM + 30 mM glucose + 100 nM insulin), to induce insulin resistance. Since subsequent studies investigated the effects of NMN on mitochondrial activity and biogenesis, the culture medium was not supplemented with antibiotics (aminoglycoside antibiotics such as streptomycin have mitochondrial toxicity effects) to avoid affecting the aerobic metabolism of the cell. The culture medium was changed every 24 hours. On the third day of culture, cells grown under hyperglycaemic conditions were divided into two groups, so that during the NMN treatment, one part of the myotubes remained in the hyperglycaemic conditions, while the medium for the other part was changed to the specific normoglycemic condition (**Figure S7**A). Thus, two experimental conditions were obtained: cells grown in hyperglycaemic and hyper insulinemic medium switched to normoglycemic during the treatment (HN), and cells grown in hyperglycaemic medium before and during the treatment (HH). The cells were maintained with NMN treatment for 24 hours. Control cells were grown similarly in parallel corresponding to each of the three conditions but no NMN treatment. The same procedure was applied to the HepG2 cells (**Figure S4**A).

After treatment, the cells were harvested to assess the effects of NMN treatment. To determine the relevant concentration of NMN in the culture, initially, five concentrations were tested for cells viability and cytotoxicity: 50, 100, 500, 1000, and 5000 µM. For subsequent experiments, 100 µM NMN was selected as representative and consistent with other published studies. Cell cultures in all experiments were from passages 5 and 8 for the C2C12 line respective passage 20 for the HepG2 line.

### 5.5. Flow cytometry

After 24 h treatment with or without 100 µM NMN in HN and HH conditions, HepG2 cells and respectively C2C12 myotubes were harvested and washed twice in cold PBS, and incubated with different fluorescent dye solutions in culture media without phenol red and serum for 30 min at 37 °C in the dark. The fluorescent markers used were as follows: 10 nM NAO (sc-214487, Santa Cruz Biotechnology) for mitochondrial mass, 25 nM MitoView Red for mitochondrial function, 2 µg /mL JC-1 for mitochondrial membrane potential, 2.8 µg /mL DHR123 for mitochondrial ROS and 100 ng /mL Nile Red (7726.1, Roth) for neutral and polar lipids. After dye incubations, cells were washed with PBS and resuspended in 500 μL culture media without phenol red. The stained cells were analysed on a Cytomics FC 500 flow cytometer (Beckman) (FL1 /FL2 /FL3 /FL4 /FL5) using Flowing software 2.5.1. version. For each sample, three replicates of 15 000 events each were acquired.

### 5.6. Fluorescence microscopy

For microscopic visualization of metabolically active mitochondria, mitochondrial membrane potential, and mitochondrial ROS, cells were cultured in 96-well plates - black/clear Sterile Imaging Plate (Falcon, BD 353219). After inducing insulin resistance and NMN treatment, the culture medium was aspirated from each well, followed by two washes with PBS. Subsequently, cells were separately incubated with 100 µL per well of MitoView Red (25 nM), JC-1 (2 µg/mL), and DHR123 (2.8 µg/mL) (GeneCopoeia) diluted in serum-free culture medium at 37°C in the dark. After 30 minutes, the staining solution was removed, and the cell surface was washed twice with PBS. Following staining, cells were maintained in phenol red-free culture medium (normoglycemic or hyperglycaemic), and were visualized using an Olympus IX73 inverted fluorescence microscope.

### 5.7. Sample preparation for mass spectrometry analysis

After NMN treatment, tissue and cell samples were minced and homogenised in ice-cold lysis buffer (containing 8 M urea, 5 mM EDTA, 25 μg /mL spermine, 25 μg /mL spermidine, 1 mM PMSF, in 50 mM Tris-HCl pH 7.8) using a Retsch MM400 homogenizer with stainless steel balls (⌀ 2 mm) for 5 min at ν = 30. The crude homogenate obtained after centrifugation was further centrifuged at 17 000 x *g* for 10 min at 4°C to isolate the protein fraction. The protein quantification was performed using the Bradford method and a bovine serum albumin 6 points (0.125 – 1.5 μg /μL) standard curve.

For proteolysis, a volume corresponding to 30 μg protein was diluted in 50 mM NH_4_HCO_3_ to a final volume of 500 μL, resulting in a urea concentration of 1.5 M, in low binding 1.5 mL tubes. The samples were then incubated with 25 μL of 100 mM DTT in 100 mM NH_4_HCO_3_ for 45 minutes at 37 °C. To carry out alkylation, 26.25 μL of 300 mM IAA in 100 mM NH_4_HCO_3_ was added and incubated for 45 minutes at 37 °C in the absence of light. Trypsin digestion was performed using a trypsin solution with a concentration of 1 μg /μL (Trypsin Gold, V528A, Promega), and 0.6 μL of the trypsin solution was added to achieve a final ratio of 1:50. The mixture was left overnight at 37 °C with gentle shaking. The digestion process was stopped by adding 10 μL of 10% trifluoroacetic acid. The resulting peptides were purified using Spec plus C18 tips, speed-vacuum dried and reconstituted in 30 μL of a solution containing 2% acetonitrile and 0.1% formic acid. Finally, the samples were transferred to a clean autosampler vial with an insert for LC-MS/MS analysis.

### 5.8. Identification of proteins by liquid-chromatography-mass spectrometry analysis

The peptides samples were subjected to LC-MS/MS analysis using an AB SCIEX TRIPLE TOF 5600+ mass spectrometer and separated with a NanoLC 425 system (Eksigent). The setup included an analytical column (Eksigent 5C18-CL-120, 300 μM ID, 150 mm length) connected to DuoSpray ion source (AB Sciex). A volume of 5 μL of the peptide samples was loaded using Solvent A (0.1% formic acid) and eluted with a gradient from 5% to 90% Solvent B (0.1% formic acid in acetonitrile) over 90 minutes at a flow rate of 5 μL /min, with a column temperature of 55 °C. Each sample was analysed in triplicate. Electrospray ionization in positive ion mode was used, with the ion spray voltage of 5500 V and a source temperature of 200 °C. The TRIPLE TOF 5600+ was operated in DIA SWATH-MS mode with 64 variable windows. The MS1 survey scan ranged from 400 to 1250 m/z, while the MS2 spectra were acquired in high-sensitivity mode from 100 to 2000 m/z. The accumulation time was set to 0.049 s, and the ion scan was sampled in 55 ms time windows in high-sensitivity mode, resulting in a cycle time of 3.5 seconds.

The mass spectrometry proteomics data have been deposited to the ProteomeXchange Consortium via the PRIDE [79] partner repository with the dataset identifier PXD043257. Reviewer account: Username; Password: M0KNo3iI.

### 5.9. Data analysis

Protein identification from DIA data was carried out in a label-free and library-free manner using DIA-NN ver. 1.8.1 [25]. The raw spectra were searched against the fasta file containing the complete mouse reference proteome (UniProt, UP000000589, November 2022, 55 275 entries) for mouse tissue sample and C2C12 myotubes, and against the complete human reference proteome (UniProt, UP000005640, November 2022, 82 492 entries) for HepG2 cells, with a precursor m/z range of 400 to 1250, and trypsin was chosen as the digestion enzyme. The data was searched with MBR enabled and robust LC was employed as the quantification strategy, with a false discovery rate (FDR) set at 0.01. C carbamidomethylation and Ox(M) modifications were considered during the search. Retention time-dependent normalization was applied, and quantitation was performed using the MaxLFQ [80] algorithm in DIA-NN.

Statistical and differential downstream analysis was conducted using the locally installed PolySTest version 1.3 (release) [20], which can be downloaded from the following link: https://bitbucket.org/veitveit/polystest/src/master/. The analysis utilized as input the unique gene matrix *tsv file obtained from DIA-NN. For significance determination, an FDR-adjusted *p*-value threshold of 0.05 was applied, along with a log_2_ fold change (log_2_FC) threshold of −0.2-0.2, or −0.3-0.3, or −0.5-0.5 to identify significantly downregulated or upregulated proteins, respectively. In PolySTest, Limma test was used for statistical significance analysis, as it is well-suited for gene or protein expression data and handles well the missing values in some replicates. Following the statistical analysis, data visualization, including heatmaps and expression profiles, were generated.

Each gene identified through DIA-NN was converted and mapped to its corresponding protein object using the ’org.Mm.eg.db’ and ’org.Hs.eg.db’ packages in Bioconductor [81,82]. The proteomic data was analysed and graphically plotted primarily using the R Studio platform (version 4.2.2).

For the differential pathway expression analysis (PEA), the list of proteins with their respective Log2 fold change and statistical significance was exported from PolyStest and imported into R, for further analysis using PathfindR version 1.6.4 [83]. PathfindR utilizes a protein-protein interaction network (PIN) analysis approach with PIN data for *Homo sapiens*. For mouse tissues and C2C12 myotubes the code was optimized to take PIN data for *Mus musculus* obtained from STRING (https://stringdb-static.org/download/protein.links.v11.5/), taxon id 10090. The output from PathfindR is a table that displays enriched pathways identified from the protein list, including fold enrichment values, the lowest and highest *p*-values generated from each pathway analysis iteration, and the upregulated and downregulated proteins associated with each pathway [83]. Additionally, we generated an enrichment chart and term graph for the top 20 and top 10 KEGG pathways, respectively, sorted by lowest *p*-value. For data integration and visualization of the main biological processes, we utilized the KEGG pathway database [21–23] (https://www.genome.jp/kegg/pathway.html) and employed the Pathview R package version 1.38.0 [84].

## Ethics

All experimental procedures have been executed in agreement with the regulations of the bioethical committee (authorisation # 01/2017) of the Independent Research Association (Bucharest, Romania) and with the European legislation concerning experiments performed using live animals.

## Declaration of AI use

We have not used AI-assisted technologies in order to generate any text for this manuscript.

## Data accessibility

The mass spectrometry proteomics data have been deposited to the ProteomeXchange Consortium via the PRIDE [79] partner repository with the dataset identifier PXD043257.

## Author Contributions

Conceptualization, G.C.M. and R.G.P.; Methodology, R.G.P and G.C.M; Investigation, R.G.P., T.S., E.V., and G.C.M.; Writing – Original Draft, R.G.P. and G.C.M.; Writing – Review & Editing, G.C.M. and A.D.; Funding Acquisition, G.C.M.; Resources, G.C.M., T.S., and E.V.; Supervision, G.C.M. and A.D.

## Acknowledgements

This study was supported by Independent Research Association, Bucharest - 012416, Romania.

## Declaration of interests

The authors declare no competing interests.

## Notes

https://www.ebi.ac.uk/pride/archive?keyword=PXD043257

### Competing Interest Statement

The authors have declared no competing interest.

