## Supplementary material for "NMN Works in HFD-Induced T2DM by Interesting Effects in Adipose Tissue, and Not by Mitochondrial Biogenesis": Document_S1.pdf

**Figure S1.** NMN effects on HepG2 cells exposed to hyperglycemic conditions: neutral lipids, polar lipids, mitochondrial mass, metabolic activity, membrane potential and reactive oxygen species (ROS).

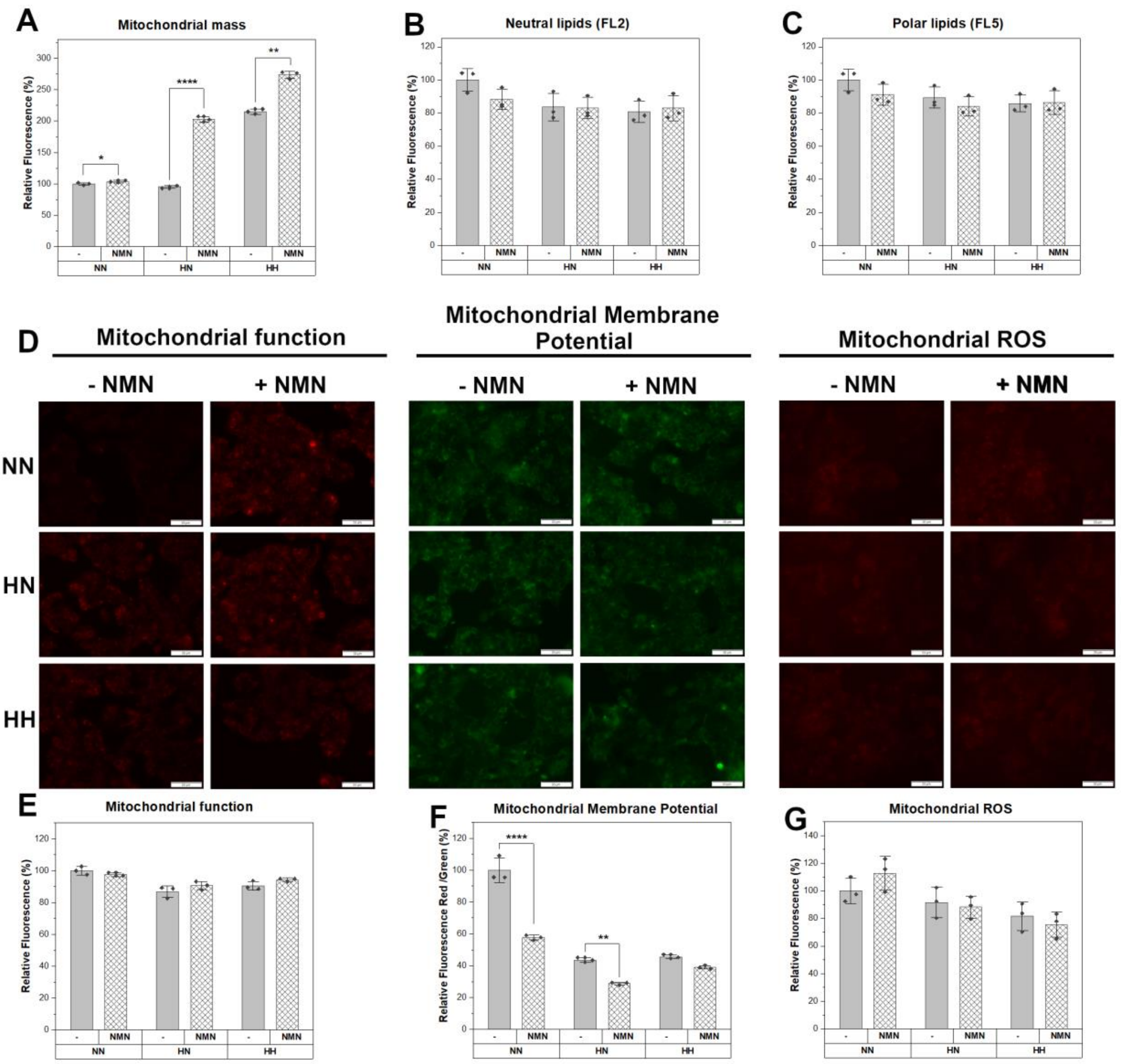


(A) Mitochondrial mass with 10-nonyl-acridine orange (NAO).
(B) Neutral lipids with Nile Red (FL2).
(C) Polar lipids with Nile Red (FL5).
(D) Fluorescence microscopy. Scale bars, 50 µm.
(E) Mitochondrial function with MitoView Red.
(F) Mitochondrial membrane potential with JC-1.
(G) Mitochondrial ROS with DHR-123 in C2C12 myotubes during 100 µM NMN treatment in normoglycemic (NN), hyperglycaemic followed by culture media switch to normoglycemic during NMN treatment (HN), hyperglycaemic before and during treatment (HH), versus untreated condition (-). The flow cytometry data are illustrated as average values of the groups ($n$ = 3 * 15 000 events) ± standard deviation of the mean (STDEV) and statistical significance between NMN treated and untreated condition. *$p$ <0.05; **$p$ <0.01; ****$p$ <0.0001.

**Figure S2.** NMN effects on C2C12 derived myotubes exposed to hyperglycemic conditions: neutral lipids, polar lipids, mitochondrial mass, metabolic activity, membrane potential, and reactive oxygen species (ROS).

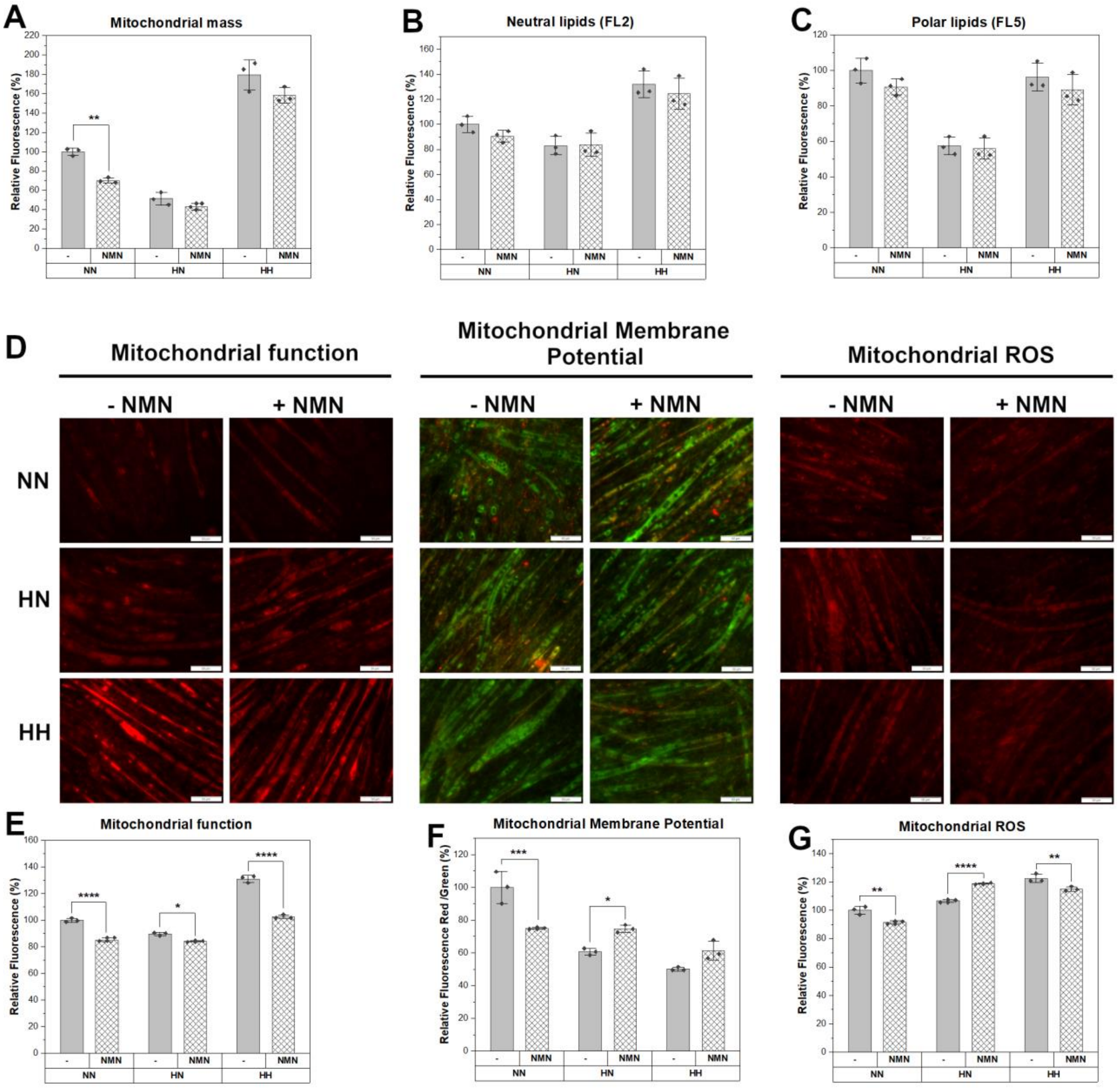


(A) Mitochondrial mass with 10-nonyl-acridine orange (NAO).
(B) Neutral lipids with Nile Red (FL2).
(C) Polar lipids with Nile Red (FL5).
(D) Fluorescence microscopy. Scale bars, 50 µm.
(E) Mitochondrial function with MitoView Red.
(F) Mitochondrial membrane potential with JC-1.
(G) Mitochondrial ROS with DHR-123 in C2C12 myotubes during 100 µM NMN treatment in normoglycemic (NN), hyperglycaemic followed by culture media switch to normoglycemic during NMN treatment (HN), hyperglycaemic before and during treatment (HH), versus untreated condition (-). The flow cytometry data are illustrated as average values of the groups (*n* = 3 * 15 000 events) ± standard deviation of the mean (STDEV) and statistical significance between NMN treated and untreated condition. *$p$ <0.05; **$p$ <0.01; ***$p$ <0.001; ****$p$ <0.0001.

**Figure S3.** Clustered heatmap of the differentially expressed proteins in HepG2 cells.

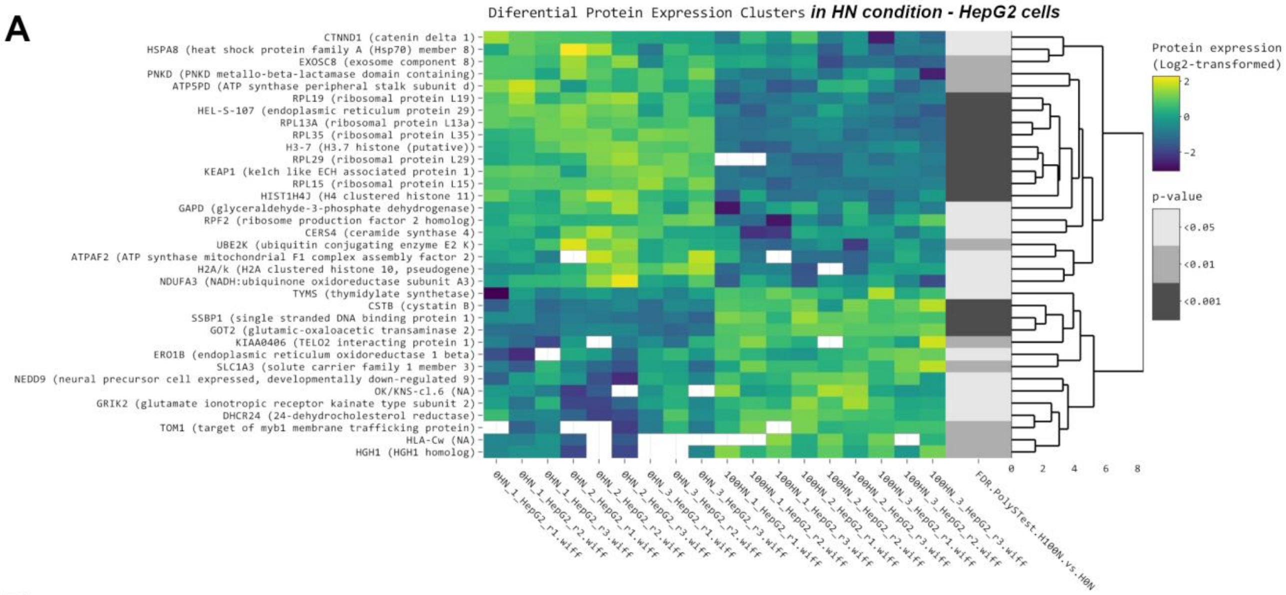


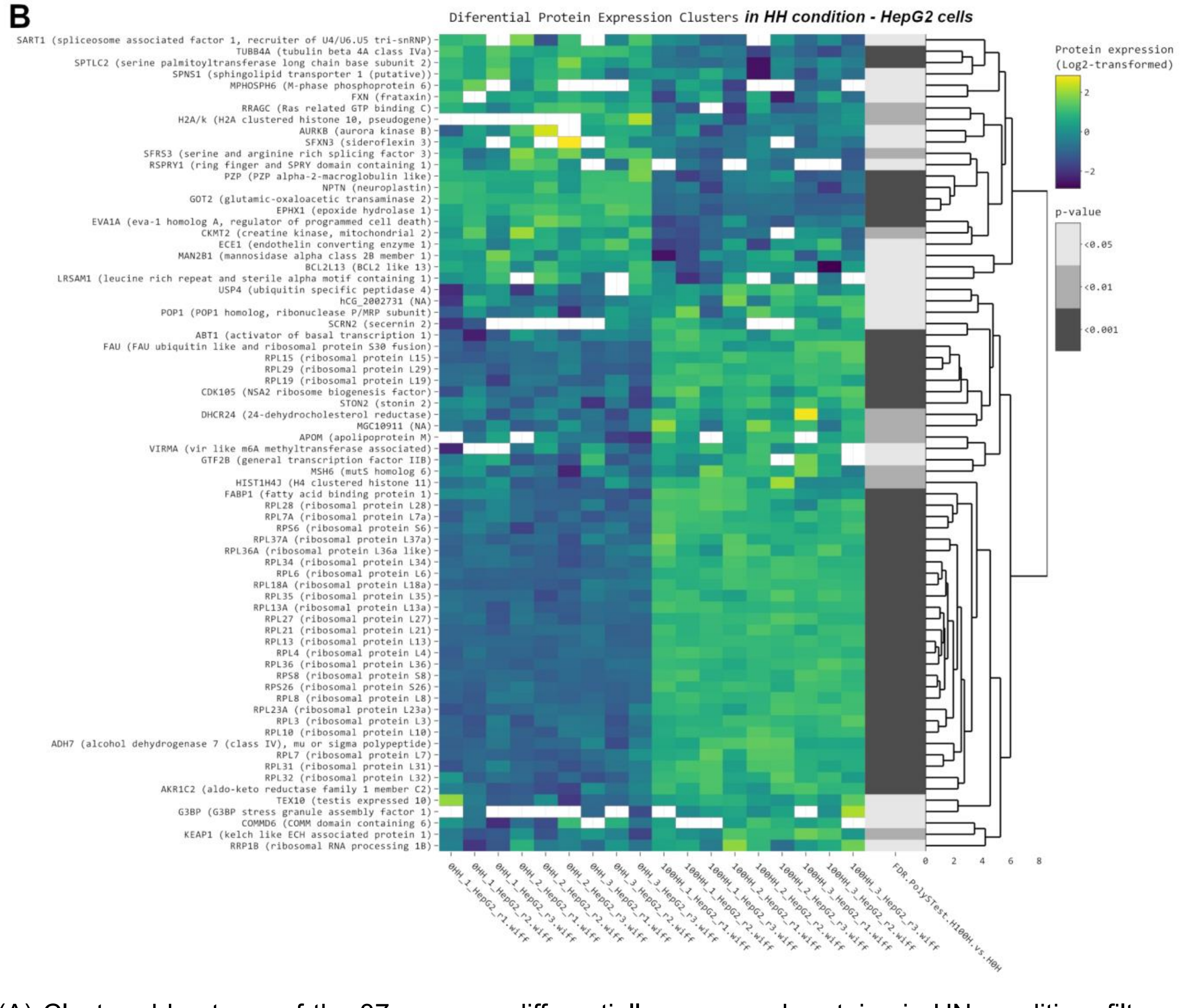


(A) Clustered heatmap of the 37 common differentially expressed proteins in HN condition, filtered with $log_2FC$ threshold set to exclude the interval -0.3, 0.3,0.05 FDR threshold. From left to right, expression values ($log_2$ transformed) for replicates (3 biological x 3 technical) are shown for the 0HN

group and for the 100HN (treated) group, followed by significance values of the comparison to 0HN group.
(B) Clustered heatmap of the 85 differently expressed proteins in HH condition, filtered with $log_2FC$ threshold set to exclude the interval -0.3, 0.3 and 0.05 FDR threshold. From left to right, expression values ($log_2$ transformed) for replicates (3 biological x 3 technical) are shown for the 0HH (untreated) group and for the 100HH (treated) group, followed by significance values of the comparison to 0HH group.

**Figure S4.** Integrated proteomics data analysis of HepG2 cell line under HN condition.

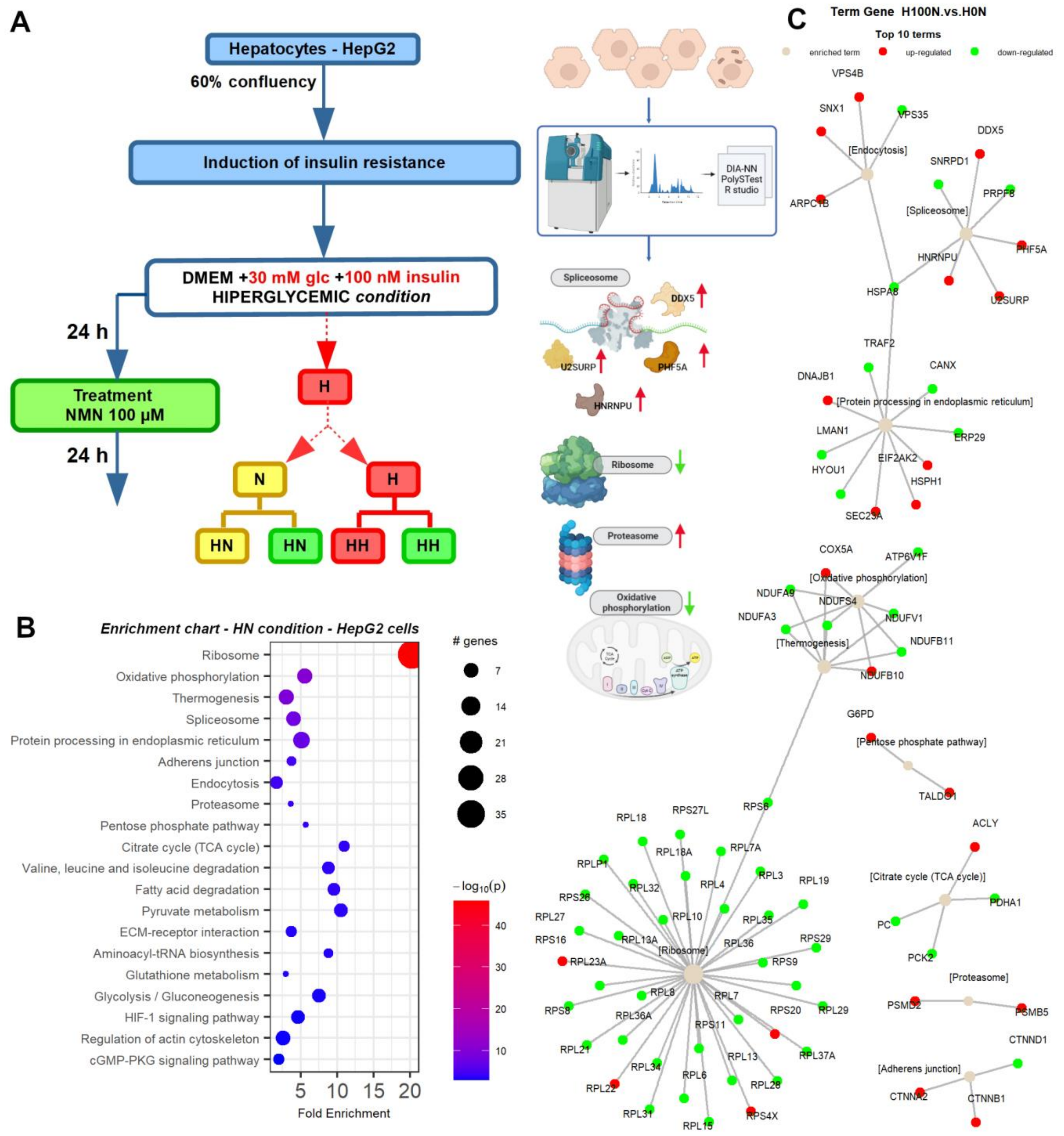


(A) Experiment summary (OpenOffice, BioRender).
(B) Enrichment chart for top 20 KEGG Pathways in HN condition for HepG2 cells sorted by lowest *p* value.
(C) Term-gene graph for top 10 terms in HN condition for HepG2 cells.

**Figure S5.** Integrated proteomics data analysis of HepG2 cell line under HH condition.

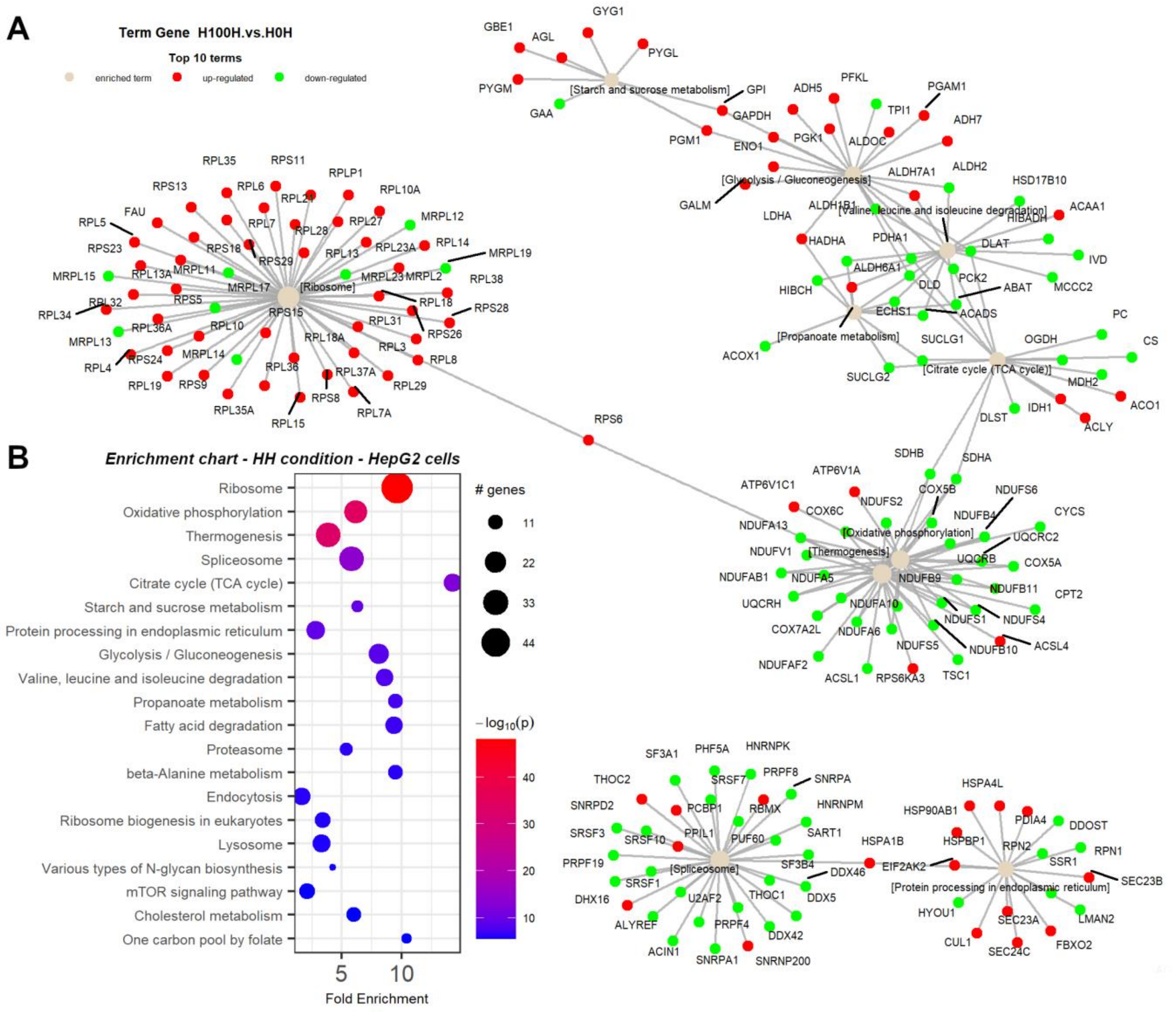


(A) Term-gene graph for top 10 terms in HH condition for HepG2 cells.
(B) Enrichment chart for top 20 KEGG Pathways in HH condition for HepG2 cells sorted by lowest *p* value.

**Figure S6.** Clustered heatmap of the differentially expressed proteins in myotubes.

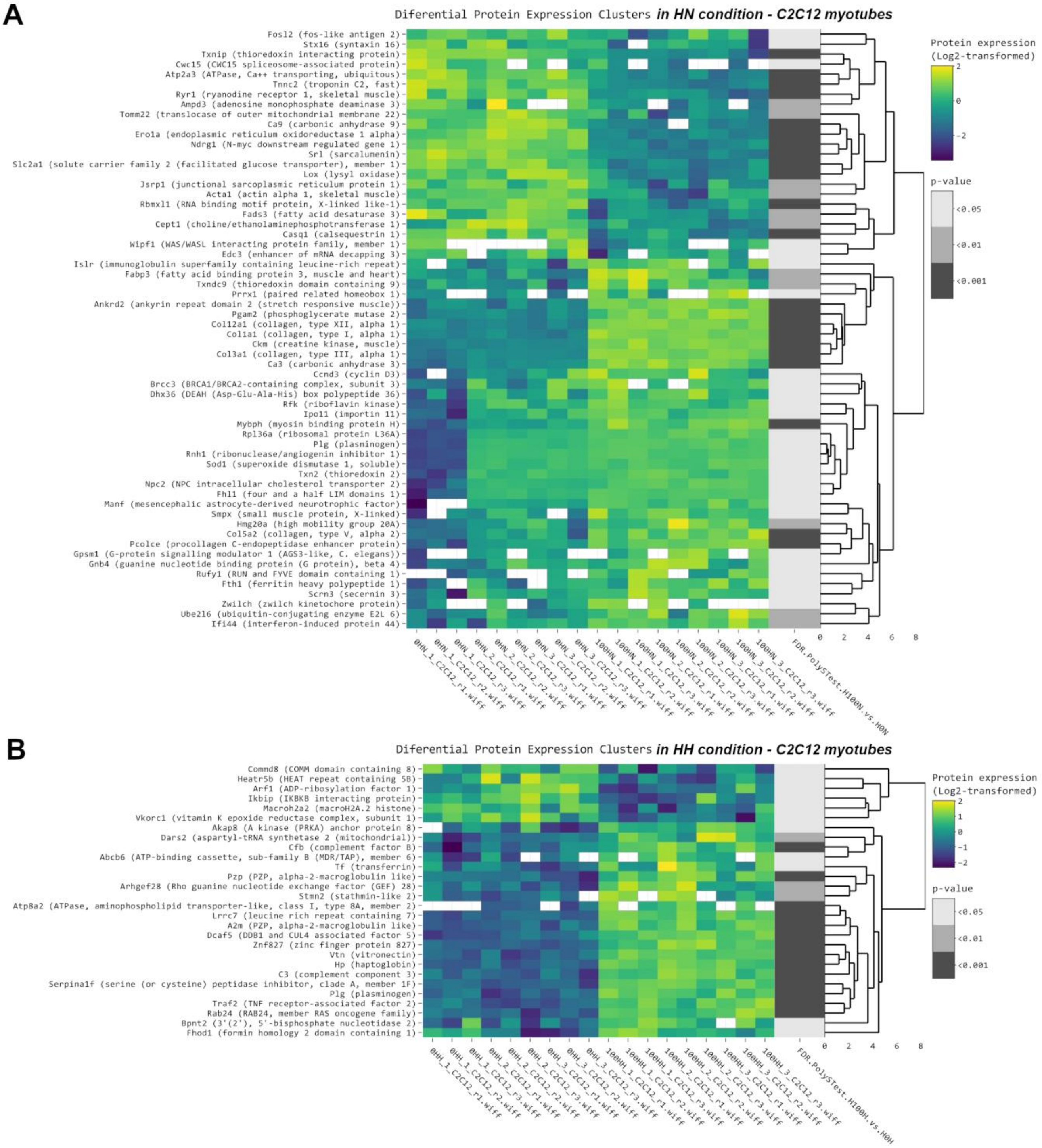


(A) Clustered heatmap of the 63 differentially expressed proteins in HN condition, filtered with $log_2FC$ threshold set to exclude the interval -0.3, 0.3 and 0.05 FDR threshold. From left to right, expression values ($log_2$ transformed) for replicates (3 biological x 3 technical) are shown for the 0HN (untreated) group and for the 100HN (treated) group, followed by significance values of the comparison to 0HN (untreated) group.

(B) Clustered heatmap of the 30 common differentially expressed proteins in HH condition, filtered with $log_2FC$ threshold set to exclude the interval -0.3, 0.3 and 0.05 FDR threshold. From left to right, expression values ($log_2$ transformed) for replicates (3 biological x 3 technical) are shown for the 0HH group and for the 100HH (treated) group, followed by significance values of the comparison to 0HH (untreated) group.

**Figure S7.** Integrated proteomics data analysis of myotubes under HN condition.

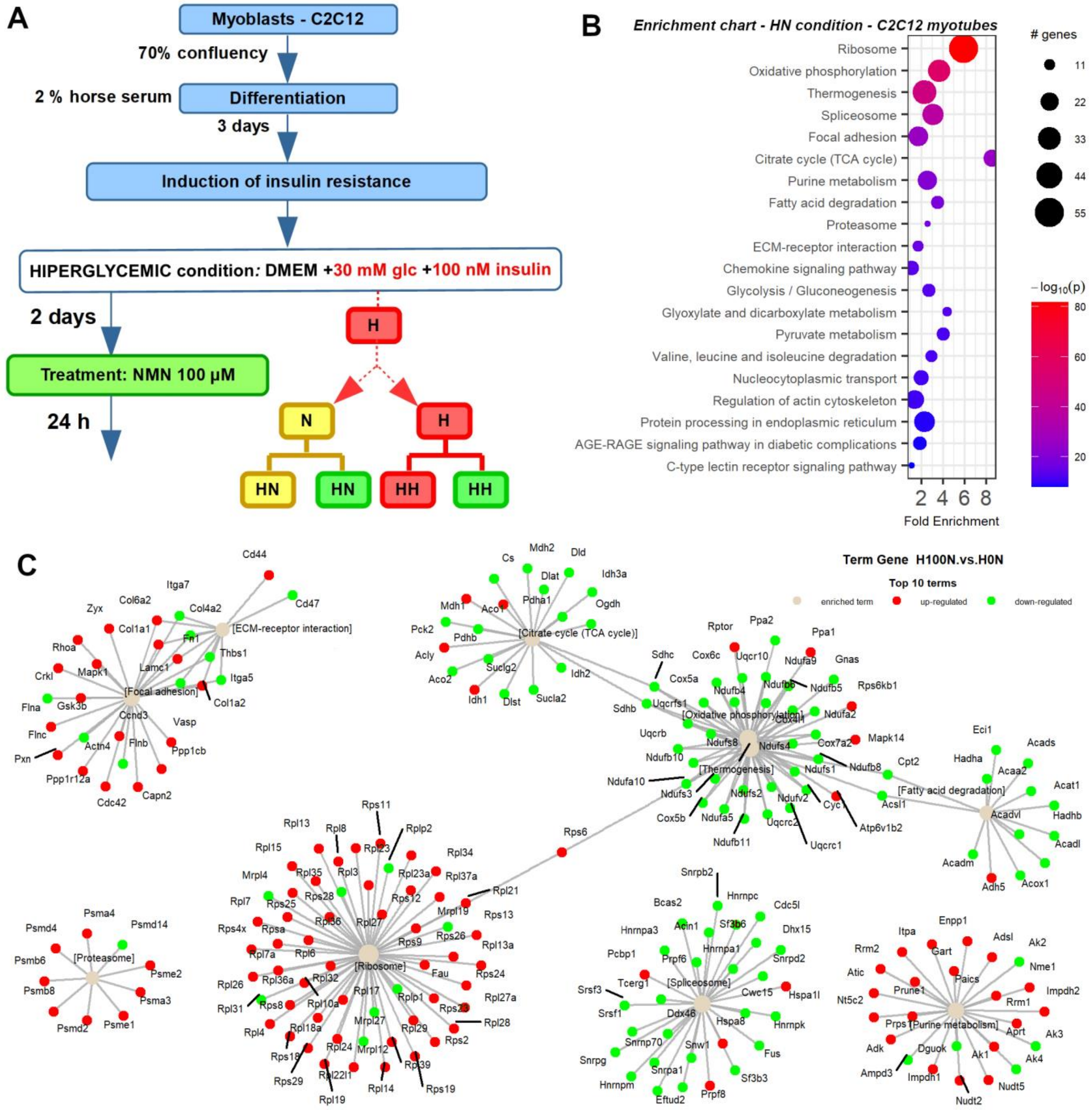


(A) Experiment summary (OpenOffice) in myotubes.
(B) Enrichment chart for top 20 KEGG pathways in HN condition for myotubes cells sorted by lowest *p* value.
(C) Term-gene graph for top 10 terms in HN condition for C2C12 derived myotubes.

**Figure S8.** Integrated proteomics data analysis of myotubes under HH condition.

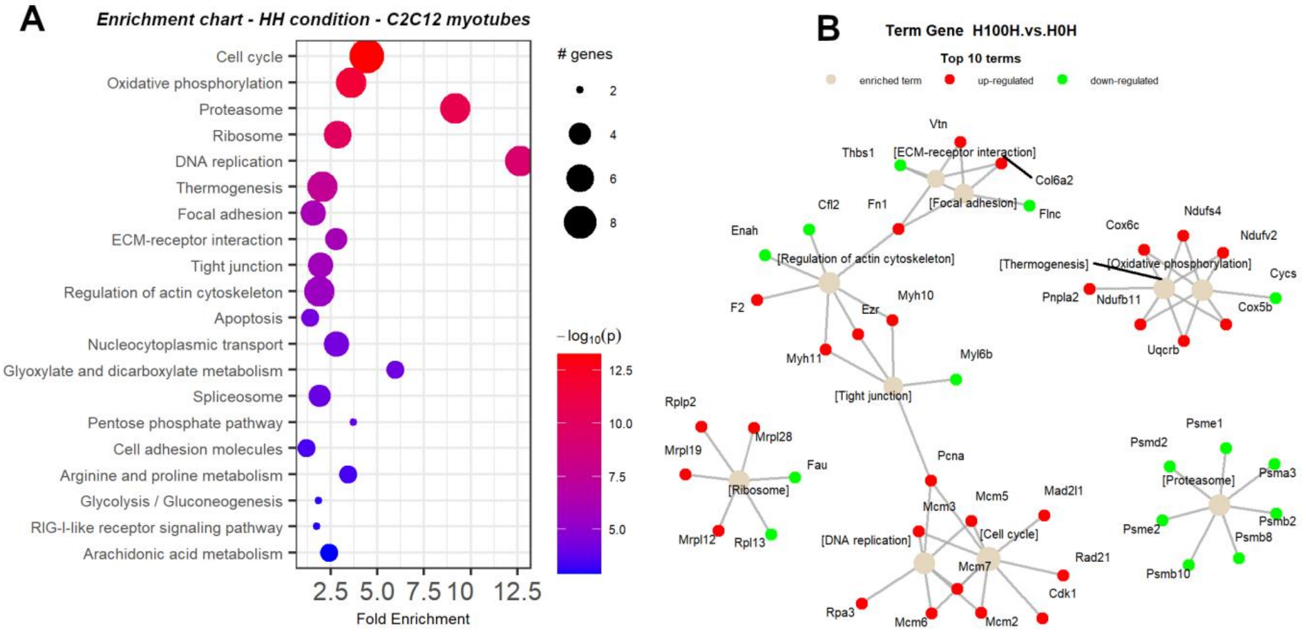


(A) Enrichment chart for top 20 KEGG Pathways in HH condition for C2C12 derived myotubes sorted by lowest *p* value.
(B) Term-gene graph for top 10 terms in HH condition for C2C12 myotubes.
