## Supplementary material for "NMN Works in HFD-Induced T2DM by Interesting Effects in Adipose Tissue, and Not by Mitochondrial Biogenesis": Document_S2.pdf

**Figure S9.** NMN treatment effects on the KEGG steroid hormone biosynthesis pathway in mouse liver. The colour of the boxes represents the log2 fold change of the protein abundances, represented for HFD+NMN group versus HFD group comparison. Red: up-regulated; green: down-regulated; grey: no significant expression change.

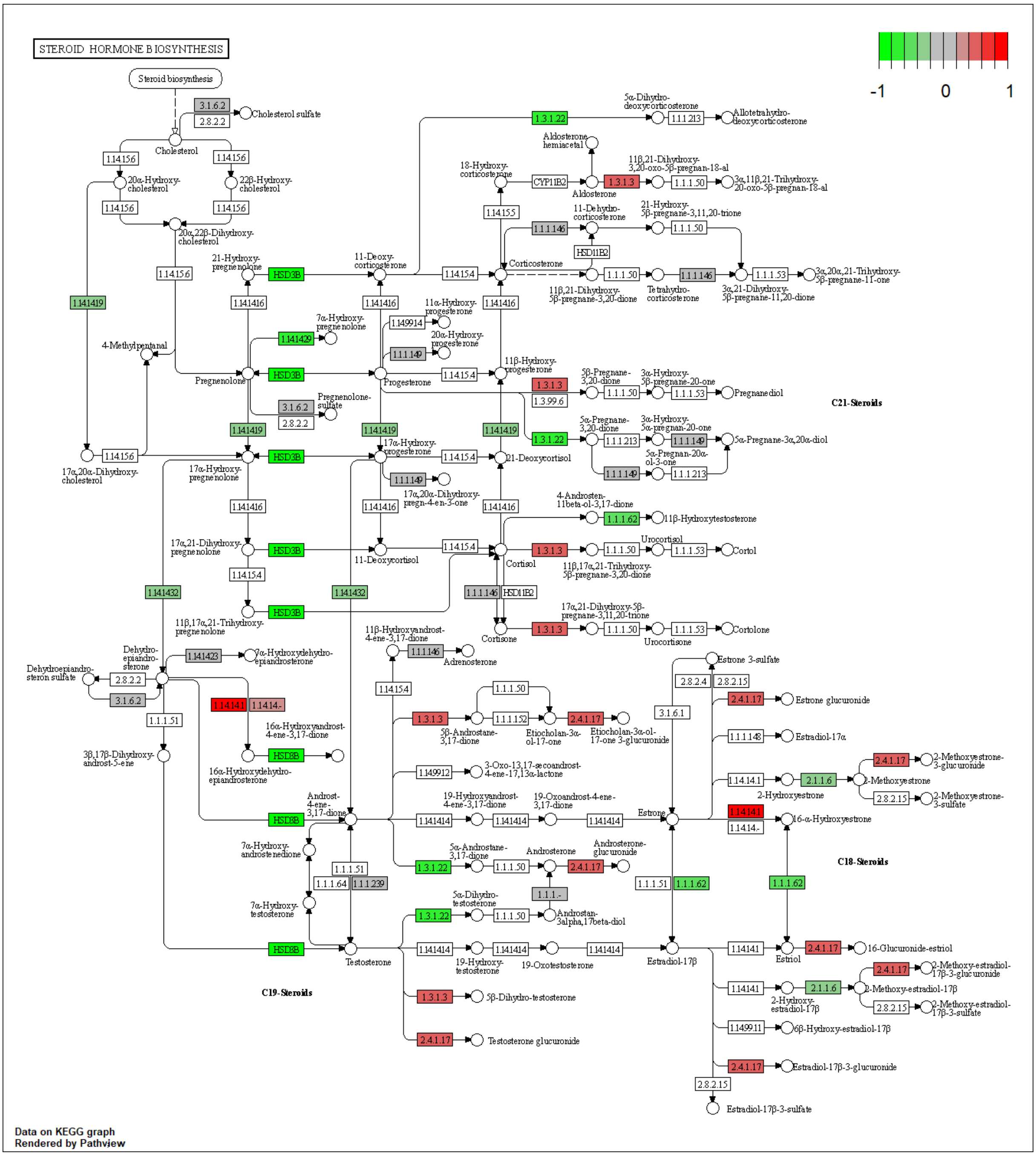

**Figure S10.** NMN treatment effects on the KEGG ribosome pathway in mouse liver. The colour of the boxes represents the log2 fold change of the protein abundances, represented for HFD+NMN group versus HFD group comparison. Red: up-regulated; green: down-regulated; grey: no significant expression change.

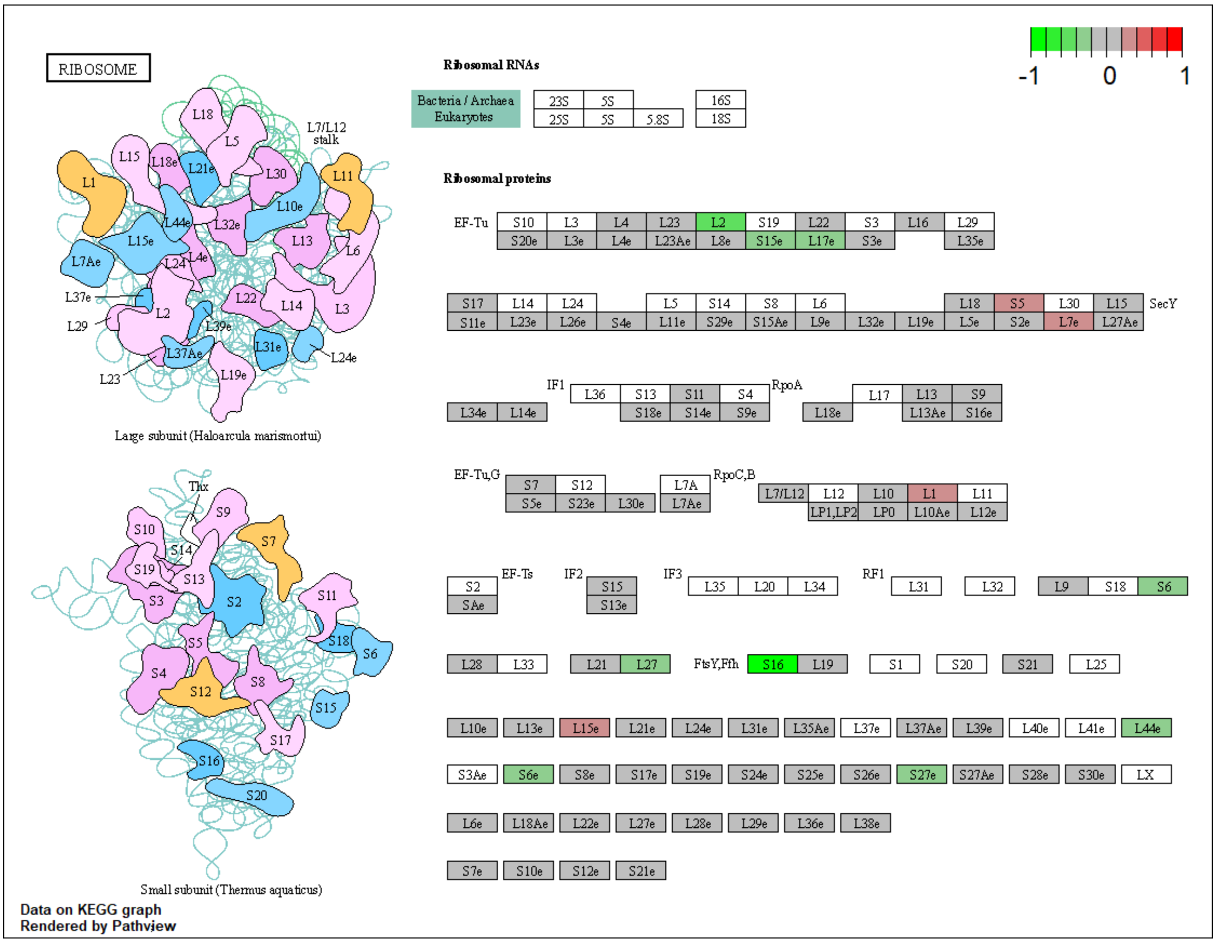


**Figure S11.** NMN treatment effects on the KEGG arachidonic acid metabolism pathway in mouse liver. The colour of the boxes represents the log2 fold change of the protein abundances, represented for HFD+NMN group versus HFD group comparison. Red: up-regulated; green: down-regulated; grey: no significant expression change.

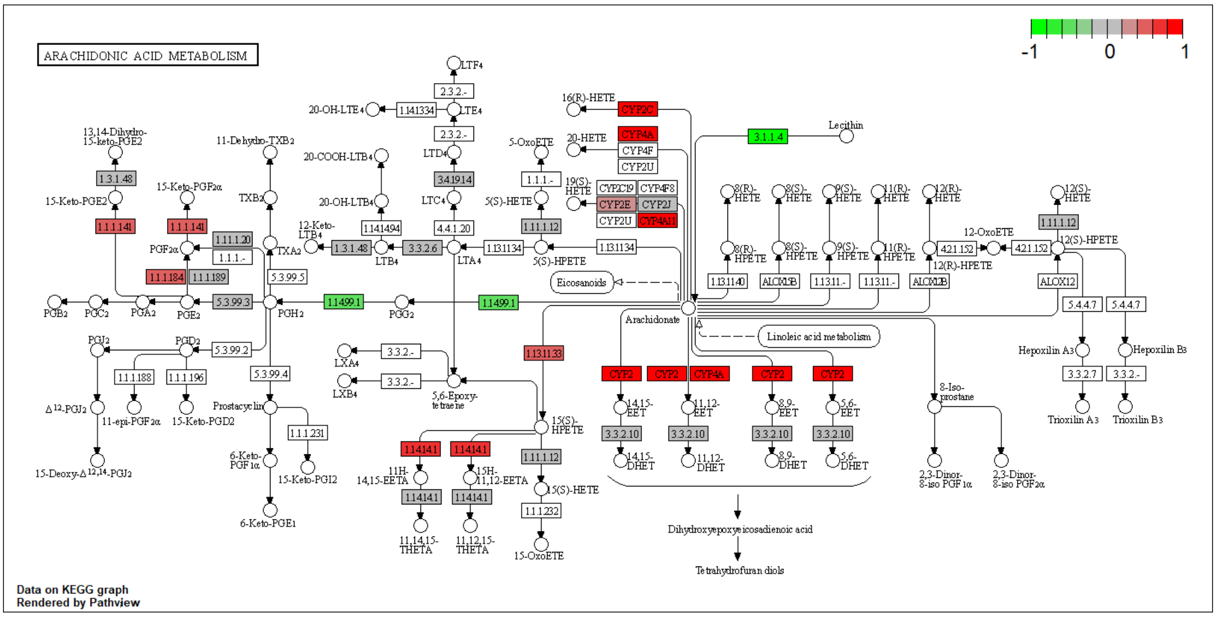

**Figure S12.** NMN treatment effects on the KEGG ribosome biogenesis pathway in mouse liver. The colour of the boxes represents the log2 fold change of the protein abundances, represented for HFD+NMN group versus HFD group comparison. Red: up-regulated; green: down-regulated; grey: no significant expression change.

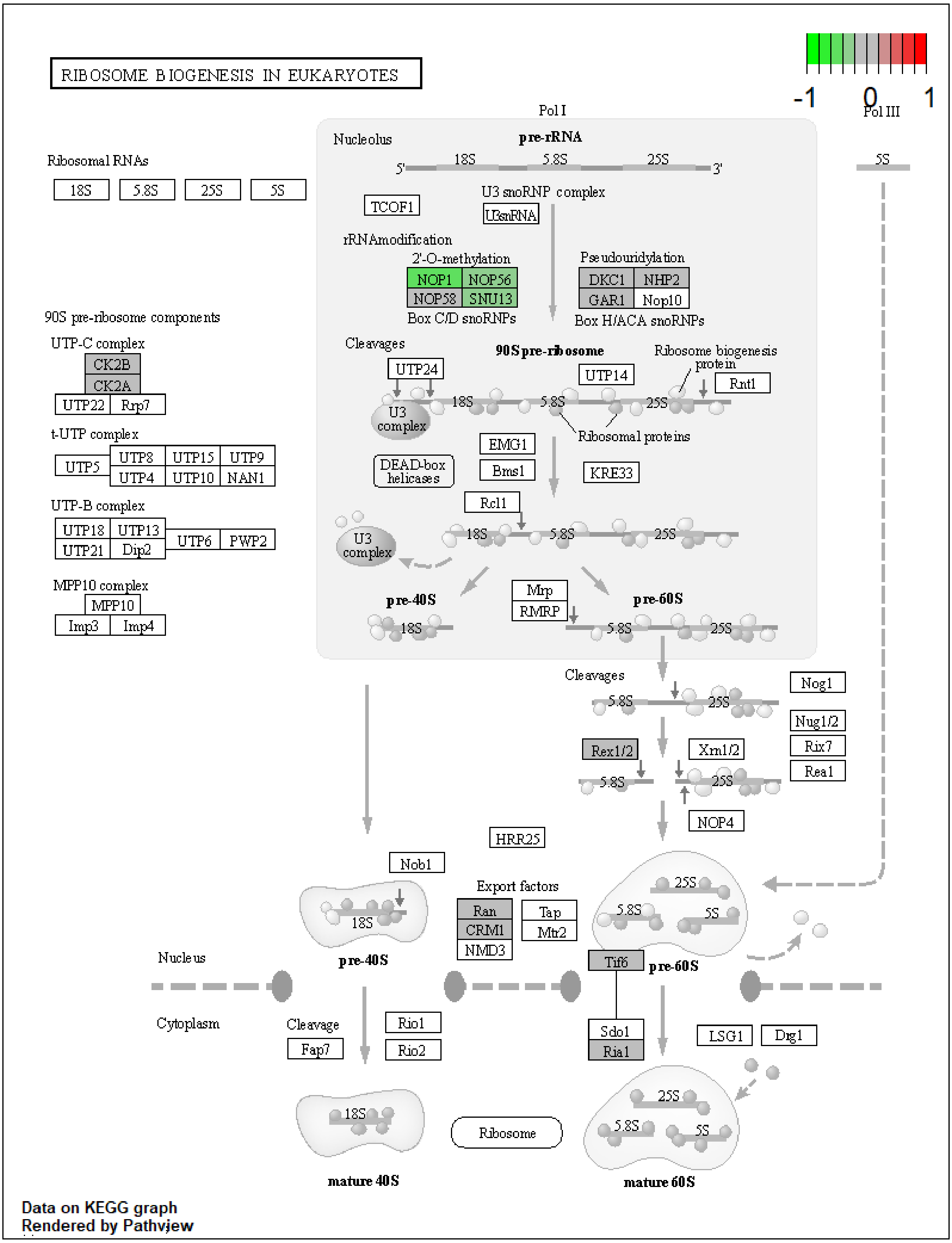

**Figure S13.** NMN treatment effects on the KEGG metabolism of xenobiotics by cytochrome P450 pathway in mouse liver. The colour of the boxes represents the log2 fold change of the protein abundances, represented for HFD+NMN group versus HFD group comparison. Red: up-regulated; green: down-regulated; grey: no significant expression change.

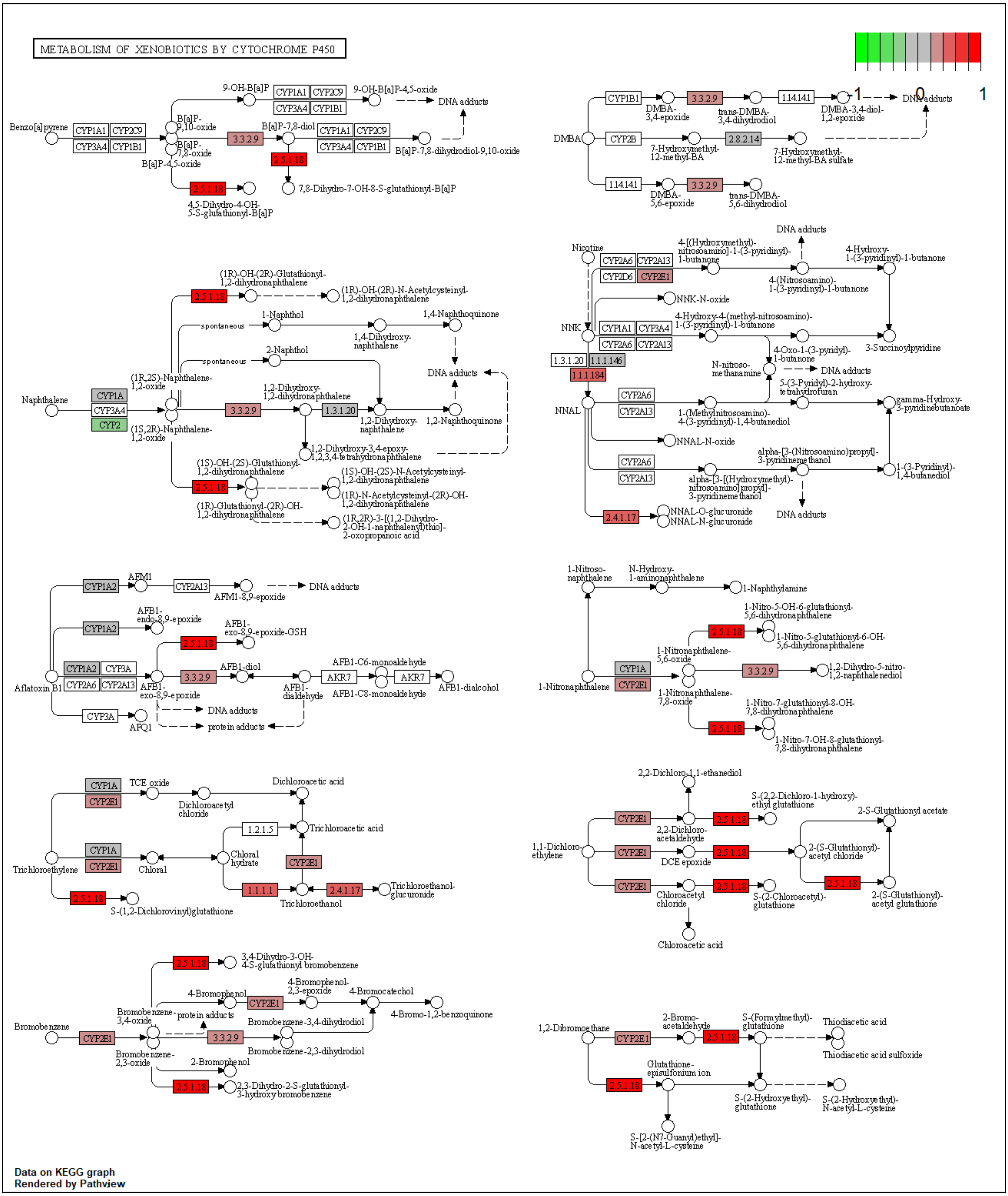

**Figure S14.** NMN treatment effects on the KEGG metabolism of spliceosome pathway in mouse liver. The colour of the boxes represents the log2 fold change of the protein abundances, represented for HFD+NMN group versus HFD group comparison. Red: up-regulated; green: down-regulated; grey: no significant expression change.

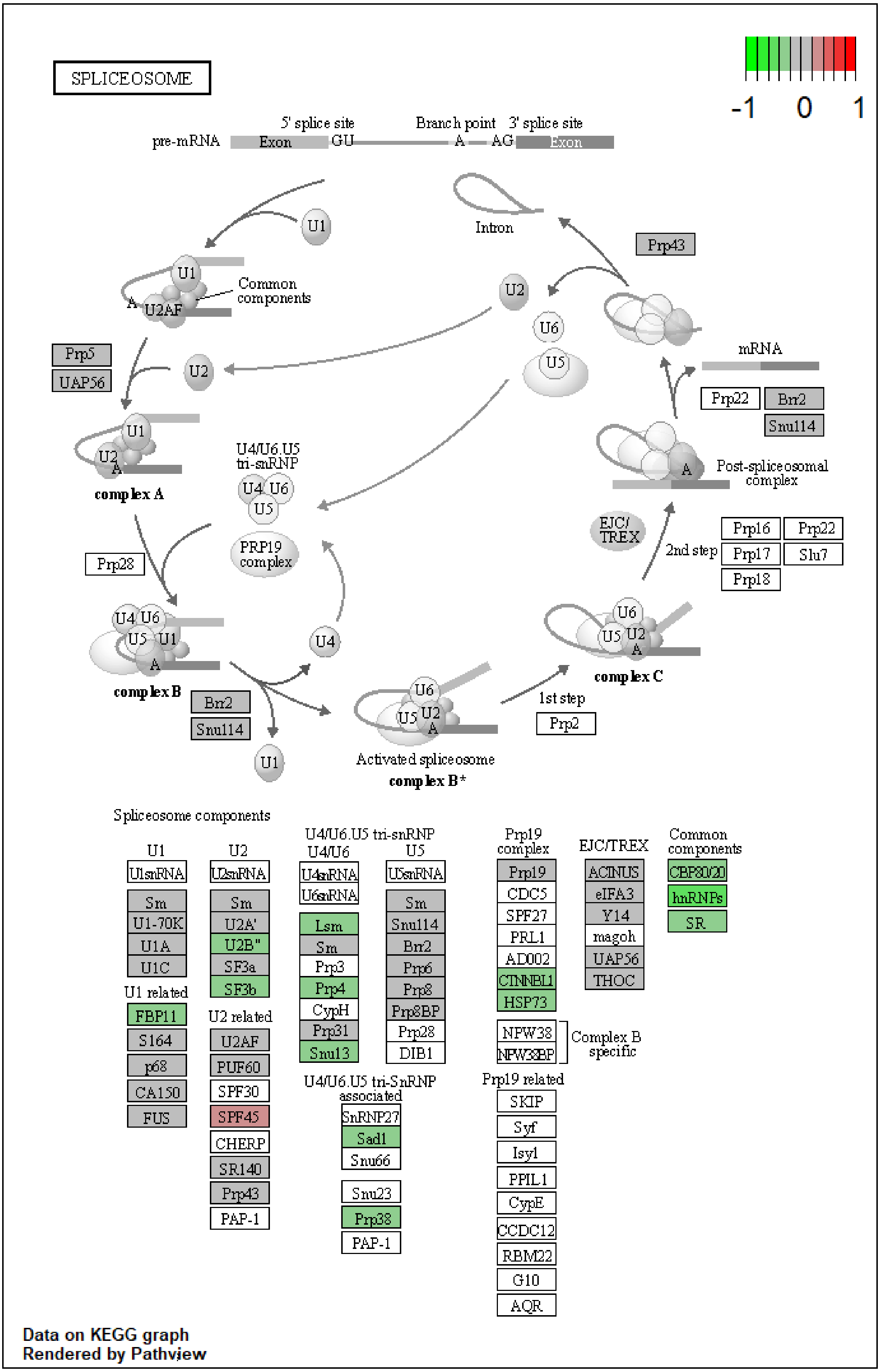

**Figure S15.** NMN treatment effects on the KEGG glycolysis /gluconeogenesis pathway in mouse muscle. The colour of the boxes represents the log2 fold change of the protein abundances, represented for HFD+NMN group versus HFD group comparison. Red: up-regulated; green: down-regulated; grey: no significant expression change.

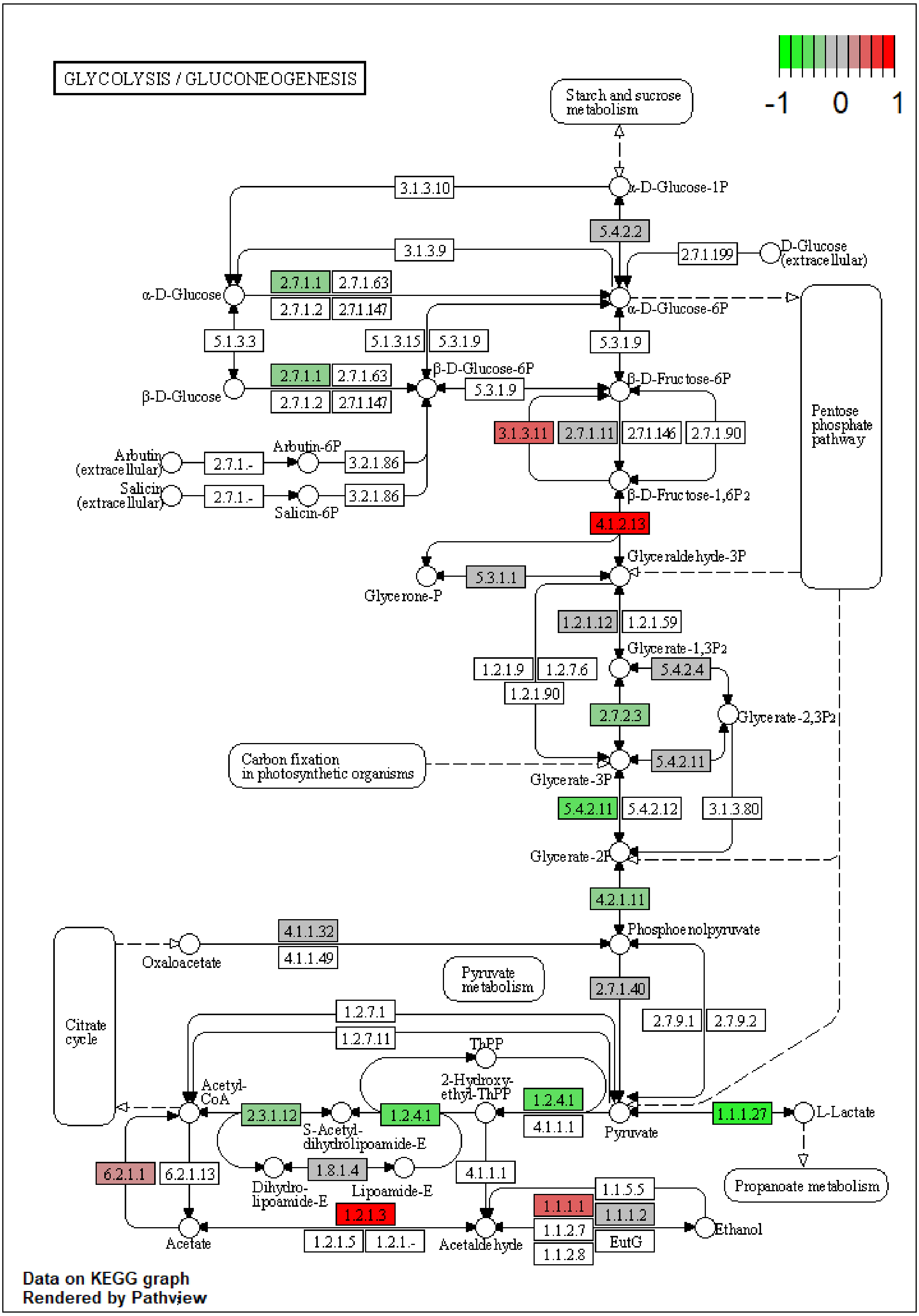

**Figure S16.** NMN treatment effects on the KEGG **g**lycine /serine /threonine metabolism pathway in mouse muscle. The colour of the boxes represents the log2 fold change of the protein abundances, represented for HFD+NMN group versus HFD group comparison. Red: up-regulated; green: down-regulated; grey: no significant expression change.

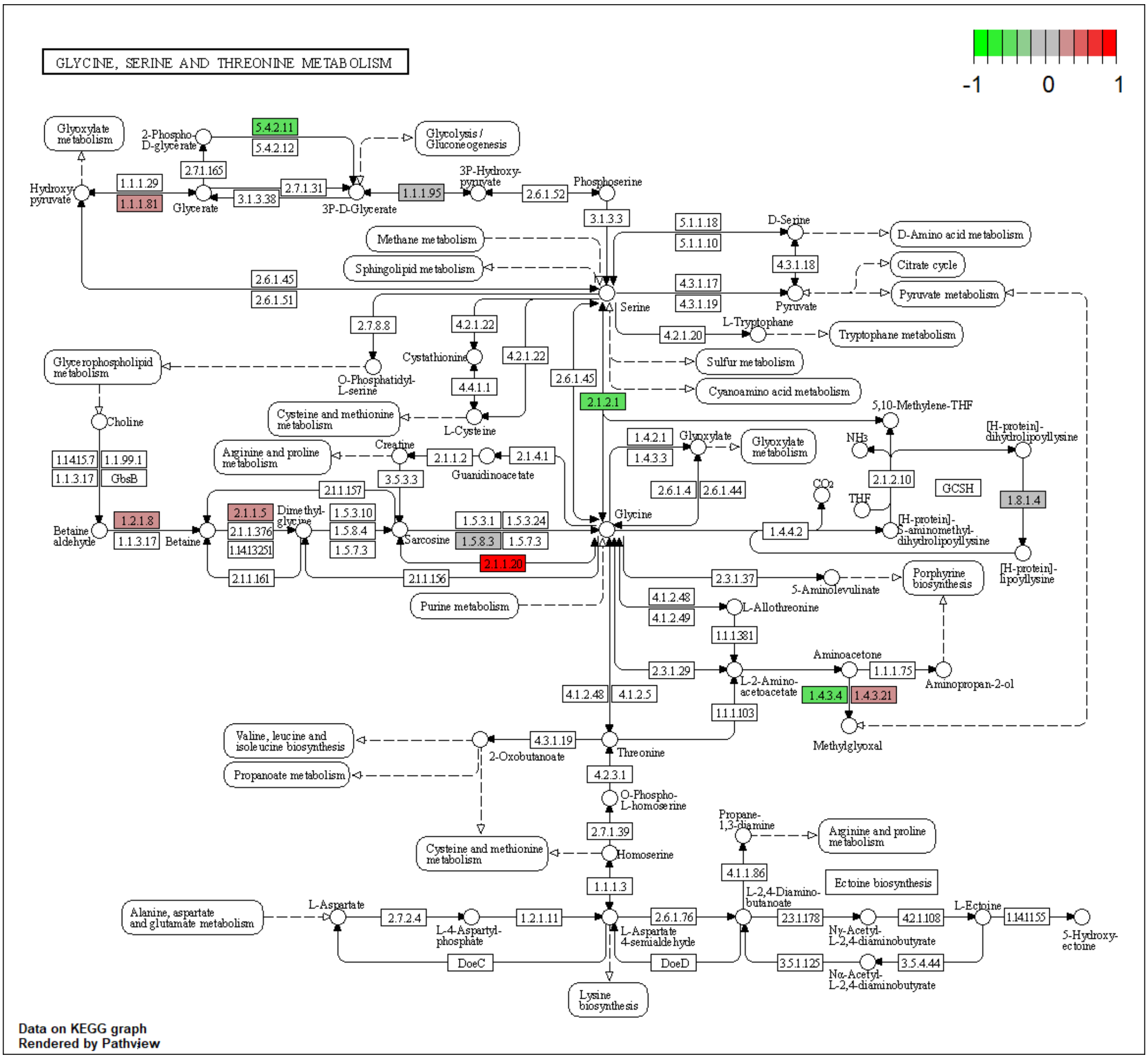

**Figure S17.** NMN treatment effects on the KEGG pyruvate metabolism pathway in mouse muscle. The colour of the boxes represents the log2 fold change of the protein abundances, represented for HFD+NMN group versus HFD group comparison. Red: up-regulated; green: down-regulated; grey: no significant expression change.

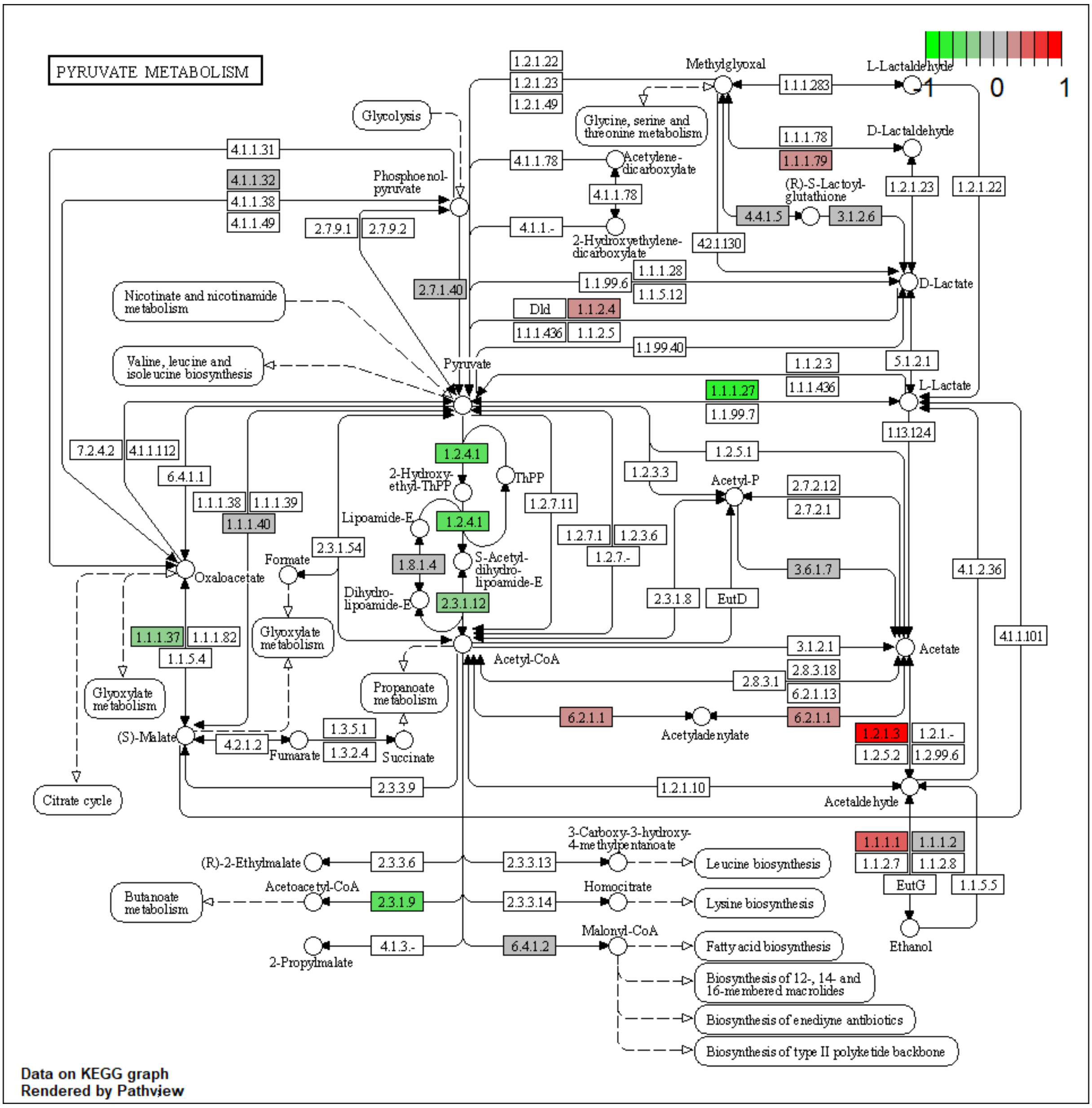

**Figure S18.** NMN treatment effects on the KEGG valine, leucine and isoleucine degradation pathway in mouse muscle. The colour of the boxes represents the log2 fold change of the protein abundances, represented for HFD+NMN group versus HFD group comparison. Red: up-regulated; green: down-regulated; grey: no significant expression change.

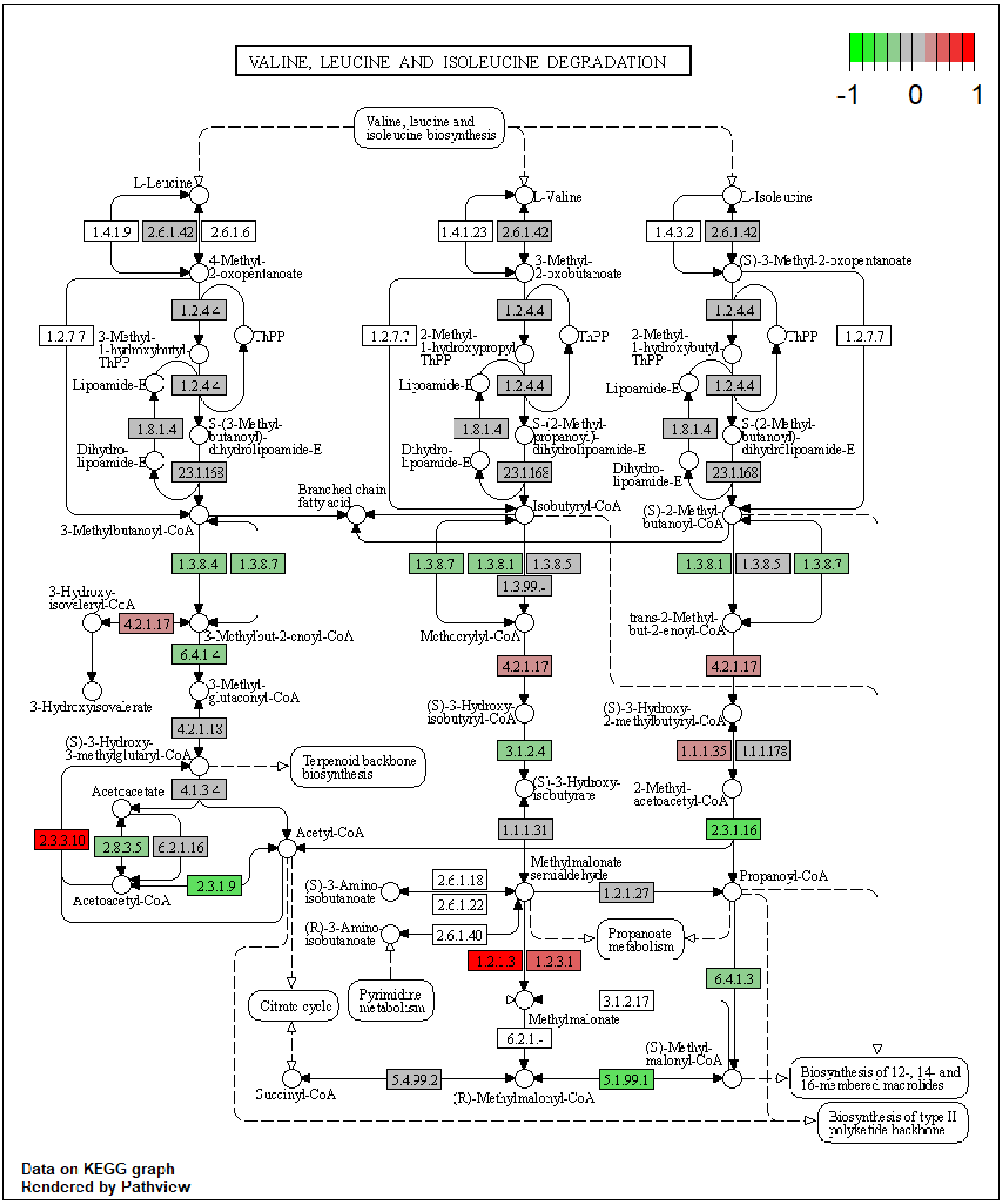

**Figure S19.** NMN treatment effects on the KEGG spliceosome pathway in mouse adipose tissue. The colour of the boxes represents the log2 fold change of the protein abundances, represented for HFD+NMN group versus HFD group comparison. Red: up-regulated; green: down-regulated; grey: no significant expression change.

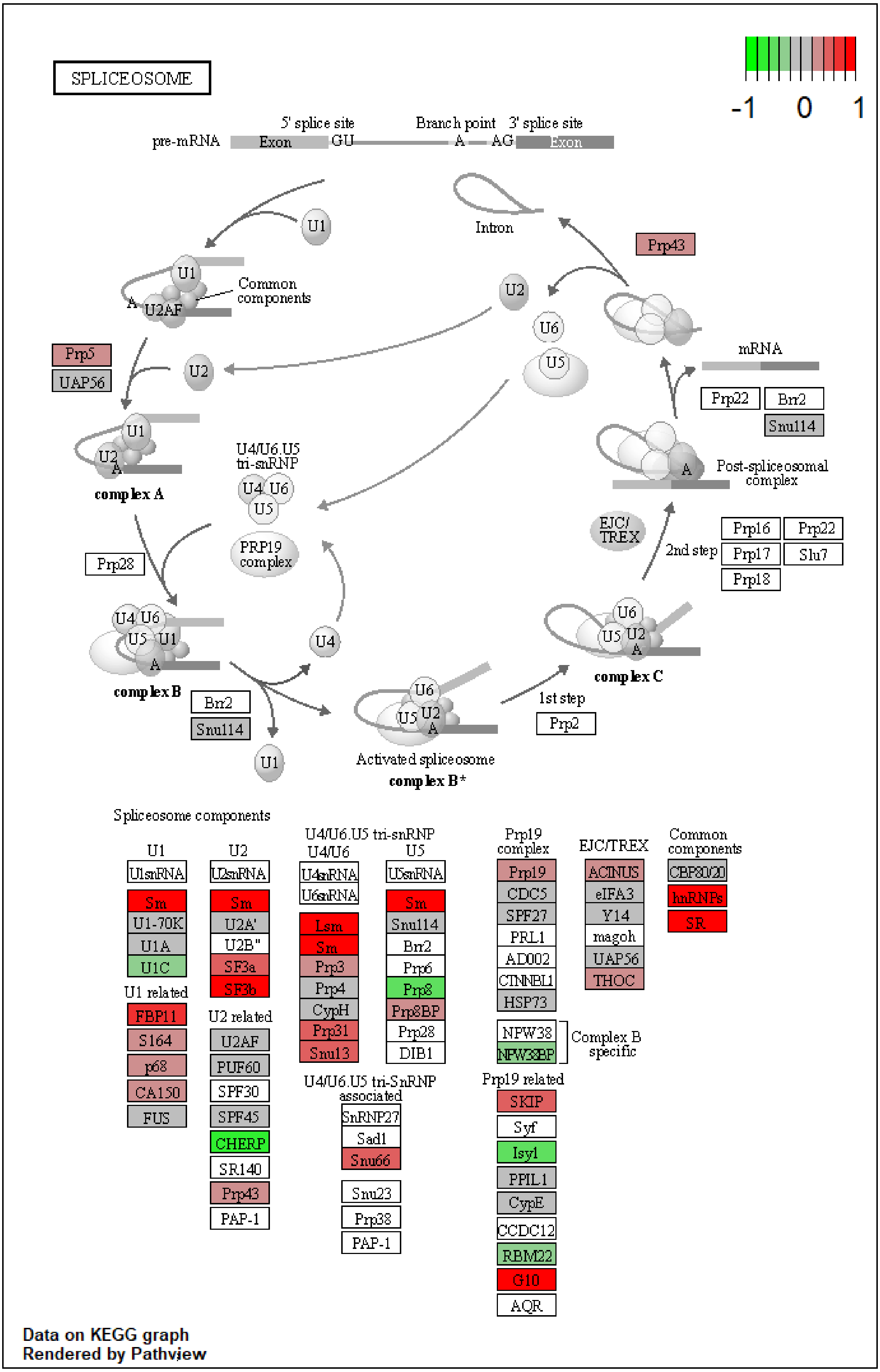

**Figure S20.** NMN treatment effects on the KEGG regulation of actin cytoskeleton pathway in mouse adipose tissue. The colour of the boxes represents the log2 fold change of the protein abundances, represented for HFD+NMN group versus HFD group comparison. Red: up-regulated; green: down-regulated; grey: no significant expression change.

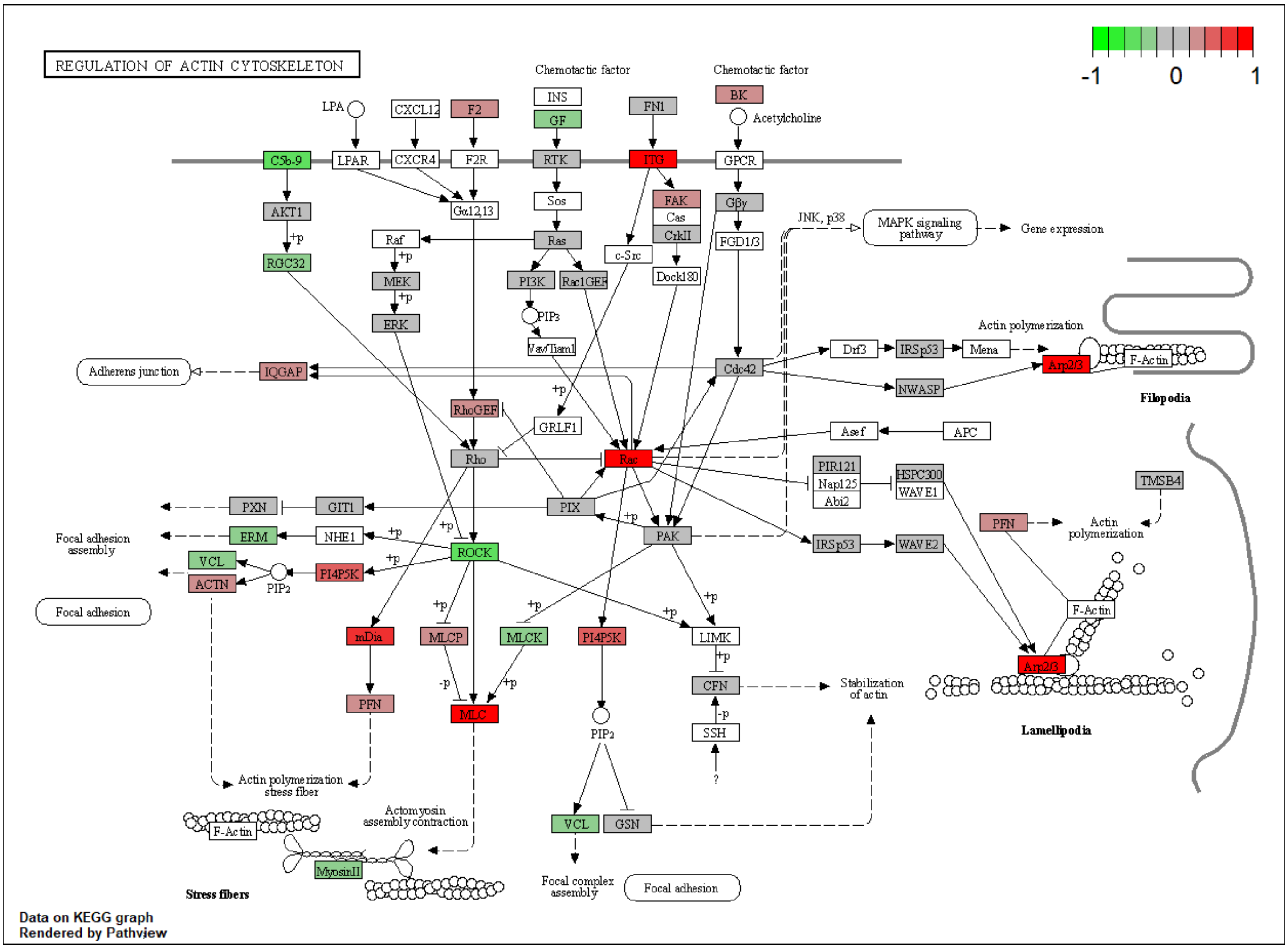

**Figure S21.** NMN treatment effects on the KEGG proteasome pathway in mouse adipose tissue. The colour of the boxes represents the log2 fold change of the protein abundances, represented for HFD+NMN group versus HFD group comparison. Red: up-regulated; green: down-regulated; grey: no significant expression change.

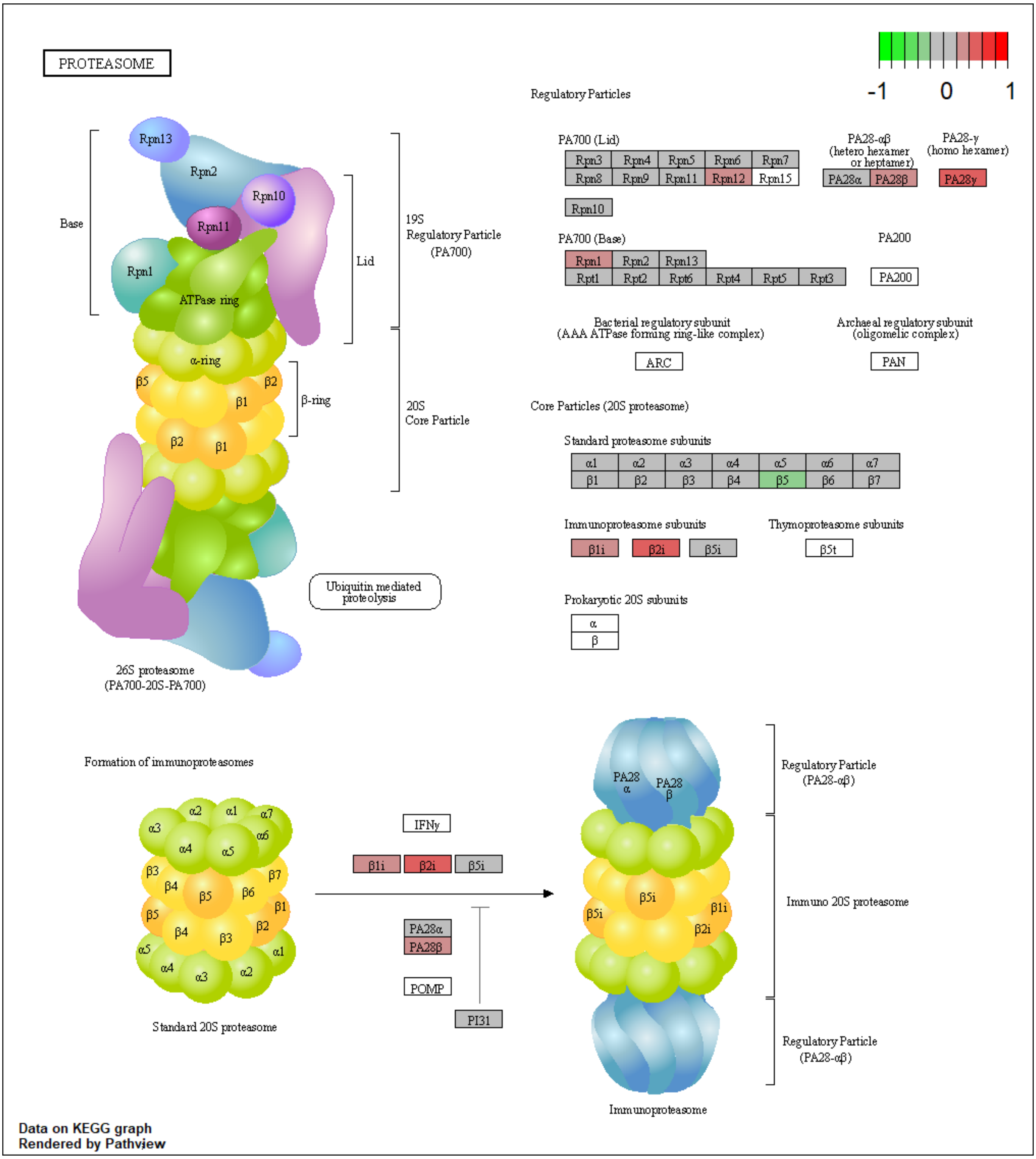

**Figure S22.** NMN treatment effects on the KEGG mTOR signaling pathway in mouse adipose tissue. The colour of the boxes represents the log2 fold change of the protein abundances, represented for HFD+NMN group versus HFD group comparison. Red: up-regulated; green: down-regulated; grey: no significant expression change.

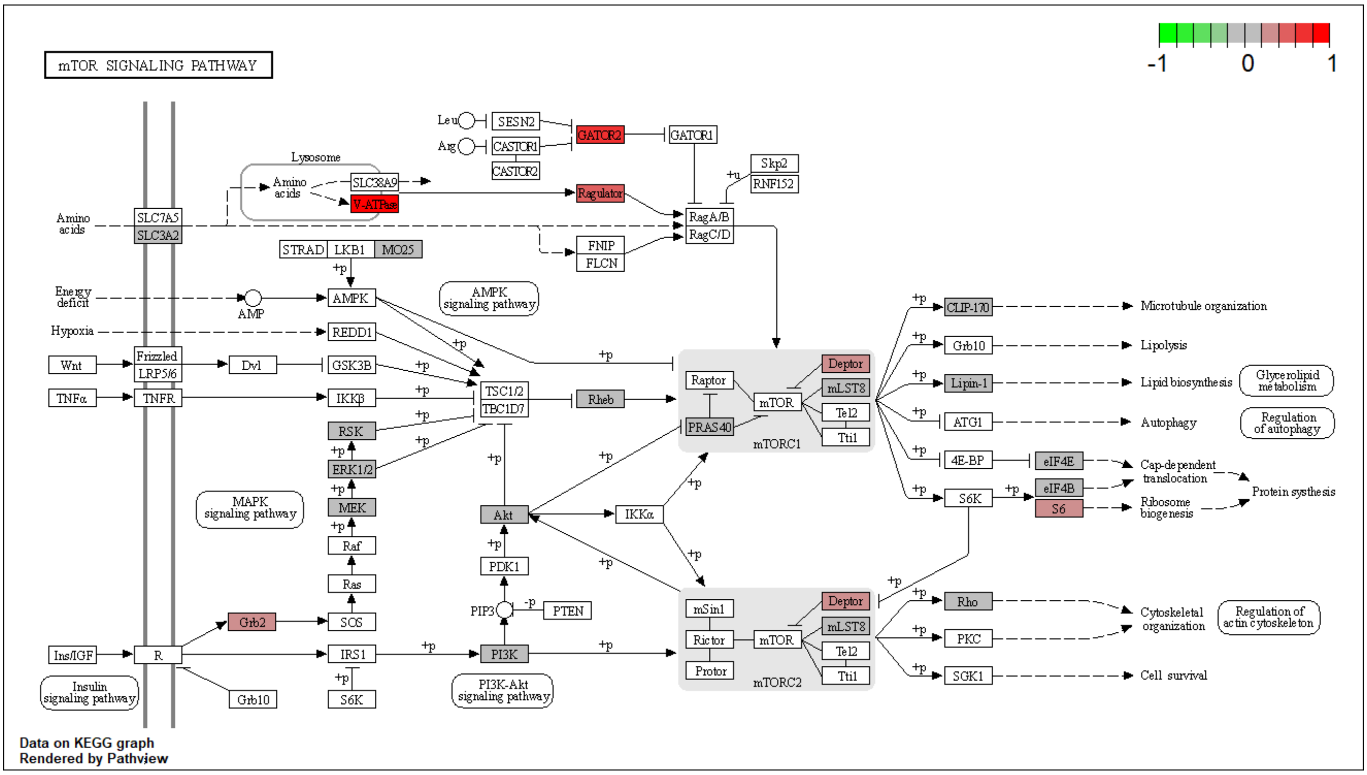

**Figure S23.** NMN treatment effects on the KEGG tight junction's pathway in mouse adipose tissue. The colour of the boxes represents the log2 fold change of the protein abundances, represented for HFD+NMN group versus HFD group comparison. Red: up-regulated; green: down-regulated; grey: no significant expression change.

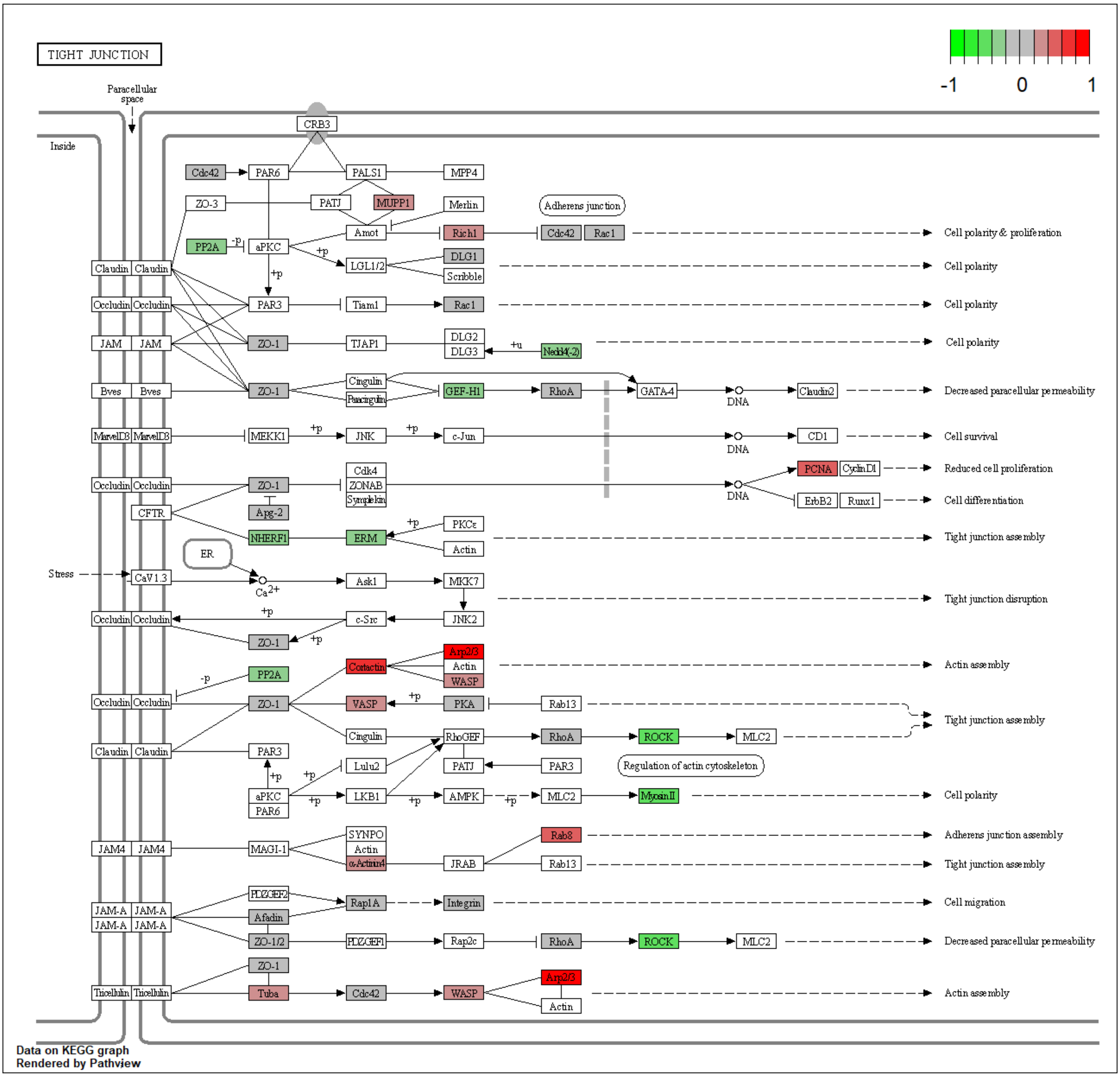

**Figure S24.** NMN treatment effects on the KEGG oxidative phosphorylation pathway in mouse brain. The colour of the boxes represents the log2 fold change of the protein abundances, represented for HFD+NMN group versus HFD group comparison. Red: up-regulated; green: down-regulated; grey: no significant expression change.

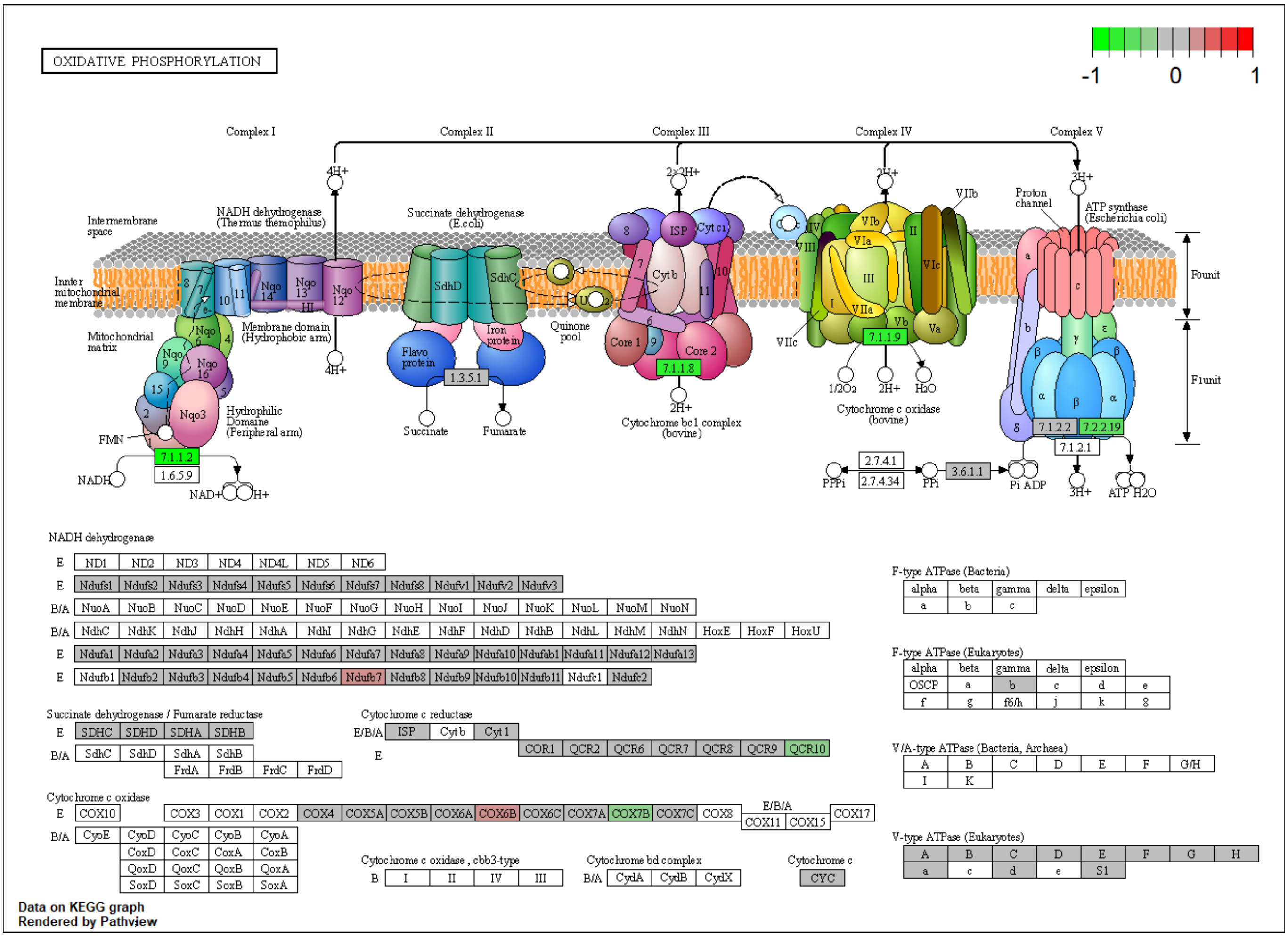

**Figure S25.** NMN treatment effects on the KEGG tight junctions' pathway in mouse brain. The colour of the boxes represents the log2 fold change of the protein abundances, represented for HFD+NMN group versus HFD group comparison. Red: up-regulated; green: down-regulated; grey: no significant expression change.

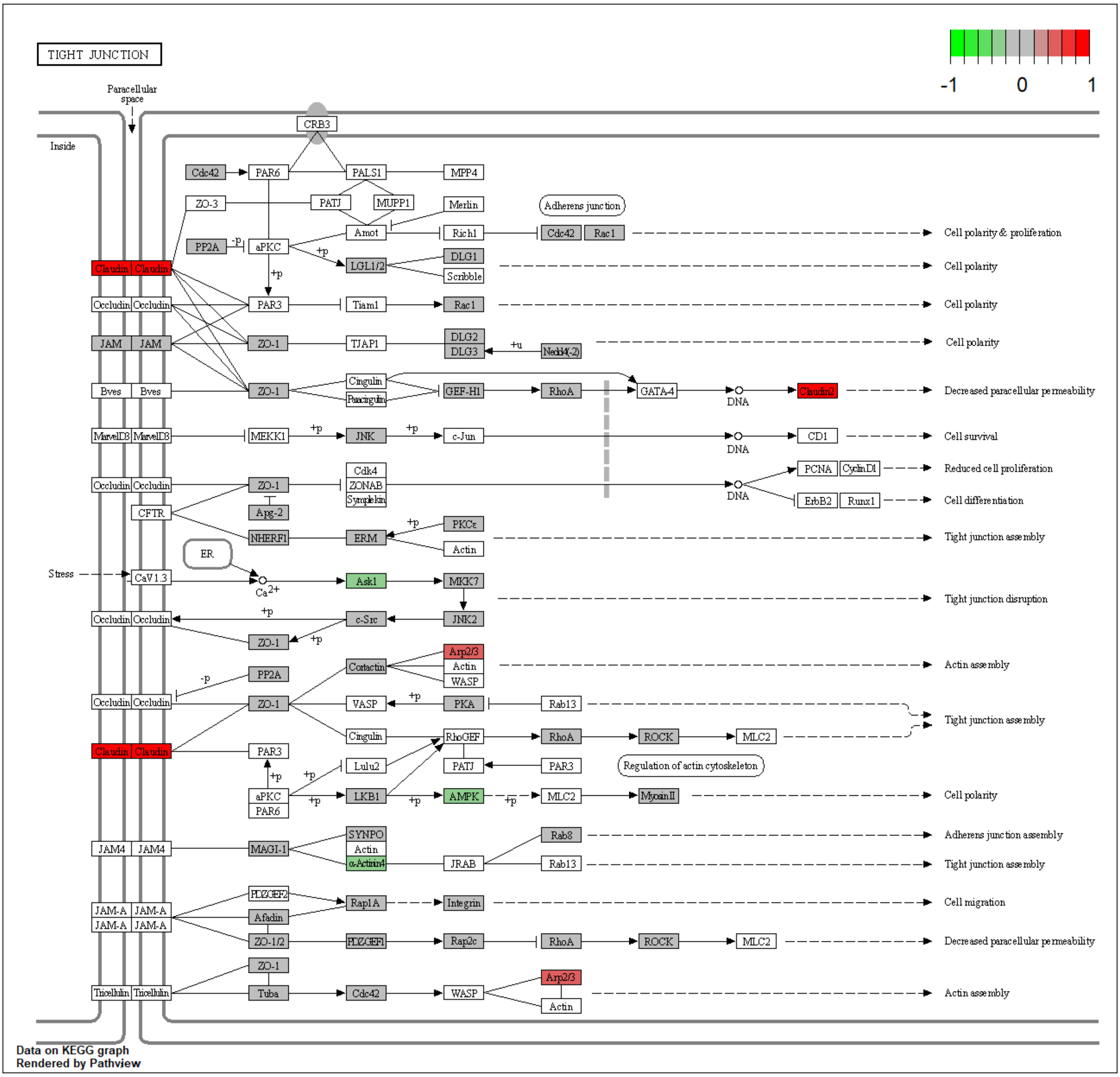

**Figure S26.** NMN treatment effects on the KEGG protein processing in endoplasmic reticulum pathway in mouse brain. The colour of the boxes represents the log2 fold change of the protein abundances, represented for HFD+NMN group versus HFD group comparison. Red: up-regulated; green: down-regulated; grey: no significant expression change.

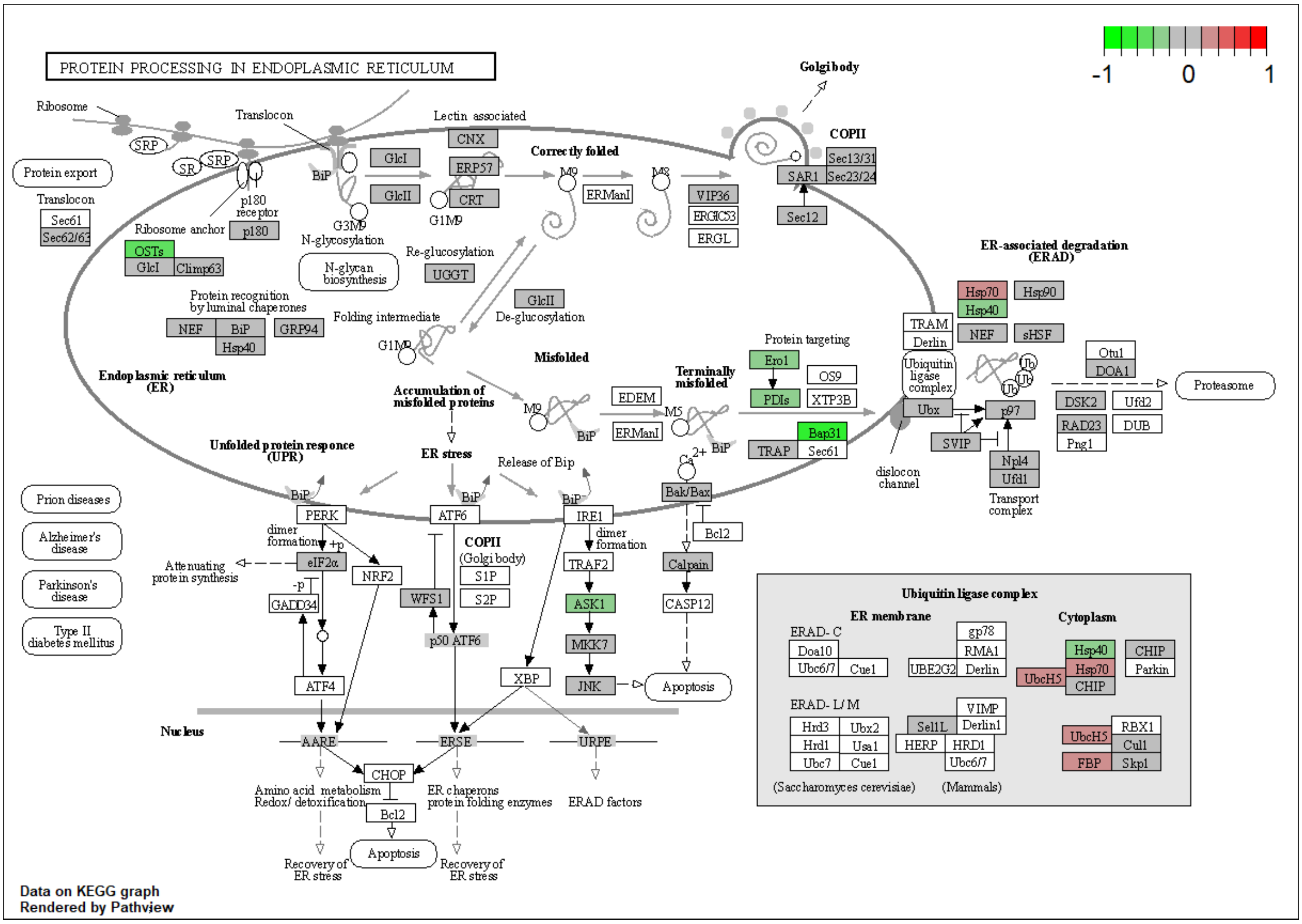

**Figure S27.** NMN treatment effects on the KEGG ribosome pathway in HepG2 cells. The colour of the boxes represents the log2 fold change of the protein abundances, represented simultaneously for HN and HH comparisons, H100N versus H0N left side, H100H versus H0H right side, in the corresponding box for each protein. Red: up-regulated; green: down-regulated; grey: no significant expression change.

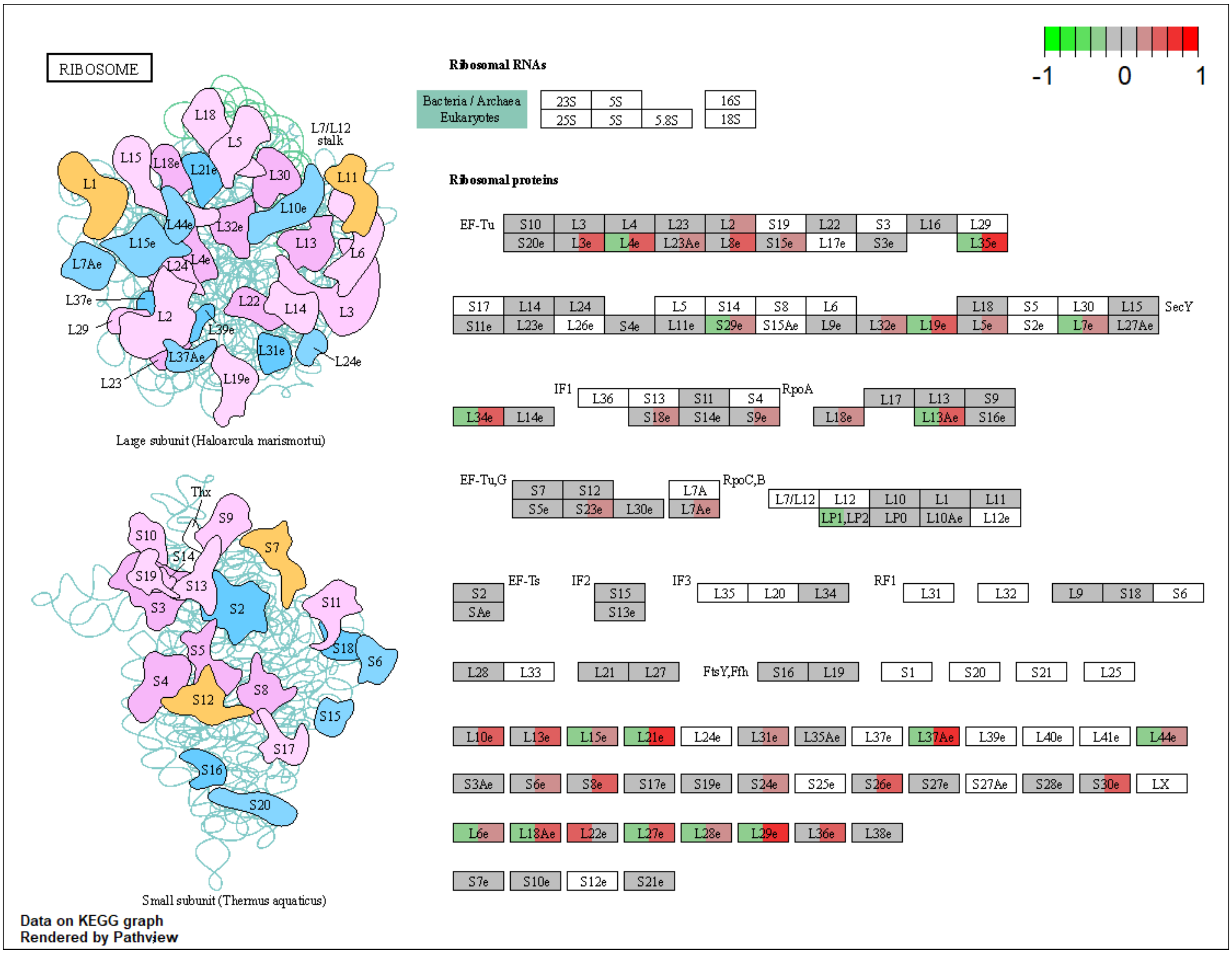

**Figure S28.** NMN treatment effects on the KEGG oxidative phosphorylation pathway in HepG2 cells. The colour of the boxes represents the log2 fold change of the protein abundances, represented simultaneously for HN and HH comparisons, H100N versus H0N left side, H100H versus H0H right side, in the corresponding box for each protein. Red: up-regulated; green: down-regulated; grey: no significant expression change.

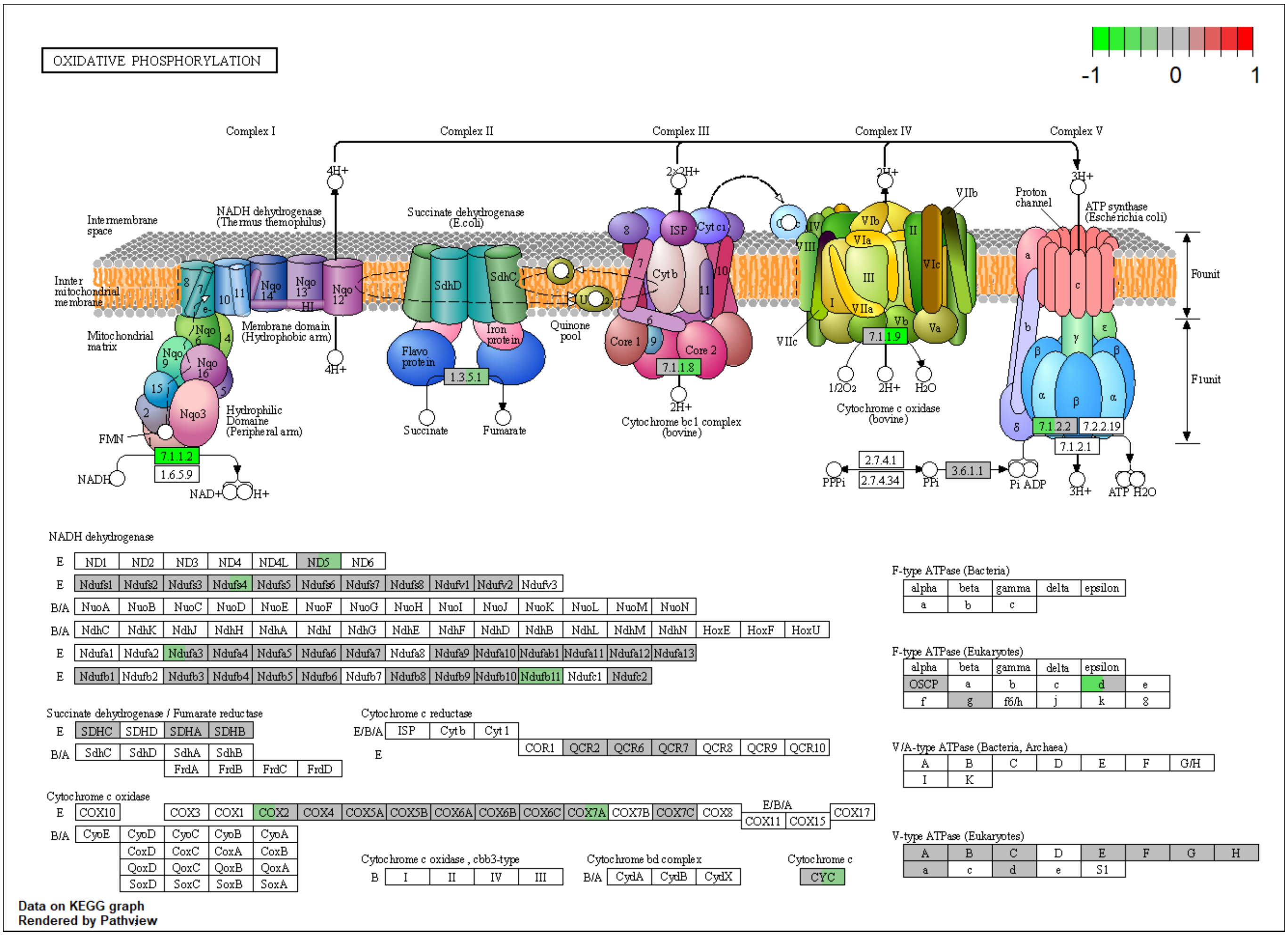

**Figure S29.** NMN treatment effects on the KEGG spliceosome pathway in HepG2 cells. The colour of the boxes represents the log2 fold change of the protein abundances, represented simultaneously for HN and HH comparisons, H100N versus H0N left side, H100H versus H0H right side, in the corresponding box for each protein. Red: up-regulated; green: down-regulated; grey: no significant expression change.

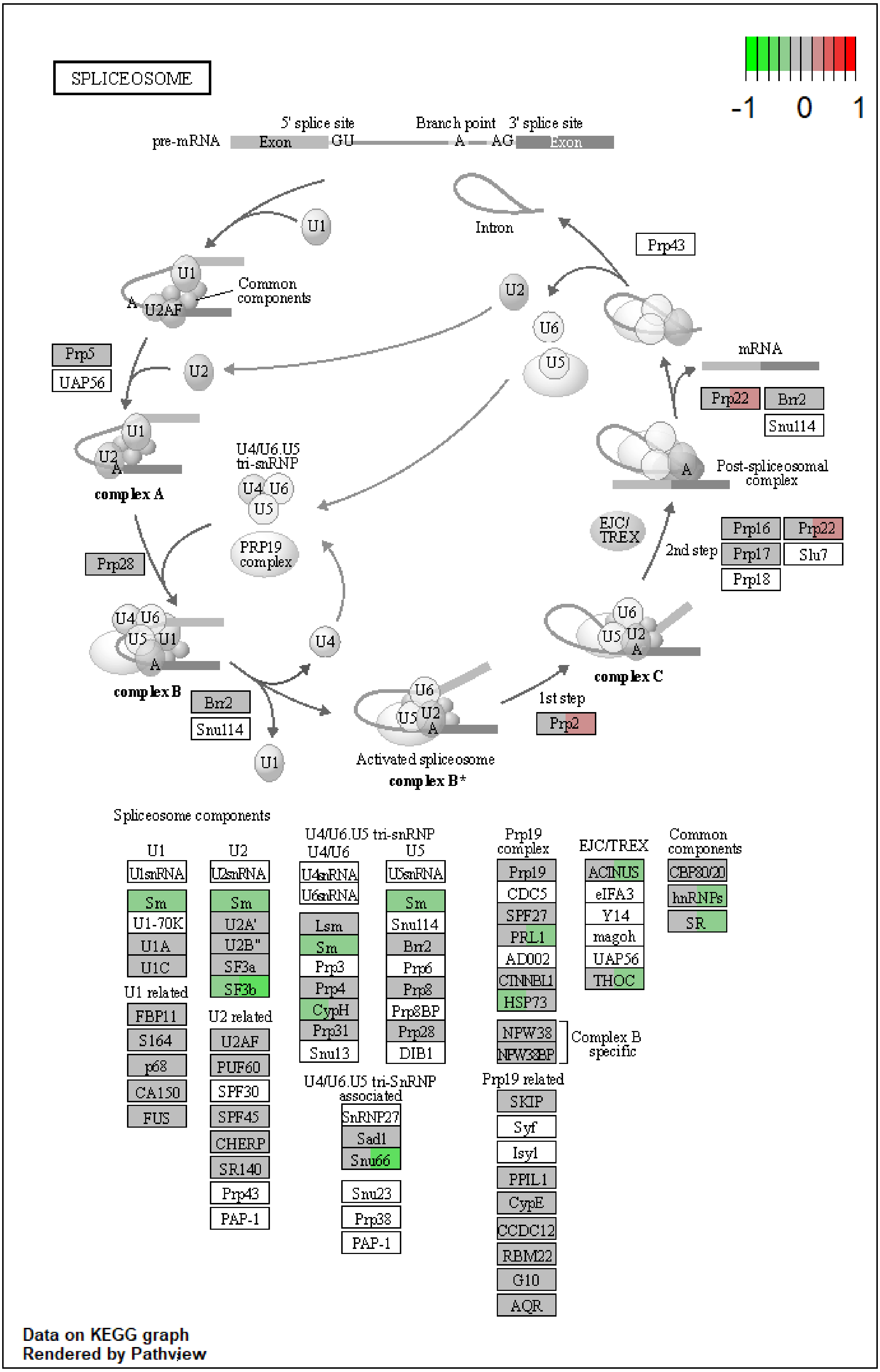

**Figure S30.** NMN treatment effects on the KEGG DNA replication pathway in HepG2 cells. The colour of the boxes represents the log2 fold change of the protein abundances, represented simultaneously for HN and HH comparisons, H100N versus H0N left side, H100H versus H0H right side, in the corresponding box for each protein. Red: up-regulated; green: down-regulated; grey: no significant expression change.

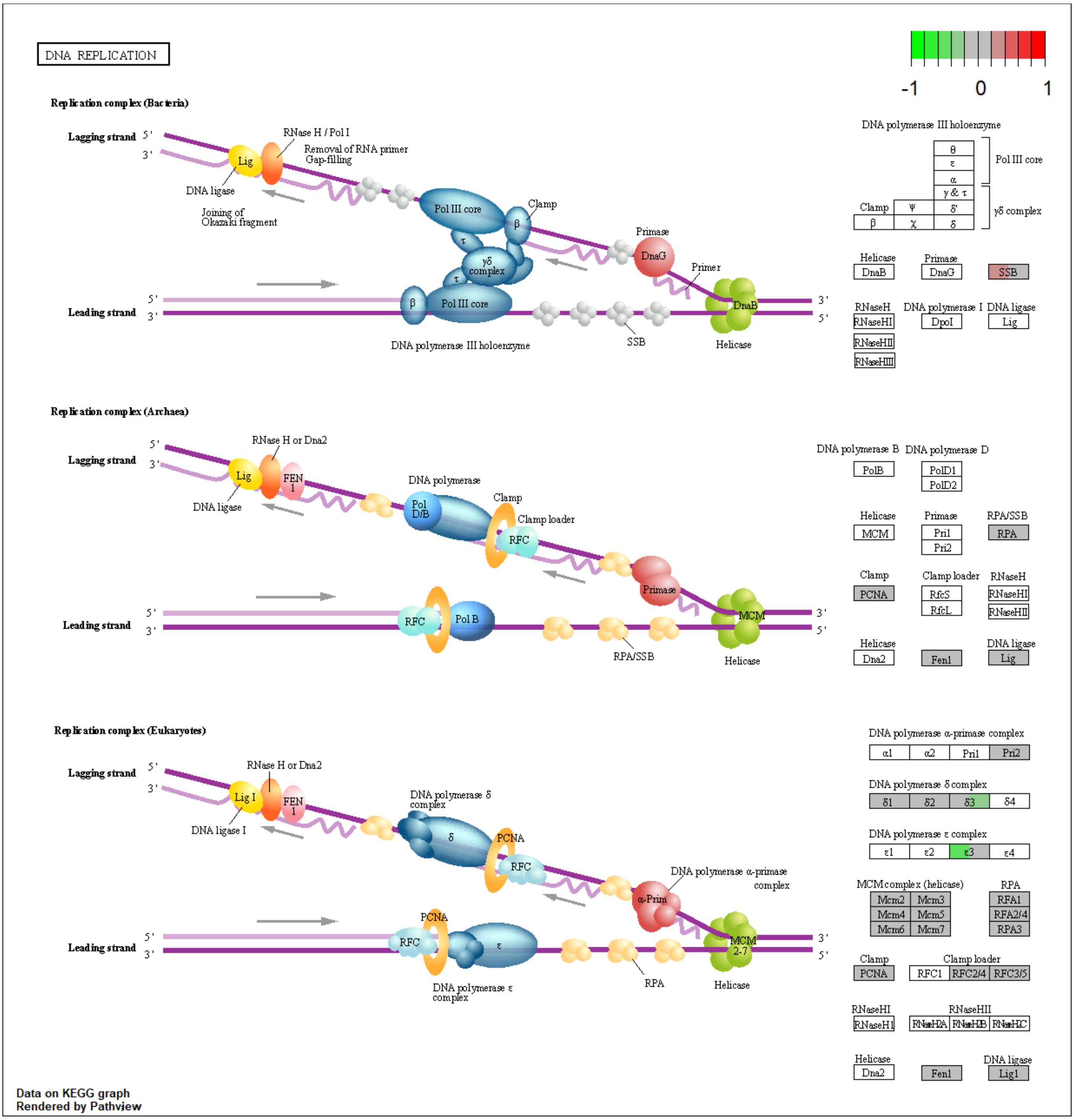

**Figure S31.** NMN treatment effects on the KEGG ribosome pathway in C2C12 derived myotubes. The colour of the boxes represents the log2 fold change of the protein abundances, represented simultaneously for HN and HH comparisons, H100N versus H0N left side, H100H versus H0H right side, in the corresponding box for each protein. Red: up-regulated; green: down-regulated; grey: no significant expression change.

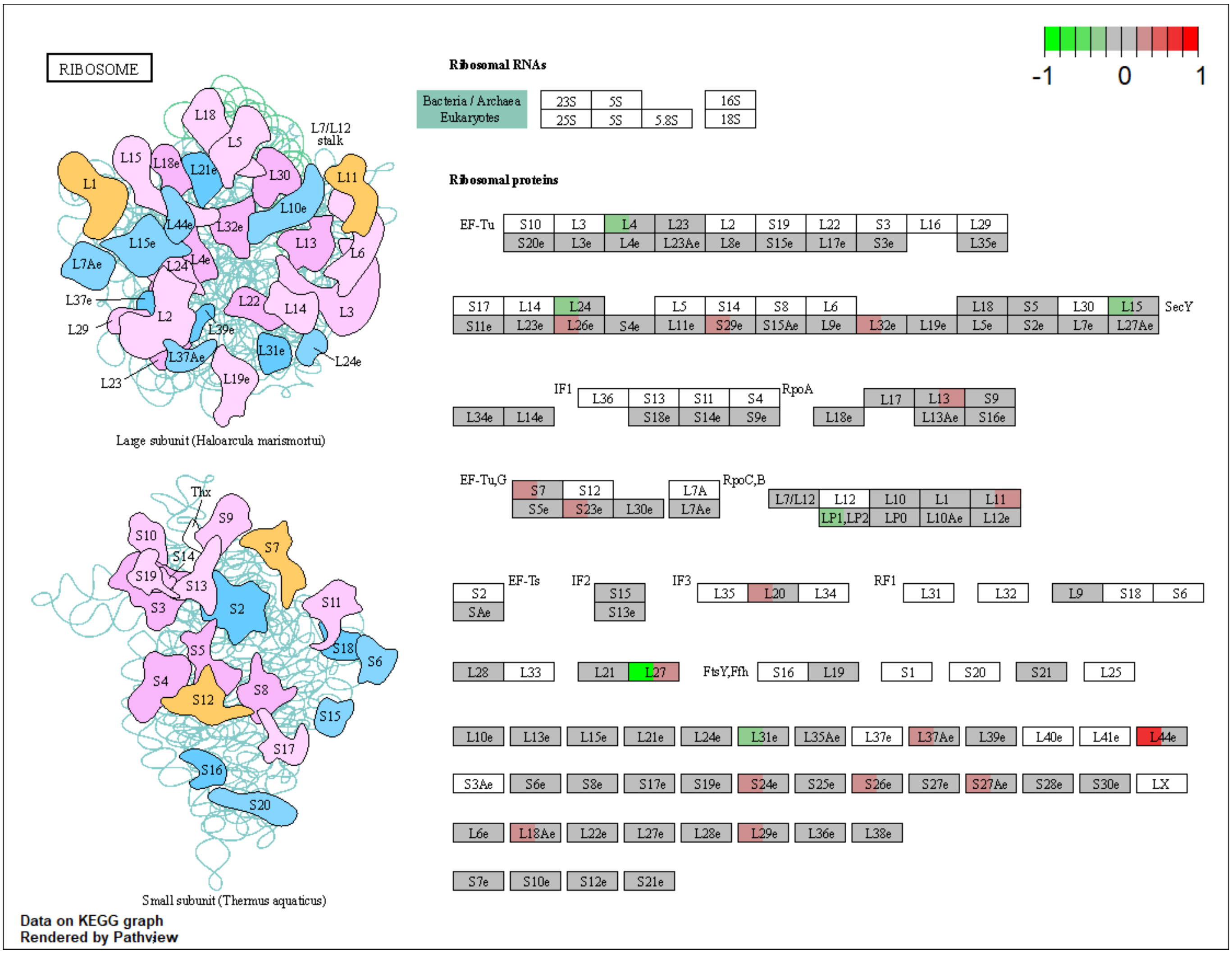

**Figure S32.** NMN treatment effects on the KEGG cell cycle pathway in C2C12 derived myotubes. The colour of the boxes represents the log2 fold change of the protein abundances, represented simultaneously for HN and HH comparisons, H100N versus H0N left side, H100H versus H0H right side, in the corresponding box for each protein. Red: up-regulated; green: down-regulated; grey: no significant expression change.

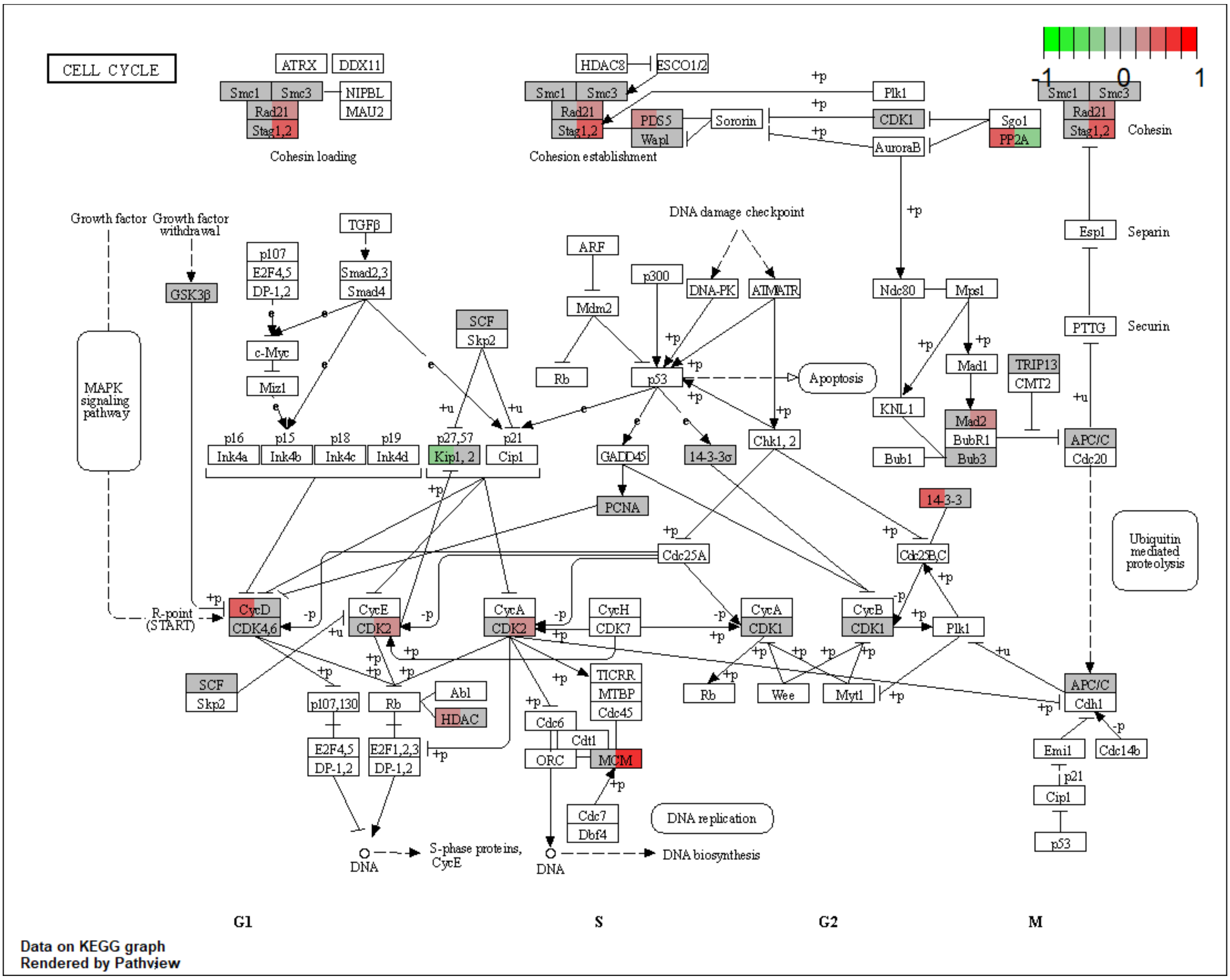

**Figure S33.** NMN treatment effects on the KEGG oxidative phosphorylation pathway in C2C12 derived myotubes. The colour of the boxes represents the log2 fold change of the protein abundances, represented simultaneously for HN and HH comparisons, H100N versus H0N left side, H100H versus H0H right side, in the corresponding box for each protein. Red: up-regulated; green: down-regulated; grey: no significant expression change.

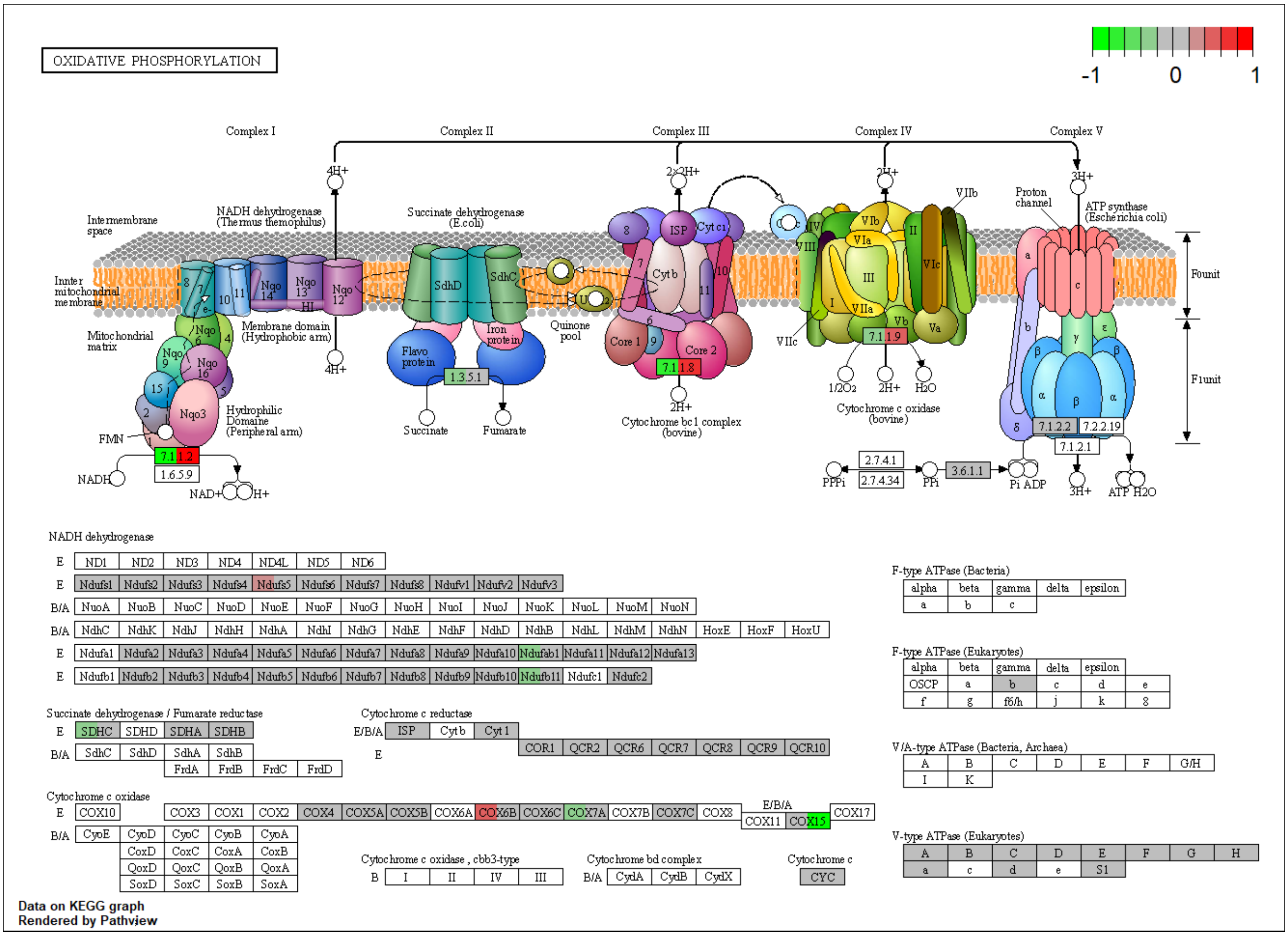

**Figure S34.** NMN treatment effects on the KEGG proteasome pathway in C2C12 derived myotubes. The colour of the boxes represents the log2 fold change of the protein abundances, represented simultaneously for HN and HH comparisons, H100N versus H0N left side, H100H versus H0H right side, in the corresponding box for each protein. Red: up-regulated; green: down-regulated; grey: no significant expression change.

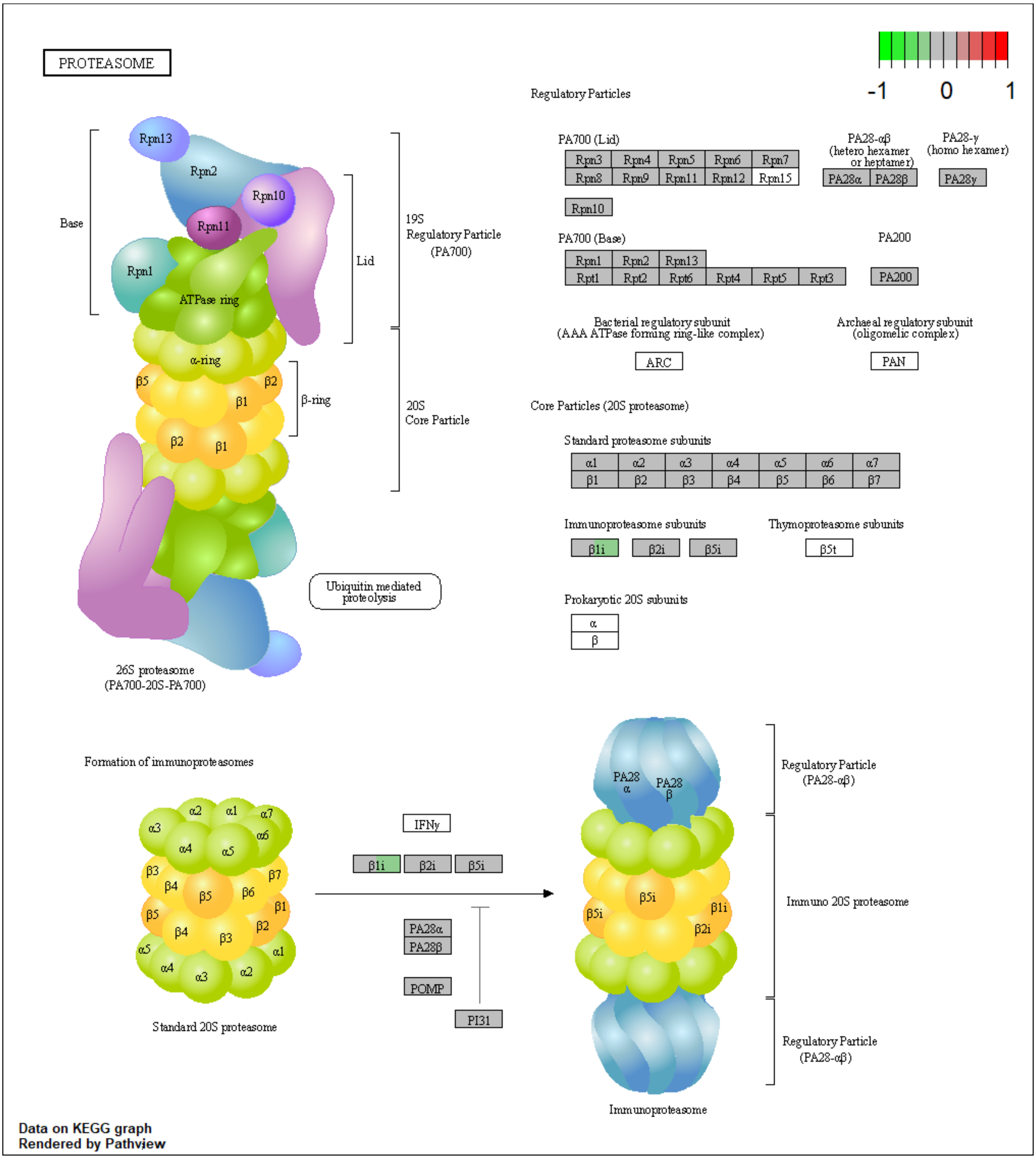

**Figure S35.** NMN treatment effects on the KEGG spliceosome pathway in C2C12 derived myotubes. The colour of the boxes represents the log2 fold change of the protein abundances, represented simultaneously for HN and HH comparisons, H100N versus H0N left side, H100H versus H0H right side, in the corresponding box for each protein. Red: up-regulated; green: down-regulated; grey: no significant expression change.

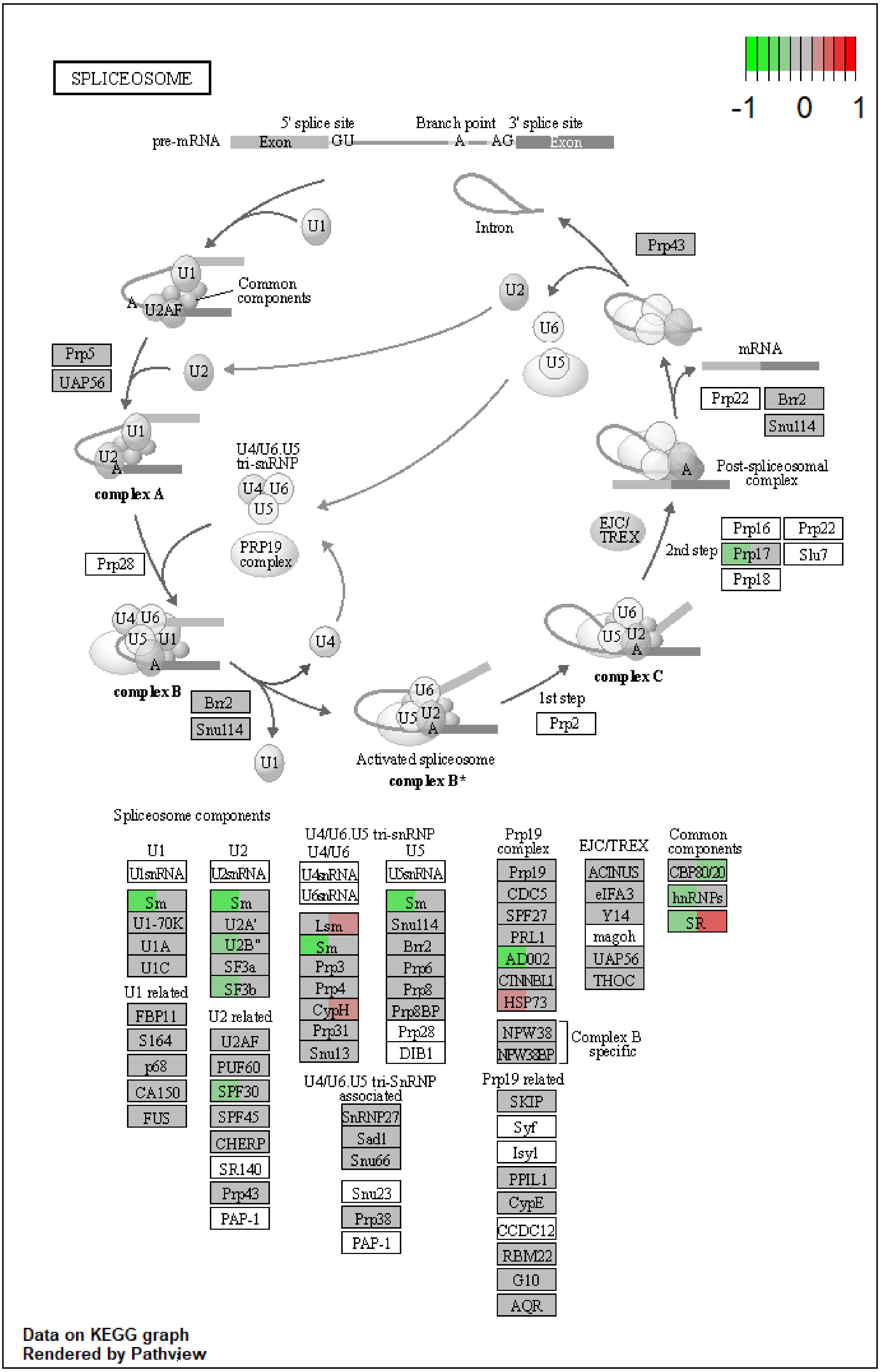
